# Aging limits stemness and tumorigenesis in the lung by reprogramming iron homeostasis

**DOI:** 10.1101/2024.06.23.600305

**Authors:** Xueqian Zhuang, Qing Wang, Simon Joost, Alexander Ferrena, David T. Humphreys, Zhuxuan Li, Melissa Blum, Klavdija Bastl, Selena Ding, Yuna Landais, Yingqian Zhan, Yang Zhao, Ronan Chaligne, Joo-Hyeon Lee, Sebastian E. Carrasco, Umeshkumar K. Bhanot, Richard P. Koche, Matthew J. Bott, Pekka Katajisto, Yadira M. Soto-Feliciano, Thomas Pisanic, Tiffany Thomas, Deyou Zheng, Emily S. Wong, Tuomas Tammela

## Abstract

Aging is associated with a decline in the number and fitness of adult stem cells^1–4^. Aging-associated loss of stemness is posited to suppress tumorigenesis^5,6^, but this hypothesis has not been tested *in vivo*. Here, using physiologically aged autochthonous genetically engineered mouse models and primary cells^7,8^, we demonstrate aging suppresses lung cancer initiation and progression by degrading stemness of the alveolar cell of origin. This phenotype is underpinned by aging-associated induction of the transcription factor NUPR1 and its downstream target lipocalin-2 in the cell of origin in mice and humans, leading to a functional iron insufficiency in the aged cells. Genetic inactivation of the NUPR1—lipocalin-2 axis or iron supplementation rescue stemness and promote tumorigenic potential of aged alveolar cells. Conversely, targeting the NUPR1— lipocalin-2 axis is detrimental to young alveolar cells via induction of ferroptosis. We find that aging-associated DNA hypomethylation at specific enhancer sites associates with elevated NUPR1 expression, which is recapitulated in young alveolar cells by inhibition of DNA methylation. We uncover that aging drives a functional iron insufficiency, which leads to loss of stemness and tumorigenesis, but promotes resistance to ferroptosis. These findings have significant implications for the therapeutic modulation of cellular iron homeostasis in regenerative medicine and in cancer prevention. Furthermore, our findings are consistent with a model whereby most human cancers initiate in young individuals, revealing a critical window for such cancer prevention efforts.

## Introduction

Cancer occurs at all ages, but incidence sharply rises in the sixth and seventh decades of life in humans^5,9–12^. This increase correlates with accumulation of somatic mutations in adult stem and progenitor cells^5,11^, which are the cell of origin for most cancer types^10^. However, the number and fitness of stem and progenitor cells declines with aging^1–4^, which has been suggested to counterbalance the accumulation of somatic mutations and inhibit tumorigenesis^5,6^. How age-associated loss of stem cell fitness impacts tumorigenesis in the context of potent cancer-causing mutations has not been investigated *in vivo*.

Telomere attrition, genomic instability, metabolic stress, cellular senescence, dysregulated cell-cell communication, and epigenetic alterations are “hallmarks of aging” that, among multiple detrimental effects in tissues, degrade stemness^13,14^, the potential of stem cells to self-renew and differentiate. Stereotypic changes in DNA methylation patterns, including hypomethylation at gene enhancers, represent the most ubiquitous molecular biomarker of aging across species and cell types^14–19^. Whether aging-associated changes in DNA methylation impact the tumorigenic potential of epithelial stem cells, the origin of most human cancers, has not been investigated *in vivo*.

Aging-associated changes in humans, including altered DNA methylation, are largely conserved in mice, rendering the mouse a prime model organism for studying mammalian aging^14^. Aging-associated phenotypes and molecular changes emerge in mice after 2 years of age, which corresponds to 65-70 years of age in humans^14^. Studies employing transplantation of syngeneic cancer cell lines into aged vs. young host mice have greatly advanced our understanding of changes in the aged host tumor microenvironment, which have pronounced effects on tumor growth and progression^20^. Although powerful, transplantation-based approaches more closely approximate advanced tumors. As such, our understanding of how aging impacts the early stages of tumorigenesis remains limited. In particular, how aging-associated cell-intrinsic changes in the cell of origin impact tumor initiation and progression have been little studied.

Lung adenocarcinoma (LUAD) is the most common lung cancer subtype and is responsible for ∼7% of all cancer mortality worldwide^9^. The median age for LUAD diagnosis is 71 years and aging correlates with an increased incidence of LUAD in both smokers and non-smokers^21^, but this incidence begins to decline after 80-85 years of age^9^. Understanding how aging impacts LUAD biology is of critical clinical importance, not least because young and very elderly patients are inadequately represented in clinical trials due to the rarity of cases^9,22^ and exclusion due to co-morbidities and frailty^7,8^, respectively. LUAD predominantly arises from alveolar type 2 (AT2) cells^7,8,23^, which function as the facultative stem cells of lung alveoli by self-renewing and replacing alveolar type 1 (AT1) cells under physiologic conditions and upon injury^24,25^.

Important insights into the biology and therapy of LUAD have been gleaned from genetically engineered mouse models (GEMMs). In the most widely used model, viral expression of *Cre* recombinase in AT2 cells leads to somatic activation of oncogenic **<u>K</u>**RAS-G12D and deletion of the **<u>p</u>**53 tumor suppressor (***<u>KP</u>*** model)^7,8^. *KP* tumors develop *de novo* from a single AT2 cell and recapitulate key molecular and histopathological features of the full spectrum of LUAD evolution in humans^7,8,23^. Notably, *KP* tumors can be engineered to express fluorescent reporters or CRISPR/Cas9, enabling isolation of cancer cells and somatic perturbation of genes of interest in incipient tumors, respectively^24,25^. Here, we leverage the *KP* model to investigate LUAD development by initiating potent cancer-causing mutations in a defined physiologically aged cell of origin *in vivo*.

## Results

### Aged AT2 cells exhibit reduced cell-intrinsic potential for regeneration and tumorigenesis

To investigate the impact of aging on lung cancer development, we induced LUAD tumors in young (12-16-week-old) and aged (104-130-week-old) *KP* mice using adenoviral vectors encoding Cre recombinase placed under the control of an AT2-specific surfactant protein C (SPC) promoter^26^ (AdSPC-Cre, **Fig. 1a**). Surprisingly, aged mice exhibited a 37% increase in median survival when compared to young mice (**Extended Data Fig. 1a**; **Extended Data Table 1**). To investigate the reason underlying this increase in survival, we examined tumor burden in lungs of *KP* mice at distinct stages of LUAD progression: atypical adenomatous hyperplasia (AAH, 4 weeks), adenoma (8 weeks), adenoma-to-adenocarcinoma transition (12 weeks), and fully formed adenocarcinoma (17 weeks) (**Fig. 1a**). We observed a dramatic decrease in the number of tumors in aged mice at all time points when compared to young counterparts (**Fig. 1b, c**; **Extended Data Fig. 1b**), suggesting that aging suppresses tumor initiation. In support, aged *KP* mice harboring a *Rosa26^LSL-tdTomato^*Cre recombination reporter (*KPT* mice) showed a significant increase in the proportion of single tdTomato^+^ cells vs. multicellular neoplastic nodules compared to the young at 2 weeks post-tumor initiation (**Fig. 1d**; **Extended Data Fig. 1c**), indicating reduced capacity of aged AT2 cells to initiate tumorigenesis. Importantly, the absolute number of AT2 cells transduced by the AdSPC-Cre vector was not different between aged and young Cre reporter mice (**Extended Data Fig. 1d**). We next examined *KP* AT2 cells at different stages of the mouse life span *in vivo* and found the tumorigenic potential of AT2 cells declined dramatically as a function of age (**Extended Data Fig. 1e**).

**Figure 1.**
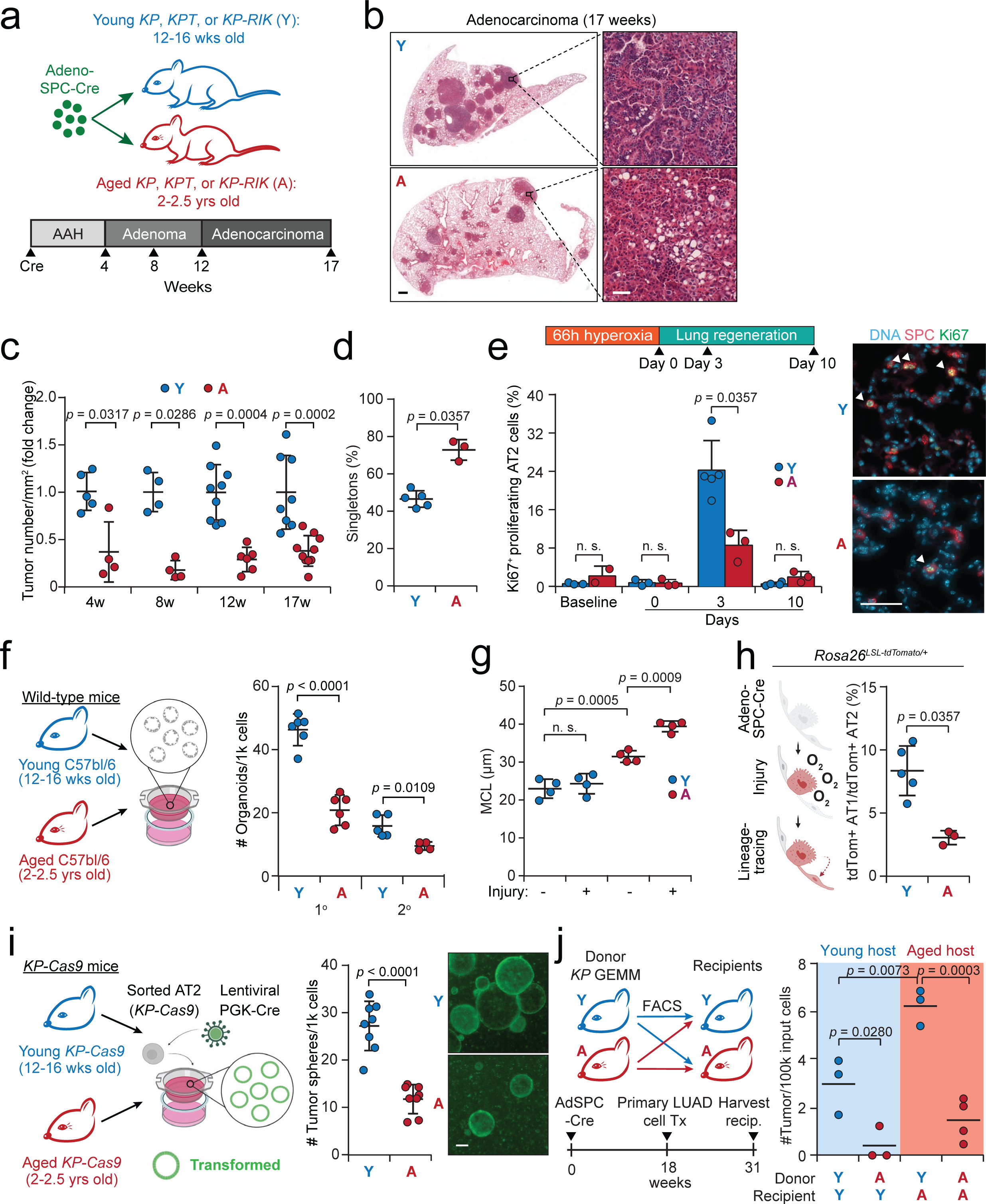
Aged AT2 cells exhibit reduced cell-intrinsic potential for regeneration and tumorigenesis. (**a**) Experimental scheme. *KP: Kras^lox-stop-lox(LSL)-G12D/+^;Trp53^fl/fl,^, KPT: Kras^LSL-^ _G12D/+;Trp53_fl/fl_Rosa26_LSL-tdTomato/+_, KP-RIK: Kras_LSL-G12D/+_;Trp53_fl/fl_;Rosa26_LSL-rtTA-IRES-mKate2/+*. SPC: surfactant protein-C gene promoter. AAH, Atypical adenomatous hyperplasia. (**b**) Representative images of hematoxylin-eosin staining of tumor-bearing lungs from aged and young *KP* mice 17 weeks post-tumor initiation. Scale bars: 1 mm (left panel) and 50 µm (right panel). (**c**) Fold change of tumor burden in aged vs. young *KP* mice at 4, 8, 12, and 17 weeks post-tumor initiation. *N* = 5, 4, 9, and 8 young mice and 4, 4, 6, and 10 aged mice at 4, 8, 12 and 17 weeks, respectively. (**d**) Percentage of single tdTomato^+^ cells (singletons) per total number of tdTomato^+^ lesions at 2 weeks post-tumor initiation using adeno-SPC-Cre in aged vs. young *KPT* mice. *N* = 5 young and 3 aged mice. (**e**) Proliferation of AT2 cells at homeostasis (baseline) and during alveolar regeneration in response to hyperoxia injury at days 0, 3, and 10 following 66-hour exposure to hyperoxia. Representative images of proliferating (Ki67^+^, green) AT2 cells (SPC^+^, red) are shown on the right. *N* = 3, 3, 5, and 4 young mice and 2, 3, 3 and 3 aged mice at baseline, at days 0, 3, and 10 post-hyperoxia, respectively. Scale bar: 20 µm. (**f**) Alveolar organoid formation assay (left). Primary and secondary alveolar organoid formation derived from aged vs. young primary AT2 cells (right). *N* = 6 biological replicates for primary organoid passage and *n* = 5 and 4 biological replicates for secondary organoid passage. (**g**) Airspace size of aged and young mice before and after hyperoxia-induced lung injury, measured by mean cord length (MCL). *N* = 4 mice per group. (**h**) Lineage-tracing of aged and young tdTomato-labeled AT2 cells following alveolar injury. Shown is the percentage of tdTomato^+^ AT1 cells (defined by co-expression of the AT1 marker podoplanin and tdTomato) per the number of tdTomato^+^ AT2 cells (defined by co-expression of the AT2 marker surfactant protein-C and tdTomato). *N* = 5 young and 3 aged mice. (**i**) *Ex vivo* transformation assay employing primary AT2 cells isolated from *KP-Cas9* (*Kras^LSL-G12D/+^;Trp53^fl/fl^; Rosa26^LSL-Cas9-2a-eGFP/+^*) mice. Shown is the number of GFP^+^ tumor spheres transformed by lentiviral delivery of Cre recombinase (*n* = 8 biological replicates per condition). Representative images of organoids are shown on the right. PGK: ubiquitously active human phosphoglycerate kinase-1 promoter. Scale bar: 200 µm. (**j**) Heterochronic transplantation of *KP* tumor cells. Shown on the left is the experimental scheme and on the right is the quantification of tumor burden (*n* = 3-4 mice per group). Mean with SD is shown in (**e**). Y: young, A: aged. *Mann-Whitney* test was used in (**c-e**) and (**h**). Student’s *t-*test was used in (**d**), (**f**) and (**i**). One-way ANOVA was used in (**g**) and (**j**).

To test whether these findings are relevant beyond the *KP* genotype we initiated lung tumors in aged vs. young wild-type mice using CRISPR-mediated engineering of *Eml4*-*Alk* fusions^27^ (**Extended Data Fig. 2a**), which represents another clinically relevant genetic LUAD subtype. Aged mice displayed a robust reduction in the number of tumors and overall tumor burden when compared to young mice (**Extended Data Fig. 2b-d**), indicating that the loss of tumorigenic potential in the lung epithelium is independent of driver oncogene or p53 status. Importantly, aged and young AT2 cells showed a similar rate of *Eml4-Alk* fusion events *in vivo* (**Extended Data Fig. 2e,f**), indicating the differences observed are not due to technical reasons.

Tumor initiation potential is tightly linked to the number and capacity for <u>self-renewal and differentiation</u> (stemness) of the cell of origin within tissues^10^. We detected a 47% reduction in the density of AT2 cells in lungs of aged wild-type mice compared to young lungs (**Extended Data Fig. 3a**). Furthermore, aging led to a progressive reduction in the proportion of AT2 cells within the total lung epithelial cell pool (**Extended Data Fig. 3b-d**), suggesting that AT2 cells decline with aging more than other lung epithelial cells. To interrogate the <u>self-renewal capacity</u> of AT2 cells *in vivo*, we stimulated alveolar regeneration in aged and young wild-type mice by hyperoxia injury^28,29^. Compared to young mice, the aged mice displayed a 69% reduction in the proportion of proliferating (Ki67^+^) AT2 cells at three days following injury, the peak proliferative phase of alveolar regeneration (**Fig. 1e**). To study the cell-intrinsic self-renewal capacity of AT2 cells within a controlled and consistent *in vitro* environment, we subjected isolated primary AT2 cells to a 3D alveolar organoid assay^30^. We observed a 54% reduction in the number of organoids formed by AT2 cells isolated from aged mice compared to the young. The reduced potential for organoid growth persisted in secondary passage (**Fig. 1f; Extended Data Fig. 3e**), further implicating cell-intrinsic changes that lead to decline of self-renewal capacity in aged AT2 cells. Of note, senescence-associated beta-galactosidase activity in the AT2 cells was nearly absent in both aged and young mice, although an increase in non-epithelial senescent cells was observed in aged lungs (**Extended Data Fig. 3f**).

To interrogate their <u>differentiation capacity</u> *in vivo*, we tested the ability of aged AT2 cells to restore alveolar integrity, a process in lung regeneration dependent on AT2 cells^31^, by assessing the alveolar airspaces after injury using mean chord length (MCL) mesurement^32^. Interestingly, the airspaces of aged alveoli were already enlarged in the absence of injury, consistent with a cumulative loss of AT2 cell regenerative capacity with age. The aged alveoli enlarged further and remained significantly more enlarged at 4 weeks following injury compared to baseline, whereas the airspaces of young alveoli were restored back to baseline (**Fig. 1g; Extended Data Fig. 3g**). We next lineage-traced AT2 cells in aged vs. young *Rosa26^LSL-tdTomato^*mice by intra-tracheal delivery of AdSPC-Cre, followed by hyperoxia injury, and found a dramatic decline in the ability of aged AT2 cells to differentiate into AT1 cells when compared to the young (**Fig. 1h**). Taken together, these results demonstrate that aging suppresses stemness of AT2 cells.

To directly interrogate transformation capacity, we isolated primary AT2 cells from aged and young *KP;Rosa26^Cas^*^9^*^-GFP/+^* (hereafter *KP-Cas9*) mice^33^ and induced transformation by lentiviral delivery of Cre recombinase in an *ex vivo* tumor sphere formation assay^34^. Similar to the *in vivo* experiments, the efficiency of transformation was significantly impaired in aged AT2 cells (**Fig. 1i**), even when adjusting for the baseline difference in alveolar organoid formation between aged and young AT2 cells (**Extended Data Fig. 3h**). This difference was not due to susceptibility to lentiviral transduction, which was similar in both aged and young AT2 cells *ex vivo* and *in vivo* (**Extended Data Fig. 3i,j**). Interestingly, the 50-70% reduction in AT2 cell self-renewal capacity (**Fig. 1e,f**) and differentiation capacity (**Fig. 1h**) closely mirrored the overall ∼60% reduction in tumorigenic potential of the aged AT2 cells (**Fig. 1d**), suggesting that loss of AT2 stemness may underlie reduced tumor initiation in the aged lung. Given the aged AT2 cells show a defect in growth and transformation potential *ex vivo*, when removed from the aged tissue microenvironment, indicates aging suppresses AT2 tumorigenesis via cell-intrinsic mechanisms.

To directly test the contribution of the lung microenvironment, we isolated primary autochthonous *KP* LUAD cells from aged vs. young donor mice and performed heterochronic transplantation into lungs of aged vs. young syngeneic recipients. *KP* cells isolated from young donors formed tumors significantly more efficiently than cells isolated from aged donors, irrespective of the age of the recipient. Interestingly, *KP* cells from young donors formed more tumors in aged recipients (**Fig. 1j**). These results suggest the aged lung microenvironment may be more permissive for LUAD tumorigenesis. Yet, despite this, aged mice develop fewer tumors, which is explained by intrinsic loss of AT2 cell transformation capacity.

### Lung tumors in aged mice show delayed progression

In addition to reduced initiation frequency, tumors that formed in aged *KP* mice were reduced in size by 40-60% at 4, 8, and 17 weeks of tumor development when compared to young counterparts; however, no difference was observed at 12 weeks (**Extended Data Fig. 4a**). The proliferative index of *KP* tumors was significantly lower in aged mice at all time points (**Extended Data Fig. 4b**). Conversely, we did not detect a difference in the number of senescent cells (**Extended Data Fig. 4c-e**) or cleaved caspase-3 positive apoptotic cells (**Extended Data Fig. 4f**) in the aged vs. young tumors. These results indicate that decreased proliferation rather than increased senescence or apoptosis explains the smaller tumor size in aged mice.

Besides reduced proliferation, the aged *KP* lungs showed a higher proportion of grade (G) 1 AAH lesions at 12 weeks and G2 adenomas at 17 weeks post-tumor initiation when compared to young *KP* tumors (**Fig. 1c; Extended Data Fig. 4g,h**). Conversely, young mice showed a higher fraction of G4 advanced adenocarcinomas than aged mice at 17 weeks (**Extended Data Fig 4h**). These findings indicate delayed histological progression of aged *KP* tumors. To investigate the molecular evolution of LUAD tumors induced in aged vs. young mice, we analyzed *KP* LUAD cells at 4, 12, and 17 weeks following tumor initiation using single-cell mRNA sequencing (scRNA-seq). AT2 cells from wild-type aged vs. young mice were also collected for analysis. Unsupervised clustering of scRNA-seq data revealed six clusters with distinct expression patterns, five of which corresponded to cancer cell states that we had previously identified during LUAD evolution^23^ and the wild-type AT2 cells formed their own cluster (**Extended Data Fig. 5a**). While both the aged and young tumors progressed along a similar trajectory, the aged tumors displayed a significant delay in progression (**Extended Data Fig. 5b-d**). For example, at four weeks, a higher fraction of young *KP* cells had progressed to a high-plasticity cell state–an indicator of early progression^23^– than in the aged tumors (**Extended Data Fig. 5b,c**). At 17 weeks more young *KP* cells were found in the endoderm-like state (**Extended Data Fig. 5d**)–a transition that occurs late in progression^23,35^–when compared to the aged *KP* cells. We validated our scRNA-seq findings by immunostaining for markers of the high-plasticity (integrin α2 and keratin-8)^23,36^ and endoderm-like (HNF4α)^35^ states at the early and late progression time points, respectively (**Extended Data Fig. 5b-d**).

We next modeled tumor evolution using our *ex vivo* AT2 cell transformation assay (**Fig. 1i**) and analyzed resulting tumor spheres for bulk RNA sequencing analysis over eight passages (∼3 months); freshly isolated primary AT2 cells and primary AT2 organoid cultures from aged and young wild-type mice were analyzed as baseline comparators (**Fig. 1f,i; Extended Data Fig. 5e**). Although the transcriptomic features of the *ex vivo KP* tumor spheres are distinct from the LUAD tumors *in vivo* (**Extended Data Table 2**), tumor spheres initiated from aged AT2 cells displayed a significant delay in their progression along an unsupervised diffusion pseudotime trajectory (**Extended Data Fig. 5f,g**). This delay was exemplified by a slower downregulation of alveolar markers in tumor spheres derived from aged vs. young AT2 cells (**Extended Data Fig. 5h**). These findings suggest that AT2 cell-intrinsic changes induced by aging delay the progression of LUAD tumors.

### Induction of Nupr1 suppresses tumorigenic potential in the aged lung

To uncover molecular mechanisms underpinning loss of AT2 transformation potential, we performed differential gene expression analysis on our scRNA-seq data comprising aged vs. young AT2 cells and LUAD cells. We detected a strong age-dependent correlation between differentially expressed genes (DEGs) in both the AT2 cells and the LUAD cells (**Fig. 2a**). Based on a two-part generalized statistical model (see **Methods**), 170 genes—8.2% of the DEGs in aged vs. young AT2 cells—showed a consistent change in expression in the AT2 cells as well as in all five transcriptional cell states that molecularly define LUAD progression (**Fig. 2b; Extended Data Table 3**). Our results suggest that a subset of the aging-associated changes in gene expression are shared between the AT2 cell of origin and the LUAD tumors that arise from them, and that these changes persist in distinct cancer cell states during tumor progression.

**Figure 2.**
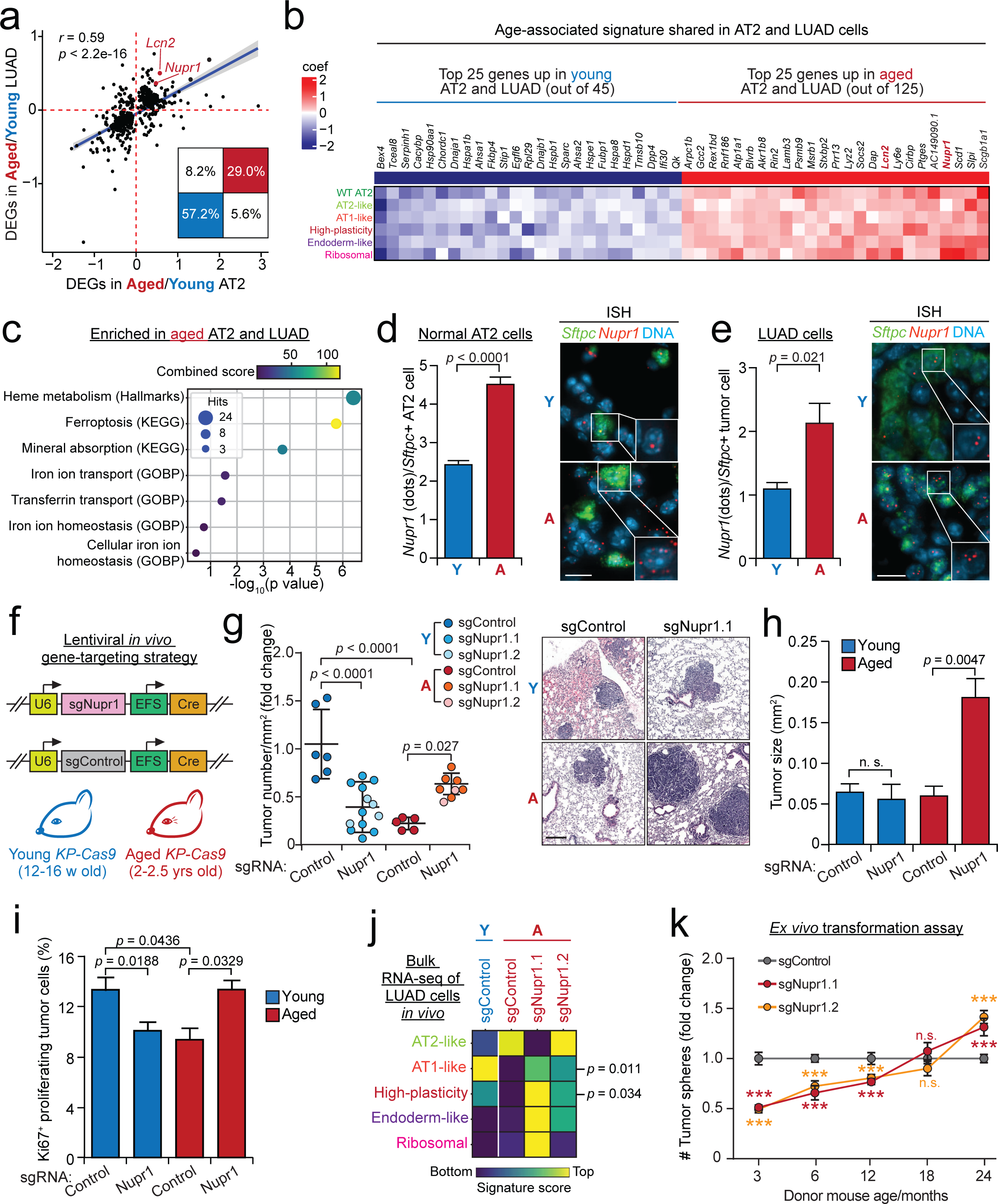
Induction of *Nupr1* suppresses tumorigenic potential in the aged lung. (**a**) Pearson correlation between the model-based analysis of single-cell transcriptomics (MAST) differentially expressed genes (DEGs) coefficient^74^ aged vs. young in normal AT2 cells (*x*-axis) and all LUAD cells (*y*-axis). The inset shows the percentage of genes in each quadrant, which is significantly different from random (binomial test *p* < 2.2^-16^). The regression line (blue) and its confidence interval (light grey) were plotted using a linear model. (**b**) Heatmap displaying the MAST DEG coefficient of the top 25 most consistently upregulated and downregulated genes across normal and tumor cell types. The top 25 upregulated and top 25 downregulated genes with aging were selected based on (i) significance in normal AT2 with FDR < 0.1; (ii) similar gene expression change trend across AT2 cells and all LUAD cell states; (iii) the sum of absolute DEG coefficient across AT2 cells and all LUAD cell states. The color bar indicates the MAST coefficient value. The genes are ordered from most downregulated (dark blue) to the most upregulated (dark red) in aged across the *x*-axis. The uniform manifold approximation and projection (UMAP) plot of the single-cell transcriptomics showing wildtype AT2 cells (cluster 0) and five LUAD cell states (AT2-like, AT1-like, high-plasticity cell state, HPCS, endoderm-like and ribosome-high, cluster 1-5) that molecularly define LUAD progression is shown in **Extended Data Fig. 5a**. Note that the expression of the DEGs in the wildtype AT2 is shown at the top row and the five LUAD cell clusters follow. The DEGs were ranked based on a two-part generalized linear statistical model where they were identified in all tumor cells with batch, sex, cell type, and cellular detection rate added as covariates. Subsequently, the aging-related DEGs in the normal AT2 cells and in all five LUAD cell states were identified separately with batch, sex, and cellular detection rate included in the model as covariates. (**c**) Dot plot showing enrichment of iron metabolism-related gene sets in the aged signature defined in (**b**) and in **Extended Data Table 3**. Combined score = log(***p***)·***z***, where ***p*** is the p-value computed using the Fisher exact test, and *z* is the z-score computed by assessing the deviation from the expected rank. (**d-e**) Quantification of *Nupr1* mRNA level (red dots) by *in situ* hybridization in AT2 cells (**d**) (*n* = 1126 and 425 AT2 cells from young and aged animals, respectively) and LUAD cells at 12 weeks post-tumor initiation (**e**) (*n* = 32 and 12 tumors from young and aged tumor-bearing mice at 12 weeks post-tumor initiation, respectively). Surfactant protein-C (*Sftpc*) mRNA (green) was used to mark AT2 and cancer cells. Scale bar: 20 µm. (**f**) Schematic summary of lentiviral *Nupr1* gene-targeting strategy in the context of LUAD tumorigenesis *in vivo*. (**g**) Quantification of tumor burden in lungs of aged and young *KP-Cas9* mice with non-targeting control or *Nupr1*-targeting sgRNAs (from left to right, *n* = 6, 13, 5, and 8 mice in each of group). Representative images of pulmonary tumors are shown on the right (HE staining). Scale bar: 200 µm. (**h-i**) Quantification of tumor size (**h**) (*n* = 87, 40, 146, and 121 tumors for young-sgControl, young-sg*Nupr1*, aged-sgControl, and aged-sg*Nupr1*, respectively) and proportion of proliferating (Ki67^+^) tumor cells per total tumor cells (**i**) (*n* = 71, 133, 40, and 93 tumors for young-sgControl, young-sg*Nupr1*, aged-sgControl, and aged-sg*Nupr1*) in the experiment described in (**f**). (**j**) Bulk RNA-seq analysis of sorted *KP-Cas9* cells from LUAD tumors harboring the indicated sgRNAs at 10 weeks post-tumor initiation. Enrichment of *KP-LUAD* cell state signatures (Fig. 2b) that define progression is shown. (**k**) *Ex vivo* transformation of AT2 cells derived from 3-24-month-old *KP-Cas9* mice transduced with lentiviral vectors delivering Cre recombinase plus sgRNAs targeting *Nupr1* or a non-targeting control (**f**). Number of tumor spheres is normalized to sgControl of each time point. *N* = 4 technical repeats. Numerical *p* value was shown in **Extended Data Table 19**. Mean with SEM is shown in (**d**), (**e**), (**h**), and (**i**). Mean with SD is shown in (**k**). Y: young; A: aged. Student’s *t* test was used in (**d**), (**e**), and (**k**) and one-way ANOVA was used in (**g-i**).

We hypothesized that the components of the shared gene programs changed by aging in both AT2 and LUAD cells may underlie the reduction in tumor initiation and progression in aged mice. Gene set enrichment analysis revealed a consistent induction of genes involved in iron homeostasis (**Fig. 2c; Extended Data Fig. 6a; Extended Data Table 4**). The top gene induced by aging within these pathways was nuclear protein, transcriptional regulator-1 (*Nupr1*) (**Fig. 2b; Extended Data Figure 6b**), a stress-induced transcriptional co-activator that was recently shown to control iron homeostasis^37,38^. In addition to iron metabolism, NUPR1 participates in the regulation of a range of other cellular processes, including DNA repair, ER stress, oxidative stress response, cell cycle, autophagy, apoptosis, and chromatin remodeling^38,39^. However, changes in these other pathways in our dataset were less pronounced or consistent across the distinct stages of progression (**Extended Data Table 4**). NUPR1 has been shown to be induced in a variety of cancers and targeting NUPR1 has anti-cancer effects in some contexts^38,39^.

We confirmed induction of *Nupr1* mRNA *in situ* in AT2 cells in aged wild-type mice and in the aged LUAD tumors (**Fig. 2d,e**). We next functionally interrogated *Nupr1* in lung tumor initiation *in vivo* by lentiviral co-delivery of single guide RNAs (sgRNAs) and Cre recombinase into lungs of aged vs. young *KP-Cas9* mice. In this system *KP* tumor initiation is coupled to activation of *Cas9* and expression of an sgRNA targeting *Nupr1* or a non-targeting control sgRNA (**Fig. 2f**)^25^. Consistent with published literature identifying NUPR1 as an anti-cancer target^38,39^, inactivation of *Nupr1* suppressed tumorigenesis in young mice at 12 weeks post-tumor initiation. However, remarkably, *Nupr1* gene targeting in aged AT2 cells promoted tumor initiation *in vivo* and *ex vivo* (**Fig. 2g; Extended Data Fig. 6c**). Similarly, administration of a small molecule inhibitor of NUPR1 nuclear import, ZZW-115 (ref.^40^) increased *ex vivo* tumor sphere formation of *KP* AT2 cells from aged mice, whereas sphere formation by *KP* AT2 cells from young mice was suppressed by NUPR1 inactivation (**Extended Data Fig. 6d**). Further, targeting *Nupr1* in aged mice led to an increase in the average size of the lung tumors (**Fig. 2h**). Loss of *Nupr1* in the aged tumors increased cancer cell proliferation to a similar level to that seen in the young control tumors, whereas the opposite—suppression of proliferation—was observed in the young *KP* cancer cells (**Fig. 2i**). To evaluate effects on molecular progression we isolated *KP-Cas9* cells from sg*Nupr1* or control sgRNA tumors for RNA-seq analysis. Loss of *Nupr1* permitted progression of aged *KP* lung tumors to the later molecular stages observed in young mice at this time point when compared to the aged controls, suggesting that *Nupr1* acts as a barrier to tumor progression in aged mice (**Fig. 2j; Extended Data Table 5**). We next interrogated the role of *Nupr1* in LUAD transformation as a function of age in the *ex vivo* transformation assay. Genetic inactivation of *Nupr1* strongly suppressed transformation of AT2 cells isolated from 3-month-old mice. However, the effect of *Nupr1* knockout gradually ameliorated with age when compared to control sgRNA, becoming neutral at 18 months, until finally showing tumor-suppressive activity at 24 months of age (**Fig. 2k**). Taken together, these results indicate that aging-associated induction of NUPR1 expression suppresses AT2 cell transformation via a cell-autonomous mechanism.

### Elevated expression of NUPR1 disrupts iron homeostasis in aged AT2 and LUAD cells

Given the suggested role of NUPR1 in controlling levels of cellular iron, we chelated iron using deferoxamine (DFO) in the context of NUPR1 loss-of-function in the *ex vivo* transformation assay. DFO blunted the increase in tumor sphere formation by *KP-Cas9* AT2 cells from aged mice in response to genetic or pharmacologic targeting of NUPR1 (**Fig. 3a; Extended Data Fig. 7a**). Conversely, DFO did not have an effect on young tumor spheres in the context of NUPR1 inactivation (**Fig. 3a; Extended Data Fig. 7a**). These findings indicate that the aging-associated induction of *Nupr1* expression leads to a functional iron deficiency in the AT2 cells. Consistent with this notion, supplementation of aged AT2 cells with transferrin-bound iron or ferric ammonium citrate (FAC) promoted tumor sphere formation in the *ex vivo* transformation assay (**Fig. 3b; Extended Data Fig. 7b**). Addition of transferrin-bound iron did not change the effect of *Nupr1* knockout in *ex vivo* transformation capacity of the young or aged AT2 cells, suggesting either that targeting NUPR1 or iron supplementation is sufficient for reversing the age-associated decline in tumorigenic potential (**Extended Data Fig. 7c**).

**Figure 3.**
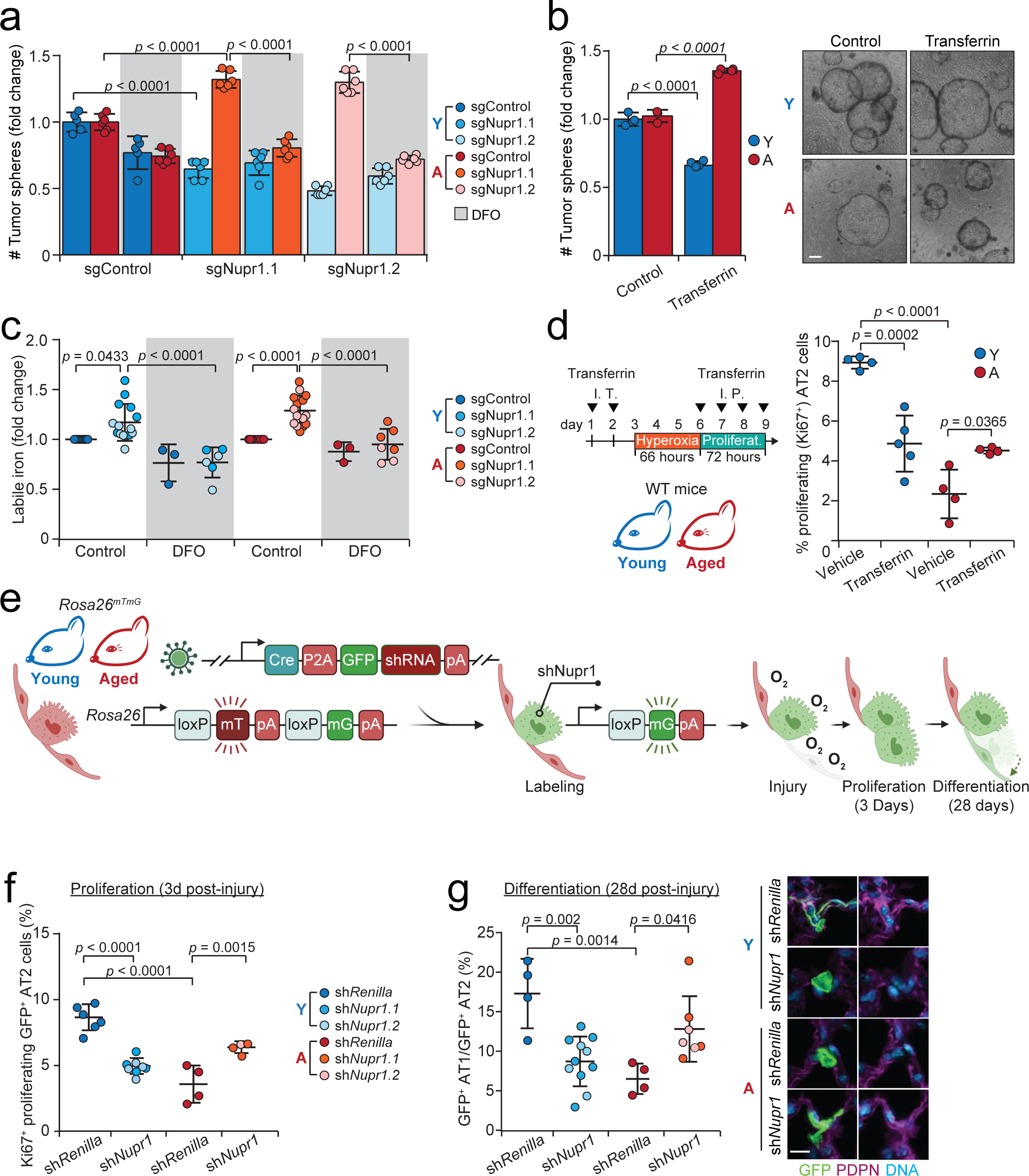
Elevated expression of NUPR1 disrupts iron homeostasis in aged AT2 and LUAD cells. (**a**) *Ex vivo* transformation of AT2 cells isolated from aged and young *KP-Cas9* mice with the indicated lentiviral vectors (Fig. 2f) delivering Cre + control sgRNA or two independent sgRNAs targeting *Nupr1*, with (grey shading) or without DFO treatment (*n* = 5-6 biological replicates). (**b**) *Ex vivo* transformation of AT2 cells isolated from aged and young *KP* mice and stimulated with transferrin (*n* = 3-4 biological replicates per group). Representative images of transformed tumor spheres are shown on the right. (**c**) Measurement of labile iron in LUAD cells derived from (**a**); data are normalized to control *N* = 9, 15, 3, 6, 8, 15, 3 and 7 biological replicates from left to right. (**d**) Proliferation of AT2 cells in young and aged C57BL/6 mice administered vehicle or transferrin, as indicated in the experimental scheme shown on the left. The percentage of Ki67^+^/SPC^+^ double positive cells per total SPC^+^ AT2 cells is shown. *N* = 4, 5, 4, and 4 mice in each group. (**e-g**) Evaluation of the proliferation and differentiation of AT2 cells following *Nupr1* knockdown *in vivo*. Experimental strategy is shown in (**e**). Note that cells expressing shRNAs and Cre recombinase were visualized by the switching of endogenous fluorescence from membrane-bound tdTomato (mT) to membrane-bound GFP (mG). Only GFP^+^ cells were analyzed in (**f-g**). Proliferating AT2 cells harboring the shRNAs were defined as Ki67^+^/GFP^+^/SPC^+^ three days after injury (**f**) and percentage of Ki67^+^ per GFP^+^/SPC^+^ AT2 cells was reported. Differentiated AT2 cells were counted as the percentage of traced AT1 cells (PDPN^+^/GFP^+^) per traced GFP^+^/SPC^+^ AT2 cells at 28 days after injury (**g**). *N* = 6, 8, 4, and 4 mice in each group in (**f**) and *N* = 4, 11, 4 and 7 mice in each group in (**g**). Representative images of GFP-labeled AT1 (PDPN^+^, top and bottom rows) and AT2 cells (PDPN^-^, middle rows) are shown on the right. Scale bar: 100 µm (**b**) and 10 µm (**g**). Mean with SD is shown in (**a-b**). One-way ANOVA was used in (**a-g**).

To directly test whether NUPR1 controls iron levels in transformed AT2 cells, we measured levels of labile cellular iron following *Nupr1* knockout in the AT2 *ex vivo* transformation assay. Loss of *Nupr1* promoted iron uptake by transformed AT2 cells isolated from either aged or young mice (**Fig. 3c; Extended Data Figure 7d**). Interestingly, the baseline levels of both labile and total iron were higher in the aged AT2 cells compared to the young (**Extended Data Figure 7e,f**), suggesting either a higher requirement of iron for cell growth or inefficient mobilization of stored iron in the aged AT2 cells.

We next investigated whether the functional iron insufficiency induced by elevated *Nupr1* expression underpinned loss of stemness and regenerative capacity of aged AT2 cells. Confirming our *ex vivo* results, provision of transferrin-bound iron or shRNA-mediated knockdown of *Nupr1* partially reversed the decline in AT2 cell <u>self-renewal capacity</u> following hyperoxia injury in aged mice *in vivo* but was detrimental to self-renewal of young AT2 cells (**Fig. 3d-f; Extended Data Fig. 7g**). Similar results were obtained in cultured AT2 organoids following transferrin supplementation or chemical inhibition of NUPR1 nuclear import (**Extended Data Fig. 7h,i**). To investigate <u>differentiation potential</u>, we performed lineage-tracing of AT2 cells in aged vs. young *Rosa26^mTmG/+^* mice by intra-tracheal delivery of lentiviral vectors harboring an shRNA targeting *Nupr1* or *Renilla* luciferase (control) and Cre recombinase, followed by hyperoxia injury (**Fig. 3e**). In this experiment, lineage-traced AT2 cells harboring the shRNA can be tracked by expression of GFP. Importantly, knockdown of *Nupr1* reversed the loss of AT1 differentiation in aged mice. However, the differentiation of young AT2 cells towards the AT1 lineage was diminished in response to *Nupr1* inactivation (**Fig. 3g**). Thus, age-associated functional iron insufficiency caused by induction of *Nupr1* suppresses stemness—the capacity for self-renewal and differentiation—of AT2 cells.

### NUPR1 induces functional iron insufficiency and ferroptosis resistance in aged AT2 cells via lipocalin-2

NUPR1 has been shown to protect pancreas cancer cells from excess iron by inducing expression of the iron sequestering protein lipocalin-2 (encoded by *Lcn2*)^37^. *Lcn2* was induced in the aged AT2 cells and LUAD cells (**Fig. 2a**; **Fig. 4a,b; Extended Data Fig. 8a**). We also detected induction of lipocalin-2 protein in the aged LUAD tumors, whereas knockout of *Nupr1* restored lipocalin-2 to similar levels as in the young tumors (**Fig. 4b**). Similarly, *Lcn2* was suppressed in response to NUPR1 targeting in both aged and young AT2 cells in the *ex vivo* transformation assay and in wild-type primary AT2 cells *in vitro* and *in vivo* (**Extended Data Fig. 8b-e**). Importantly, knockout of *Lcn2* increased tumor sphere formation capacity of aged AT2 cells and suppressed the transformation of young AT2 cells (**Fig. 4c-e**), essentially recapitulating the phenotypes observed upon *Nupr1* knockout (**Fig. 3a,c**). In contrast, knockout of a panel of other key iron transport and storage genes was equally detrimental to both young and aged AT2 cells in the transformation assay, indicating the NUPR1 – lipocalin-2 axis is unique in possessing such age-context dependent function (**Extended Data Fig. 8f**). We next investigated the significance of the genetic interaction between *Nupr1* and *Lcn2* in aged and young AT2 cells. *Lcn2* cDNA fully abrogated rescue of AT2 cell transformation and iron uptake in cells with *Nupr1* knockout (**Fig. 4e,f**), directly implicating *Lcn2* induction downstream of NUPR1 as a mechanism underpinning age-associated functional iron insufficiency and loss of AT2 transformation capacity. In orthogonal support with this finding, supplementation with recombinant lipocalin-2 blunted the increase in aged AT2 tumor sphere formation in response to chemical NUPR1 inhibition with ZZW-115 (**Extended Data Fig. 8g**).

**Figure 4.**
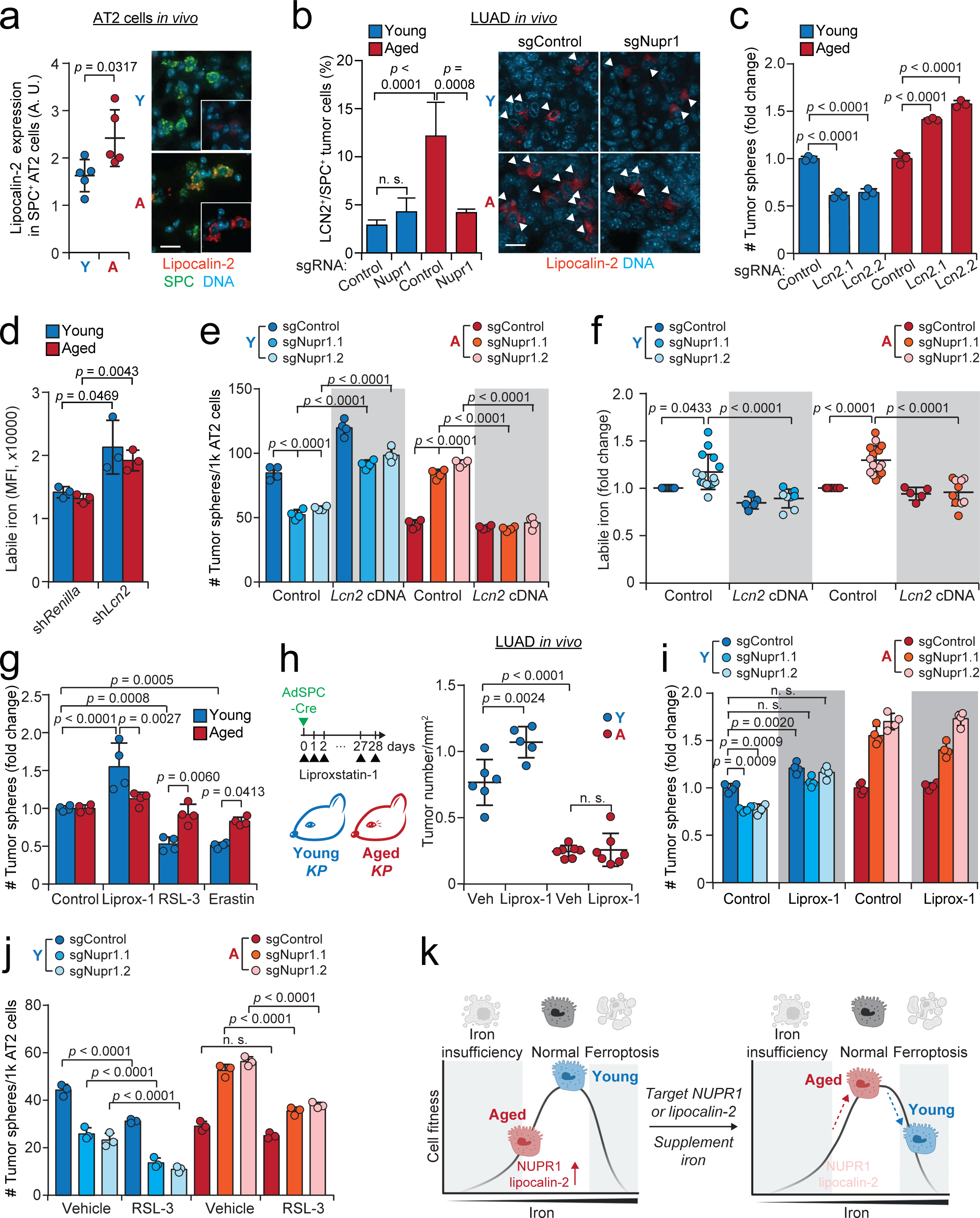
NUPR1 drives functional iron insufficiency and ferroptosis resistance in aged AT2 cells via lipocalin-2. (**a**) Quantification of lipocalin-2 immunofluorescence intensity in SPC^+^ AT2 cells (*n* = 5 young and 5 aged animals). Representative images are shown on the right. A. U.: arbitrary units. (**b**) Percentage of lipocalin-2^+^ cells in SPC^+^ *KP-Cas9* LUAD cells in tumors at 12 weeks post-tumor initiation. Representative images of lipocalin-2 immunofluorescence are shown on the right (*n* = 31, 30, 10, and 31 tumors for each condition, left to right). Arrowheads indicate lipocalin-2 positive AT2 cells. Note that the SPC staining used to identify AT2 cells is not shown for clarity. (**c**) *Ex vivo* transformation of AT2 cells isolated from aged and young *KP-Cas9* mice using the indicated lentiviral vectors (Fig. 2f) delivering Cre + control sgRNA or two independent sgRNAs targeting *Lcn2* (*n* = 3 biological replicates per condition). (**d**) Measurement of labile iron by Biotracker Ferro Far-red dye in *KP* LUAD cells derived from *ex vivo* transformation of aged and young AT2 cells with control (shRenilla) or shRNA targeting *Lcn2* (*n* = 3 biological replicates). Median fluorescence intensity (MFI) of Ferro Far-red dye staining was calculated; data is shown normalized to control shRNA. (**e**) *Ex vivo* transformation of AT2 cells isolated from aged and young *KP-Cas9* mice with sgRNAs targeting *Nupr1*, with or without *Lcn2* cDNA (grey background). *N* = 4 biological replicates per group. (**f**) Measurement of labile iron in LUAD cells derived from (**e**); data is shown normalized to control shRNA. *N* = 9, 15, 5, 8, 8, 15, 5, and 10 biological replicates from left to right, respectively. Note that data points of sgControl and sgNupr1 without *Lcn2* cDNA is the same as in Fig. 3c. (**g**) *Ex vivo* transformation of aged and young AT2 cells derived from *KP* mice subjected to the indicated chemicals. Fold change in tumor sphere number normalized to the control group is shown. *N* = 4 biological replicates in each group. (**h**) Tumor number in aged vs. young *KP* mice 4 weeks post-tumor initiation with adeno-SPC-Cre, with or without daily systemic administration of liproxstatin-1 (10 mg/kg of body weight). *N* = 6, 5, 7, and 7 mice in each group. (**i**) *Ex vivo* transformation of AT2 cells isolated from aged and young *KP-Cas9* mice with sgRNAs targeting *Nupr1*, with or without liproxstatin-1. *N* = 4 biological replicates in each group. (**j**) *Ex vivo* transformation of AT2 cells isolated from aged and young *KP-Cas9* mice with sgRNAs targeting *Nupr1*, with or without RSL-3. *N* = 3 biological replicates. (**k**) Model showing the age-dependent function of *Nupr1*. Briefly, induction of the NUPR1—lipocalin-2 axis with aging leads to a functional iron insufficiency in aged AT2 cells, which leads to reduced regenerative and tumorigenic capacity. Targeting the NUPR1-lipocalin-2 axis or iron supplementation rescues the impaired cellular fitness of aged AT2 cells. However, inactivation of the NUPR1-lipocalin-2 axis or excessive iron promotes ferroptosis in young AT2 cells. Mean with SEM is shown in (**b**). Mean with SD is shown in (**c-e**), (**g**), (**i**) and (**j**). Y: young; A: aged. Scale bar: 20 µm. *Mann-Whitney* test was used in (**a**). Student’s *t* test was used in (**d**). One-way ANOVA was used in (**b-c**) and (**e-j**).

Our results implicate NUPR1 as an upstream driver of *Lcn2* expression in AT2 and LUAD cells and implicates overactivation of the NUPR1–lipocalin-2 axis in mediating loss of stemness and tumorigenic potential in aged AT2 cells. On the other hand, inactivation of the NUPR1 – lipocalin-2 axis was detrimental to young AT2 cells both in the context of transformation and regeneration. Given the NUPR1–lipocalin-2 axis has been implicated in ferroptosis defense^37^, we hypothesized that young AT2 cells are more sensitive to ferroptosis than aged AT2 cells. Indeed, induction of ferroptosis by chemical or genetic suppression of glutathione peroxidase-4 (Gpx4) impaired *ex vivo* tumor sphere formation by young AT2 cells, whereas aged AT2 cells were resistant (**Extended Data Fig. 8h,i**). Conversely, inhibition of ferroptosis with liproxstatin-1, a radical-trapping agent highly active in lipid bilayers, promoted transformation of young AT2 cells *ex vivo* and *in vivo*, whereas, again, little effect was observed the aged AT2 cells (**Fig. 4h,i**). Furthermore, liproxstatin-1 treatment reversed the suppression of young AT2 cell transformation by chemical inhibition of NUPR1 (**Extended Data Fig. 8j**). *Nupr1* knockout exacerbated the effect of the chemical ferroptosis inducer RSL-3 in young cells and sensitized aged AT2 cells to RSL-3 (**Fig. 4j**). Similarly, supplementation with iron-bound transferrin sensitized aged AT2 cells to RSL-3 (**Extended Data Fig. 8k**). In sum, our results show age-associated induction of the NUPR-1– lipocalin-2 axis shifts the balance of iron homeostasis towards a functional iron insufficiency, which suppresses transformation and regeneration of AT2 cells while rendering them resistant to ferroptosis. Conversely, young cells maintain sufficient iron levels for growth, but are sensitive to ferroptosis (**Fig. 4k**).

### DNA hypomethylation underpins aging-associated induction of the NUPR1–lipocalin-2 axis

Given its prominence as a primary hallmark of aging^15^, we investigated the role of DNA methylation in the age-associated gene expression changes shared by AT2 cells and cancer cells. We assessed DNA methylation on a genome-wide scale in young and aged AT2 cells and in age-matched LUAD tumors (**Extended Data Fig. 9**). Consistent with findings in other cell types^15–19^, we observed a global loss of DNA methylation punctuated by CpG island-specific hypermethylation in the aged AT2 and LUAD cells and found that age was the most significant driver of unsupervised sample clustering (**Extended Data Fig. 9c,d, g, h; Extended Data Table 6; Extended Data Table 7**). To connect differentially methylated cytosines (DMCs) with promoters and enhancers in AT2 cells, we mapped DMCs to histone-3 (H3)K4me1, H3K27ac, and H3K4me3 modifications^41^ (**Extended Data Fig. 10a; Extended Data Table 8**). The most distinct difference in the number of hypermethylated and hypomethylated DMCs between aged and young samples was observed at active enhancers marked by H3K4me1 and H3K27ac (**Extended Data Fig. 10a; Extended Data Table 8**). We observed statistically significant correlations between DNA hypomethylation at promoters and enhancers of the shared set of genes changed with aging in both AT2 and LUAD cells (**Fig. 2a**; **Fig. 5a**). In aged relative to young AT2 cells, *Nupr1* was hypomethylated within an intron marked by H3K4me3 and H3K27ac and at a site marking an active enhancer (H3K4me1 and H3K27ac), whereas *Lcn2* was demethylated at an active enhancer site (H3K4me, H3K4me3, and H3K27ac) (**Extended Data Fig. 10b-d; Extended Data Table 9**). Loss of DNA methylation at these key gene-regulatory regions is consistent with epigenetic derepression of gene expression with age^18^. Similar to the AT2 cells, aging-associated DNA methylation levels between AT2 and LUAD were positively correlated at the genes that make up the age-associated transcriptomic signature shared by AT2 and LUAD cells (**Fig. 5b; Extended Data Table 3)**.

**Figure 5.**
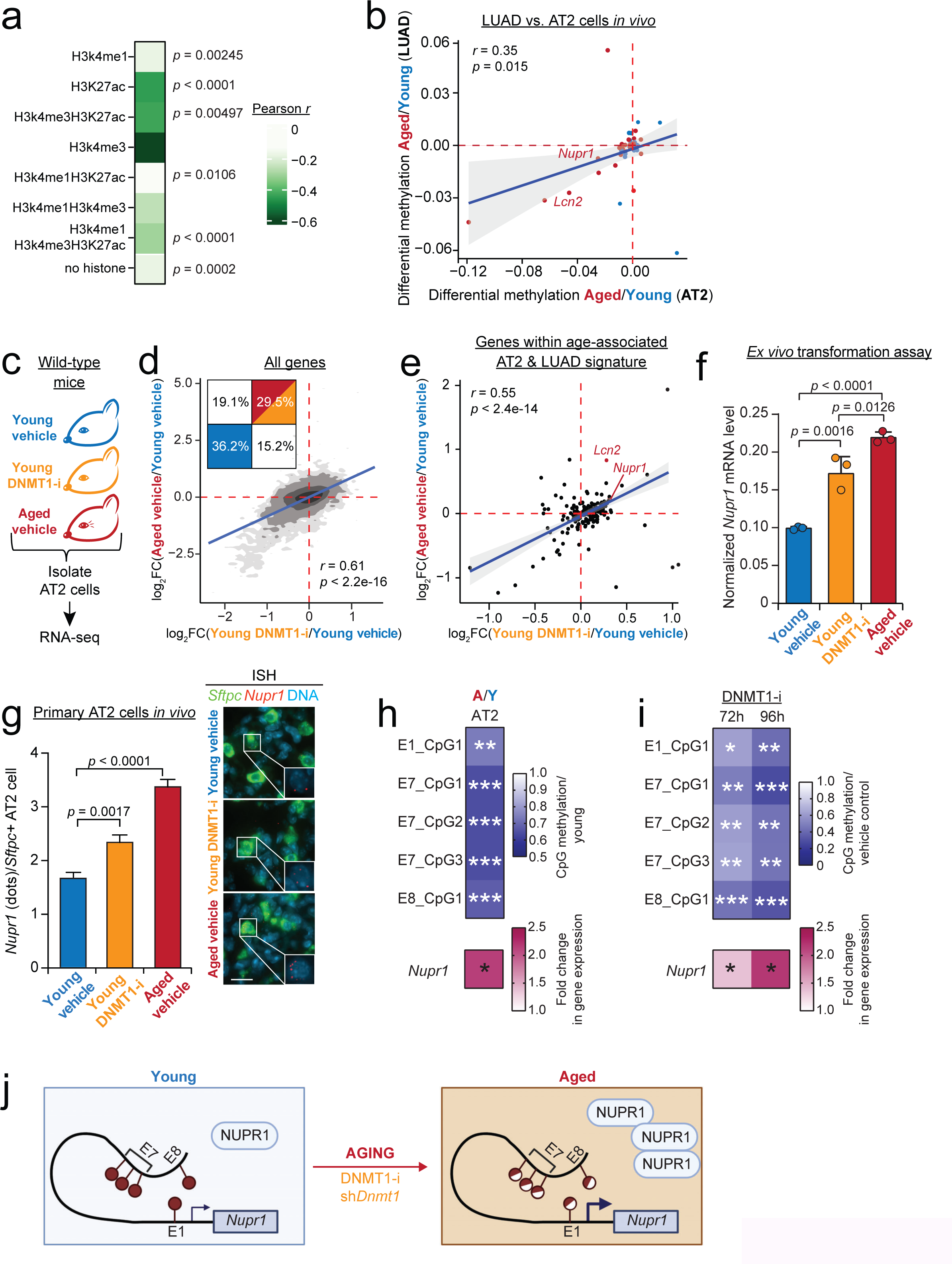
DNA hypomethylation underpins aging-associated induction of the NUPR1– lipocalin-2 axis. (**a**) Heatmap illustrating the correlation between differentially methylated cytosines (DMCs) and differentially expressed genes (DEGs) in AT2 cells (FDR < 0.05) in each AT2 histone mark category. The color bar indicates the Pearson correlation coefficient. (**b**) Scatter plot showing mean differential methylation level of promoter regions (1 kb upstream and 200 bp downstream of the TSS) of the top 25 upregulated and downregulated genes among the aging-associated signature shared by both AT2 and LUAD cells (Fig. 2b**; Extended Data Table 3**). Pearson correlation was calculated. The regression line (blue) and its confidence interval (light grey) were plotted using a linear model. Genes upregulated in the aged AT2 and LUAD cells are shown in red; genes upregulated in the young are marked in blue. (**c**) DNMT1 inhibition (DNMT1-i, orange) with GSK-3484862 over 8 days *in vivo*. (**d**) Contour plot demonstrating the correlation between the fold change (FC) of DEGs in AT2 cells isolated from young mice administered DNMT1-i or vehicle (*x*-axis) vs. aged or young vehicle-treated mice (*y*-axis). The grayscale represents the probability distribution. Pearson correlation was calculated. The percentage of genes in each quadrant is significantly different from random (binomial test *p* < 2.2^-16^). The regression line (blue) was plotted using a linear model. (**e**) Scatter plot showing the correlation between the fold change of DEGs from the signature shared by AT2 cells and LUAD cells (Fig. 2b**; Extended Data Table 3**) in AT2 cells isolated from young mice administered DNMT1-i or vehicle (*x*-axis) vs. aged or young vehicle-treated mice (*y*-axis). Pearson correlation was calculated. The regression line (blue) and its confidence interval (light grey) were plotted using a linear model. (**f**) Expression of *Nupr1* (normalized to the housekeeping gene *Gapdh*) in lung cancer cells following *ex vivo* transformation of AT2 cells from aged vs. young mice, with or without DNMT1 inhibition (*n* = 3 technical replicates). (**g**) Quantification of *Nupr1* mRNA molecules in *Sftpc^+^* AT2 cells in young mice administered vehicle (blue) or DNMT1-i (orange) for 8 days, compared to aged mice that received vehicle (red) (n = 353, 438, and 429 AT2 cells per condition, respectively). Scale bar: 20 µm. (**h-i**) Change in methylation of defined CpGs at *Nupr1* distal enhancers identified in this study (**Extended Data Fig. 12**). Change in CpG methylation in primary AT2 cells from aged vs. young C57BL/6 mice measured by EM-Seq (**h**, *n* = 4 mice per group) and cultured *KP* lung cancer cells subjected to DNMT1 inhibition for 72 or 96 h measured by pyrosequencing (**i**, *n* = 4 independent cell lines). Results are shown as fold change normalized to young AT2 cells (**h**) or vehicle-treated cells (**i**). The pink heatmap in the bottom shows expression of *Nupr1* in the above conditions. *N* = 7 young and 7 aged primary AT2 samples (**h**) and *n* = 4 *KP* lung cancer cell lines (**i**). (**j**) Aging-associated hypomethylation at distinct CpGs at novel *Nupr1* enhancers associates with induction of *Nupr1* in aged AT2 and LUAD cells and young cells subjected to DNMT1 inhibition. *, *p* < 0.05, **, *p* < 0.01, ***, *p* < 0.001. Numerical *p* value of (**h**) and (**i**) was shown in **Extended Data Table 19.** Mean with SEM is shown in (**g**). One-way ANOVA was used in (**f-g**). Student’s *t* test was used in (**i**).

To functionally interrogate whether DNA hypomethylation underpins the induction of the age-associated gene expression signature in AT2 cells, we suppressed DNA methylation in young wild-type C57BL/6J mice by systemic administration of GSK-3484862, a small molecule inhibitor of DNA methyltransferase-1 (DNMT1-i)^42^, for 8 days (**Fig. 5c**). We observed a significant reduction in 5’-methylcytosine (5-mC) staining in AT2 cells of mice subjected to DNMT1-i (**Extended Data Fig. 11a**), consistent with inhibition of DNMT1 activity. Young AT2 cells subjected to DNMT1-i showed a strong correlation with the transcriptomic changes observed in aged AT2 cells, implicating global progressive DNA hypomethylation as a key mechanistic driver of the aging transcriptome in AT2 cells (**Fig. 5d; Extended Data Table 10**). Within the age-associated gene expression signature shared between AT2 and LUAD cells (**Fig. 2a**), *Nupr1* and *Lcn2* were among the top genes induced by DNMT1-i in AT2 cells from young DNMT1-i treated mice (**Fig. 5e; Extended Data Table 10**), which we confirmed by quantitative PCR and *in situ* detection of *Nupr1* mRNA (**Fig. 5f,g**). We next interrogated the role of DNA methylation in cancer cells. Remarkably, DNMT1-i specifically suppressed the transformation of young AT2 cells, but not aged AT2 cells (**Extended Data Fig. 11b**). DNMT1-i induced *Nupr1* and *Lcn2* gene expression also in this context (**Extended Data Fig. 11c,d**). To rule out possible off-target effects of the DNMT1-i, we suppressed *Dnmt1* via lentiviral Cre-shRNA in the *Rosa26^mTmG^* mice *in vivo* (**Extended Data Fig. 11e-i**). Similar to DNMT1-i, sh*Dnmt1* induced an aged-like transcriptome in the young AT2 cells, including induction of *Nupr1* (**Extended Data Fig. 11h-j**).

To precisely pinpoint enhancers whose hypomethylation associates with induction of *Nupr1* gene expression in aged cells, we identified 11 candidate enhancer sites ±500 kb of the *Nupr1* transcription start site (TSS). We defined candidate enhancers based on chromatin accessibility, aging-associated hypomethylation of CpG sites located within methylation-sensitive binding motifs of transcription factors known to induce *Nupr1* gene expression^38^, and presence of active enhancer histone marks in AT2 cells^41^ (**Extended Data Fig. 10b-d; Extended Data Fig. 12a,b; Extended Data Table 9**). To identify functional *Nupr1* enhancers, we repressed each candidate enhancer in an arrayed CRISPR interference (CRISPRi) screen in three independent *KP* LUAD cell lines (**Extended Data Fig. 12c; Extended Data Tables 12 & 13**). This effort identified E1 (25,846 bp upstream of the *Nupr1* TSS), E7 (113,969 bp upstream), and E8 (114,169 bp upstream) as high-confidence *Nupr1* enhancers (**Extended Data Fig. 12d**). A high-resolution examination identified hypomethylation of one CpG in E1, three in E7, and one in E8 that specifically associated with aging in AT2 cells (**Fig. 5h; Extended Data Fig. 12e; Extended Data Fig. 13a**). Importantly, these CpGs showed significant hypomethylation in response to DNMT1 inhibition by target-specific pyrosequencing (**Extended Data Fig. 13b**), which correlated with elevated *Nupr1* gene expression following DNMT1 inhibition (**Fig. 5i; Extended Data Fig. 13b, Extended Data Table 14**). Furthermore, we note that repressing E1, E7, or E8 with CRISPRi led to a ∼50% reduction in *Nupr1* levels (**Extended Data Fig. 12d**), indicating these enhancers robustly regulate *Nupr1* expression. Increased hypomethylation-associated activity of these enhancers in concert is likely sufficient to explain the increase in *Nupr1* gene expression we observed in aged AT2 cells. These findings are consistent with a model whereby hypomethylation of *Nupr1* enhancers— perhaps with contribution from promoter and/or intronic hypomethylation—promotes age-associated induction of *Nupr1* expression (**Fig. 5j**).

### Aging-induced changes in AT2 cells and LUAD are conserved in humans and in adult stem cells outside the lung

To investigate the relevance of our findings to human aging, we examined tissue sections of aged and young human lungs. Similar to our findings in mice, we observed a decline in the density of AT2 cells in aged human lungs (**Fig. 6a; Extended Data Table 15**). The aged human AT2 cells displayed elevated expression of both NUPR1 and lipocalin-2 (**Fig. 6b; Extended Data Fig. 14a**). Notably, stimulation with transferrin-bound iron robustly promoted formation of organoids by human primary AT2 cells isolated from individuals 65-78 years of age (**Fig. 6c; Extended Data Table 16**), indicating that availability of iron is growth-limiting also in aged human AT2 cells. We next analyzed age-associated gene expression in LUAD bulk RNA sequencing data in The Cancer Genome Atlas^43^. Orthologs of genes expressed significantly higher in young mouse LUAD tumors showed an inverse correlation with human LUAD patient age, whereas orthologs induced in aged mouse LUAD tumors exhibited a positive correlation with human LUAD patient age (**Fig. 6d; Extended Data Table 17**). *NUPR1* expression was higher in resected human LUAD tissues from aged (>80-year-old) patients when compared to young or middle-aged (<55-year-old) patients (**Fig. 6e; Extended Data Table 16**). In sum, these findings demonstrate notable conservation of age-associated changes in AT2 cells and lung tumors across mice and humans.

**Figure 6.**
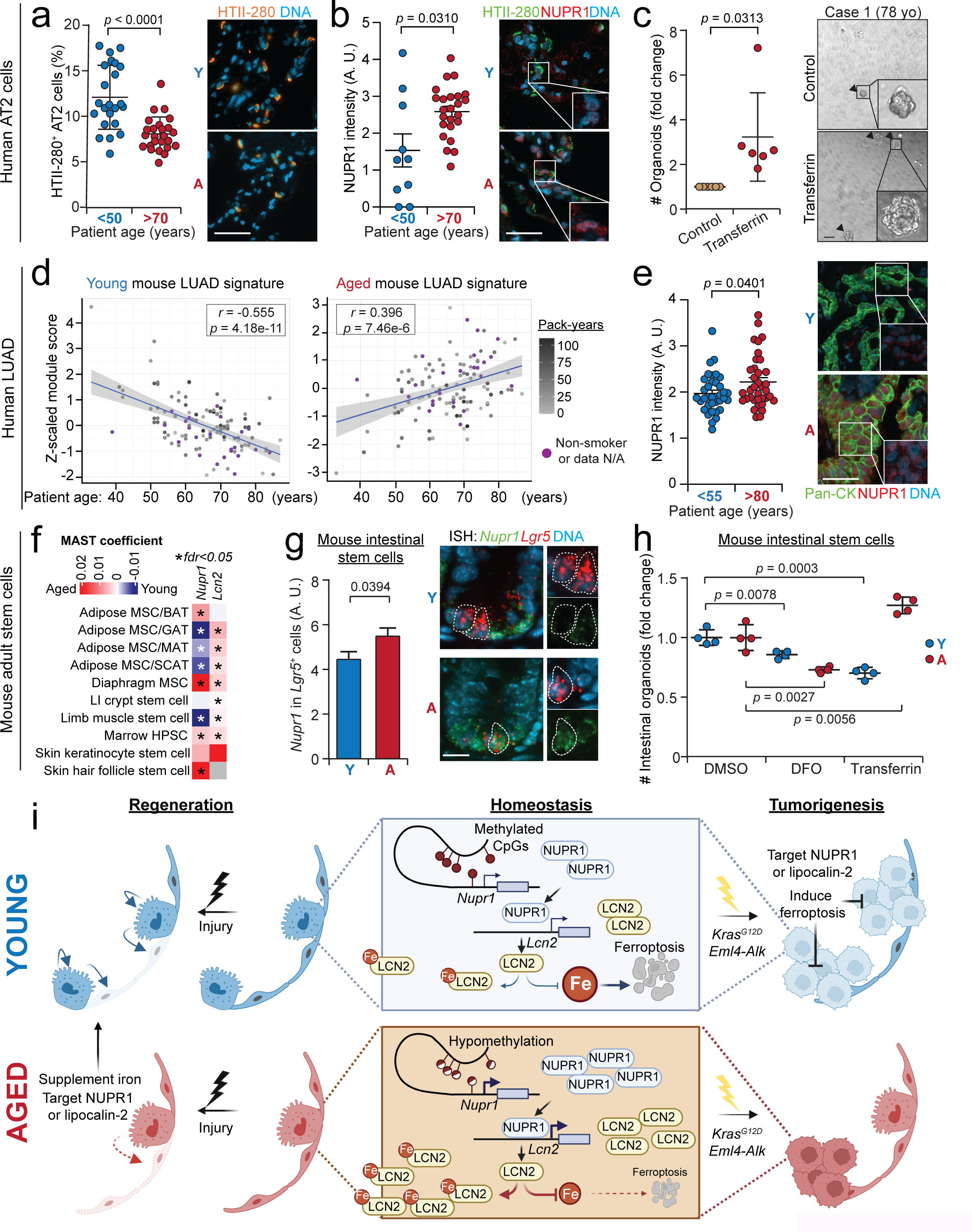
Aging-induced changes in AT2 cells and LUAD are conserved in humans and in adult stem cells outside the lung. (**a**) Quantification of AT2 cell density in healthy lungs of aged (>70 years old) and young or middle-aged (<50 years old) humans (*n* = 23 young/middle-aged and 25 aged patients, respectively). The proportion of HTII-280 positive AT2 cells (orange) per total lung cell number is shown. (**b**) Quantification of NUPR1 protein in healthy lung tissues of aged (>70 years old) and young or middle-aged (<50 years old) humans (*n* = 10 young/middle-aged and 24 aged patients, respectively). The intensity of NUPR1 (red) immunofluorescence was quantified only in cells expressing the AT2 cell marker HTII-280 (green). A. U.: arbitrary units. Note that HTII-280 signal (green) was not shown in the insets for NUPR1 (red) signal clarity. (**c**) Quantification of alveolar organoids formed by primary human AT2 cells in response to stimulation with iron-loaded human transferrin. Results are normalized to the number of control organoids. *N* = 6 patients. Scale bar: 50 µm. (**d**) Correlation of young mouse LUAD and aged mouse LUAD gene expression signatures with patient age in The Cancer Genome Atlas data. All tumors had a *KRAS* mutation. Smoking history of the patients is shown by gray shading. (**e**) NUPR1 immunofluorescence in LUAD tissue obtained from aged (>80 years old, *n* = 35) and young or middle-aged (<55 years old, *n* = 36) patients. NUPR1 signal intensity was measured in pan-cytokeratin (pan-CK) positive cancer cells. (**f**) Aging-associated changes in *Nupr1* and *Lcn2* expression in aged vs. young adult stem cells in previously published datasets^44,45^. The color gradient represents the gene expression change coefficient using MAST (red: upregulated in aged; blue: upregulated in young; grey: no difference due to low expression levels) as log_2_ fold change increasing or decreasing with aging. *, FDR<0.05. Numerical FDR value was shown in **Extended Data Table 19**. MSC, mesenchymal stem cell; BAT, brown adipose tissue; GAT, gonadal adipose tissue; MAT, mesenteric adipose tissue; SCAT, subcutaneous adipose tissue; LI, large intestine; HPSC, hematopoietic stem cell. (**g**) Quantification of *Nupr1* mRNA molecules in *Lgr5^+^* intestinal stem cells in young vs. aged mice by *in situ* hybridization (*n* = 291 and 286 *Lgr5+* cells from 3 aged and 3 young mice, respectively). (**h**) Quantification of organoid formation by intestinal crypts isolated from aged vs. young mice, with or without DFO or transferrin (*n* = 4 technical replicates per condition from one representative experiment that was repeated 3 times). (**i**) Graphical summary of findings in this study (see text). Scale bar: 50 µm in (**a-e**) and 10 µm in (**g**). Mean with SEM was shown in (**g**). Y: young; A: aged. Student’s *t* test was used in (**a**), (**e**), (**g**) and (**h**). *Mann-Whitney* test was used in (**b-c**).

We next asked whether our findings in the aged AT2 cells are generalizable to other adult stem cell types. Either *Nupr1*, *Lcn2*, or both were significantly induced with aging in 9/13 of adult stem cell types examined in published scRNA-seq datasets^44,45^ (**Fig. 6f**). Given the availability of well-established protocols for organoid culture, we chose to perform functional validation in the mouse small intestine. *Nupr1* and *Lcn2* mRNA were elevated *in situ* in *Lgr5*^+^ intestinal stem cells in aged mice compared to the young (**Fig. 6g; Extended Data Fig. 14b**). Similar to the AT2 cells, supplementation with transferrin-bound iron stimulated the growth of aged intestinal crypt organoids but was detrimental to young crypts (**Fig. 6h**). Furthermore, crypt organoids from aged mice were resistant to ferroptosis induced by RSL-3, whereas young crypts were sensitive **(Extended Data Fig. 14c**). These results suggest induction of the NUPR1–lipocalin-2 axis, functional iron insufficiency, and ferroptosis resistance may be broadly important properties of aged adult stem cells.

## Discussion

Accumulation of somatic mutations and loss of stemness occur in parallel over the course of organismal aging. The significance of this relationship for cancer development has been a subject of debate^5,12^, yet experimental studies directly addressing the capacity of aged stem cells or progenitors to transform *in vivo* have been lacking. By utilizing an autochthonous GEMM of lung cancer, *in vivo* functional genetics, primary organoid cultures, and human tissue we find that age-associated decline of lung AT2 cell stemness is tightly coupled to a reduction in tumor initiation potential. This result constitutes direct evidence for a model whereby aging-associated loss of stemness is tumor suppressive in the lung, precluding transformation of cells that acquire oncogenic mutations once stemness falls below a threshold required to initiate tumors. Thus, our results may partially explain the decline in cancer incidence that is observed in very aged individuals (>80-85-year-olds)^9^, including lung cancer, although selection for individuals resistant to cancer development via other mechanisms in this population is likely to be a significant contributor. Moreover, given that stem cells decline in all tissues with aging, this model may be broadly applicable to other cancer types.

Although we find aging-associated changes intrinsic in the cell-of-origin are tumor suppressive, our initial results utilizing heterochronic transplantation suggest the aged lung is a more hospitable microenvironment for tumor growth compared to the young (**Fig. 1j**). Similar observations have been made by others utilizing cell line transplantation in other cancer contexts^20^. Mechanisms rendering aged hosts more permissible include a decline in anti-tumor immunity (“immunosenescence”)^46^, metabolic changes^47^, and alterations in niche factors^20,48^. Thus, our results and the work of others imply that the dramatic increase in cancer incidence in 50-70-year-olds may result from pre-malignant lesions arising in relatively young individuals whose stem cells harbor high tumorigenic potential. These initial lesions may then eventually progress to clinically detectable disease as a result of mutation accumulation and development of a permissive microenvironment during aging. Indeed, pre-malignant lesions are routinely detected in autopsies of young and middle-aged adults. Further, a recent longitudinal study linked initial oncogenic events detected in young patients to lung cancers arising in older age^49^. This, in light with our findings, implies efforts aimed at cancer prevention and early interception may be best directed to the critical window in young individuals when stem cells are most susceptible to transformation. Finally, these findings raise an interesting question of whether initial oncogenic events acquired at a young age can protect the cell-of-origin from the decline in stemness and tumorigenic potential associated with aging.

Our work casts dysregulation of iron homeostasis as a mechanistic culprit for loss of AT2 cell stemness and tumorigenic potential. We reveal that aging leads to induction of NUPR1 expression that promotes exclusion of iron from the AT2 cells via upregulation of the iron sequestering protein lipocalin-2 (refs.^37,38^). The NUPR1–lipocalin-2 axis has been implicated in sensing and controlling cellular iron levels and protecting cells from ferroptosis^13,17,50^. To avoid ferroptosis, cells must balance their iron uptake with the levels of ambient oxygen, while ensuring sufficient supply for essential enzymatic reactions. This balance may be particularly precarious in the lung epithelium, which is chronically exposed to high levels of oxygen relative to most tissues. We find aged AT2 cells shift this balance by over-activating the NUPR1–lipocalin-2 axis, which leads to functional iron insufficiency and resistance to ferroptosis. To our knowledge, ours is the first demonstration that aged cells are ferroptosis resistant, which has important ramifications regarding use of ferroptosis inhibitors in regenerative medicine and organ transplantation and ferroptosis inducers in cancer therapy^50,51^. Based on our findings these therapies may be more effective in young patients (**Fig. 6i**).

Interestingly, like other cell types in aged tissues^52^, we found higher baseline levels of total iron in aged AT2 cells than in their young counterparts, suggesting that aged AT2 cells require more iron to maintain stemness and tumorigenic potential. Another possibility is that the aged AT2 cells suffer from inability to mobilize sufficient iron from cellular iron stores, maintained by ferritin and other iron-binding proteins^52^, including lipocalin-2. Future studies are needed to elucidate the mechanistic underpinnings of functional iron insufficiency in aged AT2 cells despite their higher absolute iron levels when compared to the young.

The availability of iron-sulfur clusters, key components of many essential enzymes, is rate-limiting for LUAD growth in high oxygen environments^53^, suggesting one mechanism whereby iron insufficiency may suppress tumorigenesis in the lung. Iron is also an essential co-factor for a range of enzymatic reactions, including the demethylation of histones and DNA – processes critical to cellular differentiation. In line with this, we found aged AT2 cells to be severely hindered from differentiating into AT1 cells upon injury, which was rescued by targeting NUPR1. Thus, both aspects of AT2 cell stemness, self-renewal and differentiation, are directly suppressed by age-associated functional iron insufficiency caused by over-activity of the NUPR1–lipocalin-2 axis. Interestingly, we found elevated expression of *Nupr1*, *Lcn2*, or both in multiple aged adult stem cell compartments, and demonstrated that growth of aged small intestinal stem cells was similarly limited by availability of iron. As such, our findings cast inhibition of the NUPR1–lipocalin-2 axis or iron supplementation as promising strategies for rejuvenating aged AT2 cells and possibly other adult stem cell types in regenerative medicine. Conversely, age-associated induction of the NUPR1–lipocalin-2 axis may be tumor-suppressive in adult stem cells broadly beyond AT2 cells.

In addition to its importance for physiologic differentiation, iron has been shown to be essential for epigenetic remodeling in cancer. A recent elegant study found iron is required for the demethylation of repressive histone marks critical to epithelial-mesenchymal transition (EMT) in cancer cells^54^. In line with this work, we found iron insufficiency driven by NUPR1 suppressed progression of lung tumors that managed to initiate in aged mice, suggesting that iron may be required for the cancer cell state transitions that define lung cancer progression^23^. The iron-dependent TET-family methylcytosine dioxygenases and histone demethylases are possible candidates responsible for the epigenetic remodeling enabling these cell state transitions, whose activity may be impaired by iron insufficiency^55^.

Our results indicate that aging-associated DNA hypomethylation at *Nupr1* enhancers is permissive for upregulation of *Nupr1* expression, which may be induced by various forms of cellular stress^37,38^, including the accumulation of reactive oxygen species, ER stress, DNA damage, and inflammation, all of which occur during aging^13,14^. Elevated activity of tumor suppressors, including p53, has been shown to promote age-associated decline of stem cells^56^. Interestingly, *Nupr1* is among a set of genes induced by p53 activation in AT2 cells during regeneration^57^. The combination of stress inputs and the epigenetically permissive hypomethylated state provide a mechanistic explanation for the aberrantly high levels of *Nupr1* in the aged AT2 cells, which deprives the AT2 cells of sufficient iron, leading to loss of stemness and tumorigenic potential.

We found shared age-dependent changes in gene expression between the AT2 cells and LUAD cells, suggesting that functionally important aging-associated changes may be inherited by the LUAD tumors from their origin, the AT2 cells. Our results indicate that this form of age-dependent inheritance is an important force shaping the epigenetic context in which oncogenic mutations are acquired, altering the oncogenic competence of the cell of origin and the molecular features of emergent cancers as a function of organismal age. Such age-dependent differences may enable personalization of cancer prevention and treatment strategies based on the age of the patient.

## Acknowledgments

We thank K. Birsoy, L. Jones, T. Papagiannakopoulos, L. Parada, and C.M. Rudin for critical comments on the manuscript; M. Conrad, A.-K. Hadjantonakis, D. Huangfu, A. Koff, C. Sawyers, L. Studer, M. Winslow, and members of the Tammela laboratory for helpful discussions; M. Conrad for advice on experiments involving ferroptosis; S. Persad and D. Pe’er for help with SEACells algorithm; D. Buenocore, J. Silber, R. Spencer, and K. Ventura for help with tissue microarray generation; A. Chavez Perez and S. Lowe for help with senescence-associated β-galactosidase and p16 staining; P.J. Hamard for epigenetics experiments; R. Gardner for FACS support; K. Manova for histology support; E. Chan and E. Rosiek for help with image analysis and quantification; A. Alonso for low-input DNA Methyl-Seq analysis; C. Cobbs, N. Mohibullah, and A. Viale for next-generation sequencing; J. Agnis, J. Lisanti, and C.M. Rudin for primary patient lung tissue; H. Alcorn and O. Grbovic-Huezo for laboratory management; and J. Chan, M. Gregory, G. Hartmann, A. Hudson, H. Styers, C. Sussman, and S. Torborg for help with experiments; and the Genetic Resources Core Facility, Johns Hopkins University School of Medicine (*RRID:SCR_018669*) and R. Ashworth and J. Totey, in particular, for their assistance with the pyrosequencing assay design and sequencing data. This work was supported by the Mark Foundation for Cancer Research Emerging Leader Award (to T.T.), the Go2 Foundation for Lung Cancer (to T.T. and M.J.B.), and by the NIH/NCI Cancer Center Support Grant P30-CA08748 (to MSKCC). X.Z. received support from a training award from New York Stem Cell Science NYSTEM (#C32559GG) and the Center for Stem Cell Biology at MSKCC, and S.J. from the Hope Funds for Cancer Research. R.C. is supported by The Alan and Sandra Gerry Foundation. E.S.W. is supported by a NHMRC Investigator Grant (GNT2009309), ARC Discovery Project (DP200100250), and a Snow Medical Fellowship. T.T. is supported by Josie Robertson, American Cancer Society, Rita Allen, and V Foundation Scholarships. We acknowledge the use of the Integrated Genomics Operation Core (funded by Cycle for Survival, and the Marie-Josée and Henry R. Kravis Center for Molecular Oncology), the Flow Cytometry, the Laboratory of Comparative Pathology, and Histology Core Facilities at Sloan Kettering Institute, funded by CCSG P30-CA08748. We also acknowledge The Victor Chang Cardiac Research Institute Innovation Centre (funded by the New South Wales Government Ministry of Health). Some schematic figures were created with BioRender.

## Contributions

X.Z. and T. Tammela conceived and designed the study and wrote the manuscript. X.Z., E.S.W., and T. Tammela interpreted the data. E.S.W., Q.W., S.J., M.B., and S.D. contributed to the writing of the manuscript. X.Z., M.B., S.D., K.B., and Z.L. performed experiments and data analysis. Q.W., S.J., A.F., D.H., Y.Z., R.K., and E.S.W. performed computational analysis. S.J., R.K., D.Z., and E.S.W. supervised computational analysis and interpreted data. R.C. supervised scRNA-seq, J.-H.L. contributed mouse and human alveolar organoid culture methodology, S.E.C. performed pathology review of mouse lung tumors, U.K.B. constructed tissue mircoarrays and performed pathology review of human lung and lung adenocarcinoma tissue; M.J.B. contributed primary human lung tissue, T. Thomas performed intracellular iron measurements, Y. Z. and T. P. designed and performed pyrosequencing, P.K. contributed expertise on experiments involving the small intestine, and Y.S.F. provided expertise in and performed epigenomics analysis. M.J.B., E.S.W., and T. Tammela obtained funding. All of the authors approved the final manuscript.

## Materials and Methods

### Mice

All animal studies were approved by the MSKCC Institutional Animal Care and Use Committee (protocol # 17-11-008). MSKCC guidelines for the proper and humane use of animals in biomedical research were followed. All genetically engineered mice were maintained on a mixed C57BL/6 x Sv129 background. Similar numbers of female and male aged and young mice of the following genetically engineered strains were used: *Kras^LSL-G12D^*; *Trp53^flox/flox^* (*KP*)^7,8^, with or without reporter alleles including *Rosa26^LSL-Tomato^* (*T*)^58^, *Rosa^mTmG^* (*mTmG*)^59^, *Rosa26^LSL-rtTA-IRES-mKate2^* (*RIK*)^60^, *Rosa26^LSL-Luciferase^* (*L*)^61^, and *R26^LSL-Cas9-2a-eGFP^* (*Cas9*)^33^. Wild-type C57BL/6 mice were purchased from Jackson Laboratories (strain #000664) at 10 and 77-78 weeks old and housed at MSKCC’s Research Animal Resource Center until they reached 14 (young) or 104-130 (aged) weeks of age. All mice were housed in non-pathogenic conditions, controlled temperature (20– 25 °C), and a 12-hour light–dark cycle with 30-70% relative humidity. Food and water were provided *ad libitum*. Mice were genotyped at ∼2 weeks of age.

### Genetically engineered mouse lung cancer models

Lung tumors were initiated in *KP, KPT*, *KPL*, and *KP-RIK* mice by intratracheal administration of 1.5-5 x 10^8^ plaque-forming units (pfu) of AdSPC-Cre (University of Iowa, #Berns-1168) as previously described^26^. In the survival experiment (**Extended Data Fig. 1a**), mice were euthanized upon reaching humane endpoint (respiratory distress, extreme weight loss, or lethargy). *Eml4-Alk* gene fusion-driven lung tumors were induced in young and aged wild type C57BL/6 mice by intratracheal administration of 5 x 10^8^ pfu of adenoviral vectors encoding U6-sgAlk-U6-sgEml4-CMV-Cas9, as previously described^27^. To evaluate the impact of *Nupr1* knockout on tumorigenesis *in vivo*, 1.5 x 10^5^ transduction units of lentivirus were introduced to aged and young *KP-Cas9* mice by intratracheal administration. Liproxstatin-1 (Selleck Chemicals, #S7699) and solvent control [1% DMSO + 3% Kolliphor EL (Sigma-Aldrich, #C5135) + 96% PBS] was administered by intraperitoneal injection at 10 mg/kg/day for 4 weeks starting on the day of tumor initiation.

### Hyperoxia lung injury

Alveolar injury was induced in aged and young C57BL/6 mice by a 66 hours exposure to >85% O_2_ in a hyperoxia chamber (BioSpherix). The fractional inspired oxygen concentration in the chamber was monitored by an in-line oxygen analyzer and maintained with a constant flow of gas (∼3 L/min). Mice were euthanized on days 0, 3, 10 and 28 after the 66-hour exposure, followed by collection of lungs for histological analysis. Proliferating AT2 cells were identified by immunofluorescence for the AT2 marker surfactant protein-C (SPC, EMD Millipore, #AB3786) and proliferation maker Ki67 (Invitrogen, #14-5698-82) on day 3 after the hyperoxia injury. For lineage-tracing assay, young and aged *Rosa26^LSL-tdTomato^* mice were transduced by two doses of 5 x 10^9^ AdSPC-Cre (University of Iowa, #Berns-1168) and exposed to 66-hour hyperoxia 7 days after transduction. Mice were euthanized 28 days after the injury and identity of traced cells were identified by immunofluorescent staining of podoplanin (PDPN, Thermo Fisher Scientific, Cat #14-5381-82, AT1 cell marker) and surfactant protein-C (SPC, EMD Millipore, #AB3786, AT2 marker). To investigate the role of transferrin, two doses of transferrin (Rockland, #010-0134, 150 mg/kg) or PBS control were administered by intratracheal intubation at 24-hour intervals before hyperoxia exposure. Additional four doses of transferrin and PBS control were administered by intraperitoneal injection daily after the hyperoxia exposure. To evaluate the role of *Nupr1* in the proliferation and differentiation of AT2 cells *in vivo*, 1.5 x10^6^ transduction units of lentivirus (PGK-Cre-PGK-GFP-sh*Renilla*, sh*Nupr1*.1, or sh*Nupr1*.2) were introduced by intratracheal administration. Mice were exposed to hyperoxia-induced lung injury 7 days after transduction. Proliferation and differentiation of AT2 cells was evaluated at 3 and 28 days after the completion of hyperoxia exposure, respectively.

### Lung tissue harvest

Mice were euthanized by CO_2_ asphyxiation followed by systemic perfusion with PBS to clear lungs of blood. For histological analysis, lung tissues were fixed in 10% neutral-buffered formalin (Sigma Aldrich, #HT501128) overnight and further processed and embedded in paraffin or Tissue-Tek O.C.T. compound (Sakura, #4583) using standard protocols for formalin-fixed paraffin-embedded (FFPE) sections or cryosections, respectively. Five micrometer FFPE sections were prepared by the Molecular Cytology Core Facility (MCCF) at MSKCC for hematoxylin and eosin (HE) staining, immunofluorescence, and immunohistochemistry.

### Quantification of tumor burden

Tumor burden was quantified using HE or immunohistochemical detection in 5 µm FFPE sections or cryosections of lung tissues. For early-stage tumors (2-, 4- and 8-week post-tumor initiation), tumor cells were identified using a detection of Cre-inducible tdTomato fluorescence or immunohistochemical staining for the tdTomato reporter (anti-RFP, Rockland, #600-401-379). Late-stage tumors (12- and 17-week post-tumor initiation) were identified by HE staining. Tumor burden was defined by the number of nodules in the section of tumor-bearing lungs, normalized by the size of lung tissue. Tumor area and tissue area were determined using Caseviewer (3DHISTECH).

### Histological analysis

Histopathological grading of mouse lung tumors was performed on the HE-stained sections by two methods: (i) an automated deep neural network (Aiforia Technologies, NSCLC_v25 algorithm)^62^ and (ii) an independent classification by a board-certified veterinary pathologist, Dr. Sebastian E. Carrasco, DVM, PhD at our institute. Dr. Carrasco was blinded to the sample group identifiers. Established histopathological criteria to evaluate mouse models of lung cancer were used^63^. Tumor grades in pulmonary lobes ranged from 1 to 4 with grade 1 being composed by uniform histomorphology of the neoplastic cells without nuclear atypia and grade 4 being composed of pleomorphic neoplastic cells exhibiting increased nuclear atypia, including abnormal mitotic figures and hyperchromatism, and/or frequent binucleated or multinucleated cells. Alveolar size was measured from HE-stained lung sections by mean cord length using a standardized protocol^59^.

### Immunohistochemistry

Immunohistochemistry was performed on 5 μm FFPE sections using standard staining protocols. Briefly, sections were de-paraffinized and heat-induced antigen retrieval was performed using EDTA antigen retrieval buffer (Sigma Aldrich, #E1161). Sections were blocked by BLOXALL solution (Vector laboratories, #SP-6000-100) at room temperature for 30 minutes and incubated with primary antibody at 4 °C overnight. IgG controls (Thermo Fisher Scientific, #02-6102, #02-6202 and #10400C) from the corresponding species of primary antibodies were used as negative controls. Signal development was performed by ImmPRESS Polymer Detection Kits (Vector Laboratories, #MP-7401-50, #MP-7405-50 and #MP-7802-15) and ImmPACT DAB Substrate Kit, Peroxidase (HRP) (Vector Laboratories, #SK-4105) following the manufacturer’s protocol. The sections were counterstained with hematoxylin (Thermo Fischer Scientific, #72404) and mounted with coverslips. Catalog numbers and dilutions for all antibodies are available in **Extended Data Table 18**. Mounted slides were digitally scanned by the Mirax Midi-Scanner (Carl Zeiss AG). Image analysis was performed using Fiji^64^.

### Immunofluorescence

Immunofluorescence was performed on 5 µm FFPE sections or 7 µm cryosections. FFPE sections de-paraffinized and heat-induced antigen retrieval was performed using EDTA antigen retrieval buffer (Sigma Aldrich, #E1161). For cryosections, slides were air-dried for 1 hour at room temperature and fixed by acetone at −20°C for 15 minutes. Sections were blocked in donkey immunomix [0.2% BSA (Sigma, #810533), 5% donkey serum (Thermo Fisher Scientific, #31874), and 0.3% Triton-X (Fisher Scientific, #BP151-100) in PBS (Gibco, #10010-023)] at room temperature for 30 minutes. Incubation of primary antibodies diluted in donkey immunomix was performed at 4 °C overnight. Non-specific IgG (Thermo Fisher Scientific, #02-6102, #02-6202, and #10400C) from the species corresponding to the primary antibodies were used as negative controls. AlexaFluor-conjugated secondary antibodies raised in donkey were used for signal detection (Thermo Fisher Scientific #A31571, #A21207, #A32795, #A11058, #A32787). Slides were counterstained with 1 µg/mL DAPI (Sigma Aldrich, #D9542) for 10 minutes and mounted with coverslips using Mowiol mounting reagent (EMD Millipore, #475904). Catalog numbers and dilutions for all antibodies are available in **Extended Data Table 18.** Mounted slides were digitally scanned using the Mirax Midi-Scanner (Carl Zeiss AG). Image analysis was performed using Fiji^64^.

### Detection of senescence-associated β-galactosidase activity

Fluorescent senescence-associated (SA)-β-gal labeling was performed according to manufacturer’s protocol using the ImaGene Red C_12_RG lacZ Gene Expression Kit (Invitrogen, #I2906). The tissues used for staining were flash frozen in liquid nitrogen and directly embedded in Tissue-Tek O.C.T compound (Sakura, #4583). Cryoblocks were immediately sectioned at 5 µm thickness. Sections were placed on SuperFrost microscope slides (Fischer Scientific, #12-550-15) and immediately used for staining. Sections were incubated with 1% chloroquine at 37 °C for 30 minutes. Chloroquine was removed in two washes using pre-cooled PBS. Sections were incubated with C_12_RG substrate (1:50 dilution) at 37 °C for 2 hours, followed by an immediate rinse by PBS and incubation in PETG (1:50) solution at room temperature for 20 minutes. Sections were fixed in 4% PFA for 20 minutes on ice. To identify AT2 cells and tumor cells derived from AT2 cells, we performed SPC immunofluorescence staining after C_12_RG staining, as described above. Slides were digitally scanned using the Mirax Midi-Scanner (Carl Zeiss AG) and C_12_RG positive senescent cells were quantified using Fiji^64^.

### Single-molecule mRNA *in situ* hybridization

*In situ* hybridization was performed on freshly sectioned 5 µm FFPE sections using the RNAscope Multiplex Fluorescent Reagent Kit v2 (Advanced Cell Diagnostics, #323100) following the manufacturer’s protocol. Probes detecting murine *Sftpc* (Advanced Cell Diagnostics, #314101-C2), *Nupr1* (Advanced Cell Diagnostics, #434811), *Lgr5* (Advanced Cell Diagnostics, #312171-C2) and *Lcn2* (Advanced Cell Diagnostics, #313971-C3) were used. Probe signals were developed using Opal-520, Opal-570 and Opal-690 dyes (Akoya Biosciences, #FP1487001KT, #FP1488001KT and #FP1497001KT). Slides were digitally scanned by Mirax Midi-Scanner (Carl Zeiss AG). AT2 and LUAD cells were identified by *Sftpc* positivity and *Nupr1* expression was quantified using Fiji^53^ as number of individual dots per each *Sftpc^+^*cell. Intestinal stem cells were identified by *Lgr5* positivity. *Nupr1* expression was quantified using the staining intensity due to the high expression/signal level of *Nupr1* and *Lcn2* expression was quantified as number of individual dots in *Lgr5^+^* cells using Fiji^64^.

### Lung dissociation

For isolation of normal AT2 cells and LUAD cells at 4, 8 and 12 weeks post-tumor initiation, lungs were inflated with 3 ml digestion buffer [S-MEM (Thermo Fisher Scientific, #11380037) with 1.7 IU/ml dispase (Corning, #354235), 0.5 UI/µl collagenase IV (Thermo Fisher Scientific, #17104019) and 10 µg/ml DNase I (StemCell Technologies, #07900)] through the trachea and then finely minced. For the 17-week time point, tumors were micro-dissected, finely minced, and suspended in ∼2-3 volumes of digestion buffer. Tissues were dissociated in a 37 °C oven with gentle agitation for 45-60 minutes. The dissociated tissue cells were filtered using a 100 μm strainer (Corning, #431752) and centrifuged at 1500 rpm for 5 minutes at 4 °C. The supernatant was removed and red blood cell lysis was performed using BD PharmLyse (BD Biosciences, #555899) following the manufacturer’s protocol. Cells were washed with S-MEM (Thermo Fisher Scientific, #11380037) with 2% heat-inactivated fetal bovine serum (HI-FBS, Hyclone, #SH30910.03), filtered through a 40 µm strainer (Corning, #431750), and pelleted by centrifuging at 1500 rpm for 5 minutes at 4 °C. Cell pellets were resuspended in 2% HI-FBS S-MEM for staining with fluorescent antibodies.

Human lung tissues were obtained from 6 patients undergoing pulmonary resections for lung cancer at MSKCC (**Extended Data Table 16**). Normal tissue was obtained from outside the tumor margin, as determined by intraoperative pathology examination. The human lung tissues were dissociated using a similar protocol as the mouse lungs. Briefly, the lung tissue was minced into small cubic pieces and washed several times by pre-cooled PBS. Excessive PBS was drained, and tissues were further minced and dissociated in 8-10 volumes of digestion buffer for 1-1.5 hours. Dissociated tissues were processed as described above and cell pellets were resuspended in 2% HI-FBS S-MEM for staining with fluorescent antibodies.

### Fluorescence activated cell sorting (FACS)

Single cell solutions of lung cells were incubated with mouse FcR block (BD Biosciences, #553142) on ice for 10 minutes, followed by 30 minutes of staining with desired antibody panels on ice. To sort cells for single-cell sequencing, cell-hashing method was used to maximize the number of biological replicates and control for batch effect in each single-cell sequencing lane. To do this, the cell suspension derived from each mouse was labeled by a different hashtag antibody (Biolegend, TotalSeq Panel B, #B0301-B0310, 2 µl/test) together with the desired antibody panel, following the manufacturer’s protocol. After the 30-minute staining period, cells were centrifuged at 1500 rpm for 5 minutes at 4 °C and washed twice with 2% HI-FBS S-MEM. Cell pellets were resuspended in PBS with 2% HI-FBS (Hyclone, #SH30910.03) containing 1 µg/mL DAPI (Sigma Aldrich, #D9542) to detect dead cells. Samples were sorted using a BD FACS Aria Sorter using 4-way purity mode. AT2 cells were defined as MHCII^+^/EpCAM^+^/Sca1^-^/podoplanin^-^/lineage^-^ (CD45, CD31, CD11b, CD11c, F4/80 and Ter119) ^-^/DAPI^-^ cells. Sca1 and podoplanin (PDPN) were included to maximize the purity of AT2 cells. Early-stage cancer cells (4, 8 and 12 weeks post-tumor initiation) were isolated by selecting for tdTomato^+^ (for *KPT* mice), mKate2^+^ (for *KP-RIK* mice), or GFP^+^ (for *KP-Cas9* mice) and EpCAM^+^/lineage^-^/DAPI^-^ cells. For late-stage tumors, EpCAM^+^/lineage^-^/DAPI^-^ cancer cells were isolated from micro-dissected tumors.

For the isolation of human AT2 cells, a single-cell solution of lung cells was incubated with Human TruStain FcX (Biolegend, #422302, 1:20) for 10 minutes, followed by staining with anti-human CD45 (Biolegend, #368538), CD31 (Biolegend, #303116), EpCAM (Biolegend, #324222), and HTII-280 (Terrace Biotech, #TB-27AHT2-280, 1:40)^65^ for 30 minutes on ice. Cells were washed once with 2% HI-FBS S-MEM and further stained with PE-conjugated anti-mouse IgM (Thermo Fisher Scientific, #12-5790-81) for 30 minutes to detect the HTII-280 primary antibody. Cell samples were centrifuged at 1500 rpm for 5 minutes at 4 °C and washed twice with 2% HI-FBS S-MEM. Cell pellets were resuspended in 2% HI-FBS PBS with 1 µg/mL DAPI (Sigma Aldrich, #D9542) to stain dead cells. Samples were sorted using a BD FACSAria Sorter on purity mode. AT2 cells were defined as HTII-280^+^/EpCAM^+^/CD45^-^/CD31^-^/DAPI^-^ (ref.^65^). All antibodies used for flow cytometry are listed in **Extended Data Table 18**.

### Heterochronic transplantation

LUAD cells were isolated from micro-dissected autochthonous *KP* lung tumors at 18 weeks post-tumor induction by FACS as described above. Tumor cells were pelleted, resuspended in 100 μ1 S-MEM and transplanted to sex-matched, syngeneic 10 (young) or 46-54 week-old (middle-aged) B6129SF1/J recipient mice purchased from Jackson Laboratories (strain #101043). Lung tissues from recipient mice were harvested 13 weeks after transplantation for tumor burden quantification.

### Alveolar organoid culture

FACS-purified AT2 cells were co-cultured with primary mouse pulmonary endothelial cells to generate alveolar organoids, as before^30^. Endothelial cells (CD31^+^/CD45^-^/DAPI^-^) were isolated from 4-week-old *Rosa26^mTmG^* mice^59^ by FACS. Primary endothelial cells were expanded briefly using endothelial cell media [Advanced DMEM (Thermo Fisher Scientific, #12491015), 20% FBS (Hyclone, #SH30910.03), 1% GlutaMax (Thermo Fisher Scientific, #35050061), 1% Pen/Strep (Thermo Fisher Scientific, #15070063), endothelial cell growth supplement (ECGS, Sigma Aldrich, #E2759, 0.1 mg/ml), heparin (Sigma Aldrich, #H3149, 0.1mg/ml), and 25 mM HEPES (Thermo Fisher Scientific, #15630080)]; only cells from passages 2-3 were used. One to five thousand freshly sorted AT2 cells with 50,000 endothelial cells were resuspended in 50 µl alveolar organoid culture media [Ham’s F-12 (Thermo Fisher Scientific, #11765047), 10% FBS (Hyclone, #SH30910.03), 1% GlutaMax (Thermo Fisher Scientific, #35050061), 1% Pen/Strep (Thermo Fisher Scientific, #15070063), 1% ITS (Millipore Sigma, #I3126) and 1% HEPES (Thermo Fisher Scientific, #15630080)] and mixed with 50 µl Matrigel (Fisher Scientific, #CB-40230C). Cell mix was placed in cell culture inserts (Thermo Fisher Scientific, #08-770). Alveolar organoid culture media (500 µl) was added to the reservoir outside the insert (Thermo Fischer Scientific, #353504) and replaced every 3 days during culture. ZZW-115 (Cayman Chemical Company, #34974, 2 µM), deferoxamine mesylate (DFO, Selleck Chemicals, #S5742, 2 µM), or iron-loaded mouse transferrin (Rockland, #010-0134, 50 µg/ml) were added to the media on day 3 and refreshed every 3 days. Organoids were imaged with an EVOS M5000 microscope and the number of organoids > 50 µm in diameter was counted at 3 weeks after plating. For secondary organoid cultures, primary organoid culture was digested with 5 UI/ml dispase (Corning, #354235) for 1 hour at 37 °C. Organoids released from the Matrigel were collected by centrifuging at 1000 rpm for 5 minutes. Organoids were further dissociated into single cell solution with TrypLE (Thermo Fisher Scientific, #12604013) for 10 minutes at 37 °C. Live AT2 cells were purified from single-cell solution by sorting for MHCII^+^/EpCAM^+^/DAPI^-^ and re-plated using the same protocol as for primary AT2 culture.

### Ex vivo transformation assay

AT2 cells isolated from *KP-RIK* and *KP-Cas9* mice were transduced by lentivirus at multiplicity of infection (MOI) of 10 by spinfection (600 g, 37 °C, 30 minutes). One to five thousand AT2 cells were plated in the inserts with endothelial cells, as described above. Alveolar organoid culture media was used and replaced every 3 days. ZZW-115 (Cayman Chemical Company, #34974, 2 µM), deferoxamine mesylate (DFO, Selleck Chemicals, #S5742, 2 µM), GSK3685032 (Selleck Chemicals, #E1046, 1 µM), iron-loaded recombinant mouse transferrin (Rockland, #010-0134, 50 µg/ml), recombinant mouse lipocalin-2 (R&D Systems, #1857-LC-50, 100 ng/ml), liproxstatin-1 (Selleck Chemicals, #S7699, 5 μM), RSL-3 (Selleck Chemicals, #S8155, 0.2 μM), erastin (Selleck Chemicals, #S7242, 5 μM), or ferric ammonium citrate (FAC) (Sigma-Aldrich, #F5879, 25 μM) were added to the media on day 3 and refreshed every 3 days. Transformed tumor spheres were identified by mKate2 or GFP fluorescence. Organoids were imaged with an EVOS M5000 microscope and tumor spheres > 50 µm in size were counted 2 weeks after plating. Short hairpin and single guide RNA sequences listed in **Extended Data Table 13** were introduced into pLenti-TRONO-PGK-Cre-PGK-GFP-shRNA or pLenti-TRONO-PGK-Cre-U6-sgRNA constructs. To overexpress *Lcn2*, cDNA sequence of *Lcn2* was linked to Cre recombinase by a P2A sequence^66^ (pLenti-PGK::Cre-P2A-Lcn2-U6::sgNupr1/sgControl).

### Tumor sphere culture

For serial passage of *ex vivo* transformed tumor spheres, primary tumor spheres in Matrigel were digested with 5 UI/ml dispase (Corning, #354235) for 1 hour at 37 °C. Organoids released from Matrigel were collected by centrifuging at 1000 rpm for 5 minutes. Organoids were further dissociated into single-cell solution with TrypLE (Thermo Fisher Scientific, #12604013) for 10 minutes at 37 °C. Cells were re-plated in Matrigel plugs at 3000 cells/plug as before^23^ and were grown in tumor sphere media [Advanced DMEM/F-12 (Thermo Fisher Scientific, #12634028) with 2% HI-FBS (Hyclone, #SH30910.03), 1% GlutaMax (Thermo Fisher Scientific, #35050061), 1% Pen/Strep (Thermo Fisher Scientific, #15070063), 1% HEPES (Thermo Fisher Scientific, #15630080), 10 ug/mL gentamicin (Thermo Fisher Scientific, 15750060)] without endothelial cells. Cells were grown for eight passages. Following each passage, a portion of cells were preserved in RLT Plus buffer (Qiagen, #1053393) with 1% 2-mercaptoethanol (Sigma Aldrich, #M6250) and stored at –80 °C for RNA extraction. RNA was extracted with the RNeasy Mini Kit (Qiagen, #74104) following the manufacturer’s protocol.

### Human alveolar organoid culture

Human alveolar organoids were established from primary AT2 cells isolated from human lungs using the insert culture system. One to four thousand AT2 cells were resuspended in 50 µl complete human alveolar organoid media [Advanced DMEM (Thermo Fisher Scientific, #12491015), 1% GlutaMax (Thermo Fisher Scientific, #35050061), 1% Pen/Strep (Thermo Fisher Scientific, #15070063), 1% HEPES (Thermo Fisher Scientific, #15630080), B27 supplement (Thermo Fisher Scientific, #17504044, 1:50), recombinant human FGF7 (Peprotech, #100-19, 100 ng/ml), recombinant human FGF10 (Peprotech, #100-26, 100 ng/ml), recombinant human Noggin (Peprotech, #120-10C, 100 ng/ml), recombinant human EGF (Thermo Fisher Scientific, #PHG0311, 50 ng/ml), N-acetylcysteine (Sigma Aldrich, #A9165, 1 mM), Nicotinamide (Sigma Aldrich, #N0636, 10 mM), A083-01 (Selleck Chemicals, #S7692, 1 µM), CHIR99021 (Sigma Aldrich, #SML1046, 3 µM), and Rspo3-Fc Fusion Protein Conditioned Medium (Immunoprecise, #R001-100ml, 2%)] and mixed with 50 µl Matrigel (Fisher Scientific, #CB-40230C). Human alveolar organoid media was refreshed every 3 days until the end of the experiment. Iron-loaded recombinant human transferrin (Optiferrin Recombinant Transferrin, InVitria, #NC9954311, 100 µg/ml) was added to the media on day 3 and refreshed every 3 days. Organoids were imaged using an EVOS M5000 microscope and organoids >30 µm in size were counted 3 weeks after plating.

### Intracellular iron measurement

Total cellular iron was measured by induction-coupled plasma mass spectrometry (ICP-MS; Agilent 7900) equipped with integrated sample introduction system (ISIS3) in high energy helium mode (10 ml/min; 1 point/peak, 4 replicates, 25 sweeps/replicate). The method was linear between 5-1000 ng/mL. Cell pellets (16,000-10,000,000 cells) were digested overnight in 100 µl of tetramethylammonium hydroxide. The cell pellet digestates were diluted 1:80 in diluent (4% 1-butanol, 1% TMAH, 0.01% Triton X-100, 0.01% ammonium pyrrolidinedithiocarbamate) and quantified using the iron isotope 56 relative to a 7-point calibration and germanium as an internal standard. Results are reported relative to cell number.

Labile iron was measured by staining of FerroFarRed dye and quantified by flow cytometry. Cancer cells arising from AT2 cell *ex vivo* transformation were isolated and dissociated into single-cell solutions as described above. Cells were stained by BioTracker Far-red Labile Iron Dye (Sigma-Aldrich, # SCT037, 5 μM) and EpCAM antibody at 37°C for 1 hour, washed twice by HBSS and resuspended in HBSS with 1 µg/mL DAPI (Sigma Aldrich, #D9542) to detect dead cells. Tumor cells were identified by EpCAM positivity and labile iron level was determined by the median fluorescence intensity. Data were collected and analyzed by a BD LSRFortessa Cell Analyzer and FlowJo software, respectively.

### Lentivirus production

HEK293T cells were transfected with a shuttle vector plasmid and packaging plasmids pMD2.G (Addgene, #12259) and psPAX (Addgene, #12260) using TransIT-LT1 Transfection Reagent (Mirus Bio, #MIR 2305). Virus-containing media was collected 48 and 72 h post-transfection, concentrated by ultracentrifugation (31500 rpm, 4 °C, 2 hours), resuspended overnight at 4 °C in D-MEM and stored at −80 °C. Lentiviral vectors encoding Cre recombinase were titered using GreenGo Cre recombination reporter cells, as before^24^.

### Measurement of transduction efficiency

Adenoviral transduction efficiency of AT2 cells *in vivo* was measured following intratracheal delivery of 5 x 10^8^ pfu AdSPC-Cre (University of Iowa Viral Vector Core, # Berns-1168) to lungs of aged and young *Rosa26^mTmG^* mice^55^. Transduced AT2 cells were identified by flow cytometry as GFP^+^/MHCII^+^/EpCAM^+^/lineage^-^/DAPI^-^. Alternatively, cryosections of lungs from aged and young *Rosa26^LSL-tdTomato^*mice infected as described above were used to quantify tdTomato^+^ AT2 cells as an additional method to quantify transduction efficiency *in situ*. Lentiviral transduction efficiency was measured using a similar flow cytometry approach. Lentivirus harboring phosphoglycerate kinase-1 promoter-driven Cre (PGK-Cre) was prepared and titered in-house, as before^25,56^. Fifty thousand transduction units of PGK-Cre lentivirus were intratracheally delivered to lungs of *Kras^+/+^;Trp53^flox/flox^; rtTA-IRES-mKate2 (P-RIK)* mice, which were littermates of the *Kras^LSL-G12D/+^; Trp53^flox/flox^;rtTA-IRES-mKate2 (KP-RIK)* mice used in the lung tumor studies. Lung tissues were dissociated and the total number of mKate2 positive AT2 cells (mKate2^+^/EpCAM^+^/MHCII^+^/lineage^-^/DAPI^-^) was calculated by flow cytometry to measure transduction efficiency.

To measure the transduction efficiency of AT2 cells in the *ex vivo* transformation assay, PGK-GFP lentivirus, produced and titered in-house, was introduced to isolated wild-type aged vs. young primary AT2 cells by spinfection. The transduction efficiency was measured as a ratio of GFP positive organoids in the total pool of alveolar organoids. High multiplicity of infection (MOI, 10∼20) was used in *ex vivo* transformation assays to ensure high (∼100%) transduction efficiency.

### Pyrosequencing

Genomic DNA from cultured *KP* tumor cells, transduced with lentiviral control shRNA (shRenilla) or shRNAs targeting *Dnmt1* or treated with DNMT1 inhibitor (GSK3685032, Selleck Chemicals, #E1046, 5 µM), was purified using DNeasy Blood & Tissue Kit (Qiagen, #69506). Five hundred nanograms of each DNA sample and control DNA were bisulfite-treated (BST) using the EZ DNA Methylation-Lightning Kit (Zymo Research, # D5030) following the manufacturer’s protocol. Amplification and sequencing primers for pyrosequencing analysis are listed in **Extended Data Table 14** and were designed using PyroMark Assay Design Software (Qiagen). PCR reaction consisted of 1.5 µl of BST samples or control DNA, 1x PyroMark PCR Master Mix (Qiagen, #978703), 1x CoralLoad Concentrate (Qiagen, #201203), 1x EvaGreen (Biotium, #31000), and 0.2 µM of a forward primer and a reverse primer. Cycling conditions were 95 °C for 15 mins, 45 cycles of [94 °C for 30 s, specific primer annealing temperature for 30 s and 72 °C for 30 s], followed by a final extension at 72 °C for 10 mins. PCR amplification was confirmed by evaluation of cycle of quantification (Cq) values and confirmation of melt curve profiles derived from high-resolution melt (HRM) analysis. HRM was performed immediately after amplification with a temperature range of 60 °C to 90 °C at 0.2 °C increments. PCR was conducted using a CFX 96 Touch Real-time PCR Detection System (Bio-Rad) and analyzed using the accompanying software CFX Manager. The biotinylated PCR product was purified and subjected to pyrosequencing using the PyroMark Q24 System (Qiagen), using forward sequencing primers according to the manufacturer’s protocol. Sequencing results were reported as the percentage of cytosine vs. thymine at each CpG site.

### Quantitative PCR (qPCR)

RNA was extracted from sorted cancer and AT2 cells using the RNeasy Micro Kit (Qiagen, #74004) or TRIzol reagent (Invitrogen, #15596026) by standard chloroform-isopropanol precipitation. cDNA was synthesized using the PrimeScript RT Reagent Kit (Clontech, #RR037B). Quantitative PCR was performed in triplicate with 30 ng of cDNA using the Powerup SYBR Green Master Mix (Applied Biosystems, #A25778) on the QuantStudio 7 Flex Real-Time PCR System. The ΔΔCT method was used to compare markers of interest and expression was normalized to the housekeeping gene *Gapdh*. The primer sequences used in this study are listed in **Extended Data Table 13**.

To quantify the efficiency of *Eml4-Alk* gene fusion formation, genomic DNA was extracted from FACS-sorted AT2 cells of both aged and young C57BL/6 mice 5 days after intra-tracheal intubation with the virus. *Eml4-Alk* fusion-specific primers, listed in **Extended Data Table 13**, were used to detect the fusion gene and miR17-92 locus was used as control^27^.

### Single-cell mRNA sequencing (scRNA-seq)

Wild-type AT2 cells and *KP* LUAD tumor cells (4, 12, and 17 weeks post-tumor initiation) were isolated by FACS as described above. LUAD cells from tumors at 19-20 weeks post-initiation, representing all mature cancer cell states, were added to the analysis to facilitate unsupervised clustering and cell state identification. Ten to forty thousand freshly sorted cells were suspended in PBS containing 0.04% BSA at 1000 cells/µl for droplet-based scRNA-seq. Encapsulation and library preparation were performed using the 10X Genomics Chromium Single Cell 3’ Library & Gel bead Kit V3 (10X Genomics, PN-1000121) according to the manufacturer’s protocol. Libraries were sequenced using the Nova-Seq 6000 platform (Illumina).

### Bulk mRNA sequencing of AT2 cells following DNMT1 inhibition *in vivo*

DNMT1 inhibitor GSK3685032 (Selleck Chemicals, #E1046) or vehicle [10% captisol (sulfobutylether-β-cyclodextrin, MedChemExpress, #HY-17031) adjusted to pH 4.5–5, stored for up to 1 week at 4 °C] was administered by intraperitoneal injection, twice daily, at the dose of 10 mg/kg body weight. After an eight-day treatment, AT2 cells were isolated from all mice by FACS, as described above. Cells were pelleted and resuspended in 350 µl Buffer RLT plus (Qiagen, #1053393) and stored at –80 °C. RNA extraction was performed using RNeasy Mini Kit (Qiagen, #217004) on the QIAcube Connect (Qiagen) according to the manufacturer’s protocol. Samples were eluted in 35 µL RNase-free water. After RiboGreen quantification and quality control by Agilent BioAnalyzer, 2 ng total RNA with RNA integrity numbers ranging from 9.0 to 9.7 underwent amplification using the SMART-Seq v4 Ultra Low Input RNA Kit (Clonetech, #63488), with 12 cycles of amplification. Subsequently, 10 ng of amplified cDNA was used to prepare libraries with the KAPA Hyper Prep Kit (Kapa Biosystems, #KK8504) using 8 cycles of PCR. Samples were barcoded and run on a NovaSeq 6000 in a PE100 run, using the NovaSeq 6000 S2 Reagent Kit (200 Cycles) (Illumina). An average of 34 million paired reads were generated per sample and the percentage of mRNA bases per sample ranged from 89% to 93%.

### Genome-wide methylome sequencing of AT2 cells and tumors from aged and young mice

Fifty to one hundred thousand wild-type mouse AT2 cells or LUAD tumor cells (12 weeks post-tumor initiation) were isolated by FACS, as described above. Genomic DNA was isolated using the DNeasy Blood & Tissue Kit (Qiagen, #69504). NEB Next Enzymatic Methyl-seq (EM-seq, New England Biolabs, #E7120S) was used to identify 5-methylcytosine and 5-hydroxymethylcytosine bases and sequencing libraries were constructed by NEBNext Ultra II DNA Library Prep Kit for Illumina (New England Biolabs, #E7645S), following the manufacturer’s protocol. After PicoGreen quantification and quality control by Agilent TapeStation, libraries were pooled equimolar and run on a NovaSeq 6000 at PE150, using the NovaSeq 6000 S4 Reagent Kit (300 cycles) (Illumina). The loading concentration was 0.6-0.7 nM and a 1% spike-in of PhiX was added to the run to increase diversity and for quality control purposes. The conversion efficiency of 5-methylcytosine and 5-hydroxymethylcytosine bases was over 98% (**Extended Data Table 20**). The runs yielded on average 795 million reads per library.

### Perturbation of Nupr1 in vivo using CRISPR/Cas9

Lungs of aged and young *KP*-*Cas9* mice were intratracheally transduced with U6-sgRNA-PGK-Cre lentivirus containing a guide RNA targeting *Nupr1* or a non-targeting control sgRNA^25^. Lungs with tumors were harvested at 10 weeks post-tumor initiation. Whole lungs were paraffin-embedded and stained for GFP (Abcam, #ab5450) and Ki67 (Thermo Fisher Scientific, #14-5698-82) to quantify the tumor burden and percentage of proliferating tumor cells, respectively. Additionally, some lung tumors were micro-dissected and dissociated for isolation of GFP^+^/EpCAM^+^/Lineage^-^/DAPI^-^ tumor cells for bulk mRNA sequencing.

### Perturbation of Nupr1 and Dnmt1 in vivo using shRNAs

Lentivirus harboring shRNAs targeting murine *Nupr1* or *Dnmt1* (pLenti-TRONO-PGK-Cre-P2A-GFP-shRNA-pA) were prepared and titered in-house as described above. One to two million transduction units of lentivirus was administered to both aged and young *Rosa26^mTmG^* mice by intratracheal intubation. Transduced cells were marked by the transition of endogenous fluorescence from membrane-bound tdTomato (mT) to membrane-bound GFP (mG). For *shDnmt1*, mice were harvested 12-14 days post-viral infection and live GFP^+^/EpCAM^+^ cells were isolated by FACS, followed by single-cell mRNA sequencing. For *shNupr1*, mice were exposed to hyperoxia-induced lung injury 7 days post-viral infection. Lung tissues were harvested 3 and 28 days after hyperoxia exposure for the evaluation of proliferation and differentiation of AT2 cells, respectively.

### Arrayed CRISPR interference screen to identify *Nupr1* distal enhancers

Enzymatically dead Cas9-KRAB fusion protein (Lenti-dCas9-KRAB-blast, Addgene, #89567) was introduced into three *KP* LUAD cell lines by lentiviral transduction and blasticidin selection. Three sgRNAs targeting each candidate enhancer were cloned into LentiGuide-Puro backbone (LentiGuide-Puro, Addgene, #52963). A non-targeting sgRNA and sgRNAs that target the *Nupr1* gene body were used as negative and positive controls, respectively. The Cas9-KRAB cell lines were transduced by the array of sgRNAs and selected with puromycin. Cells were harvested 10 days post puromycin selection and qPCR was performed to evaluate the expression of *Nupr1*. SgRNA sequences are listed in **Extended Data Table 11**.

### Intestinal organoid culture

Small intestines from both young and aged C57BL/6 mice were isolated, thoroughly flushed with PBS, opened, and cut into 5-8 mm pieces. These tissue pieces were moved to 50 ml conical tubes and washed by vigorous shaking in PBS 4 times until the supernatant was clear. The clean tissue fragments were transferred to 10 ml 5 mM EDTA in PBS, shaken vigorously for 10 times and kept under agitation at 4°C for 2 hours. The supernatant was replaced by 10 ml sterile PBS with 200 units of DNaseI and after 30 shakes supernatant was collected as fraction I. This step was repeated and supernatant (fraction II) was combined with fraction I. Fractions I + II were filtered through a 100 μm cell strainer and intact crypts were examined under the microscope and counted. Five hundred to one thousand crypts were seeded in each well in a 24-well plate using 50 μl crypt growth media [Advanced DMEM (Thermo Fisher Scientific, #12491015), 1% GlutaMax (Thermo Fisher Scientific, #35050061), 1% Pen/Strep (Thermo Fisher Scientific, #15070063), 1% HEPES (Thermo Fisher Scientific, #15630080), 25% L-WRN condition media (prepared in-house by collecting conditioned media from cell culture with 100% confluency for 8 days^67^, recombinant mouse Noggin (Peprotech, #250-38, 50 ng/ml), recombinant mouse EGF (Thermo Fisher Scientific, #PMG8045, 40 ng/ml), CHIR99021 (Sigma Aldrich, #SML1046, 5 µM), and Rspo3-Fc Fusion Protein Conditioned Medium (Immunoprecise, #R001-100ml, 5%)], and mixed with 50 µl Matrigel (Fisher Scientific, #CB-40230C). Media was refreshed every three days and the treatment [Liproxstatin-1 (Selleck Chemicals, #S7699, 10 μM), RSL3 (Selleck Chemicals, #S8155, 1 μM), transferrin (Rockland, #010-0134, 75 µg/ml), and DFO (Selleck Chemicals, #S5742, 0.5 µM)] started with the first media refreshment. Organoids over 50 μm in diameter were quantified 10 days after seeding.

### Human tissue

Archived and fresh human normal lung and LUAD tissues obtained under MSKCC Institutional Review Board approval (IRB #06-107 and IRB #12-245). Archived normal human lung tissue sections were obtained from both young and middle-aged (< 50 years old) vs. aged (>70 years old) patients. Areas chosen for analysis were evaluated by Dr. Umeshkumar Bhanot, MD, PhD, a board-certified pathologist at MSKCC to contain normal healthy lung. The clinical diagnoses of these patients were included in **Extended Data Table 15**. A human LUAD tissue microarray (TMA) including 35 young and middle-aged (<55 years old) and 37 aged (> 80 years old) cases was constructed. Tissue blocks from each case were examined by Dr. Bhanot to select for regions with maximal presence of tumor tissue and three cores (2.0 mm in diameter) of each case were taken to construct the TMAs (**Extended Data Table 16**). In the analysis, one aged case was excluded because the tissue detached from the glass slide.

Fresh human lung tissue to generate alveolar organoids was obtained with informed consent from patients under protocols approved by the MSKCC Institutional Review Board (IRB #12-245). Normal lung tissue was obtained from the distal tissue of surgical resections of patients with lung adenocarcinomas. Complete lists of patients involved were included in **Extended Data Table 16**.

### Computational Methods

#### Processing and analysis of single-cell RNA-sequencing data

FASTQ files of single-cell RNA-sequencing data generated on the 10X Chromium platform were processed using the standard CellRanger pipeline (version 5.0.0). Reads were aligned to a custom GRCm38 / mm10 reference containing additional transgenes used in the study. The generated cell-gene count matrices were analyzed using a combination of published packages and custom scripts in either the scanpy / AnnData^68^ (Python) or Seurat^69^ (R) ecosystems. The specific analytic workflows employed are summarized below.

#### Pre-processing, quality control, and filtering of single-cell RNA-sequencing data

Single-cell RNA-sequencing data from young / aged *KP* LUAD tumors (4 weeks, 12 weeks, 17-18 weeks, and 19-20 weeks following initiation) and healthy lungs were de-hashed and compiled into a combined count matrix. Cells with less than 500 UMIs, more than 20% mitochondrial UMIs and low complexity based on the number of detected genes vs. number of UMIs were removed. Doublets were filtered using *scrublet*^70^. UMI counts were normalized using the size factor approach, as previously described^71^.

Single-cell transcriptomes that passed filtering were subjected to an initial round of unsupervised clustering and low-dimensional embedding. Specifically, highly variable features were selected using a variance stabilizing transformation and dimensionality reduction was performed on normalized, log2-transformed count data using either principal component analysis or non-negative matrix factorization. The dimensionality reduced count matrices were then used as input for unsupervised clustering with the *leiden* algorithm and two-dimensional embedding using UMAP; *bbknn* was used to control for batch effects. Epithelial and stromal clusters were identified using a set of previously published marker genes.

#### Clustering of cancer cells from young and aged *KP* LUAD tumors

Young and aged cells belonging to epithelial clusters (AT1 cells, AT2 cells, and cancer cells) were selected. To ensure that the downstream analysis was not biased by any particular tumor stage or age, cells were randomly subsampled to approximately equalize cell counts between tumor stages and ages, resulting in the following cell numbers:

| Model | Number of mice | Number of cells |
| --- | --- | --- |
| **Normal AT2 cells** |  |  |
| **Healthy - young** | **4** | **1922** |
| **Healthy - aged** | **4** | **2161** |
| **Lung tumors at 4 weeks PTI** |  |  |
| **4wk - young** | **4** | **3000** |
| 4wk - young - KPT | 4 | 3000 |
| **4wk - aged** | **3** | **2535** |
| 4wk - aged - KPT | 3 | 2535 |
| **Lung tumors at 12 weeks PTI** |  |  |
| **12wk - young** | **7** | **3000** |
| 12wk – young - KPT | 2 | 1064 |
| 12wk – young – KP-RIK | 5 | 1936 |
| **12wk - aged** | **5** | **3000** |
| 12wk – aged - KPT | 1 | 658 |
| 12wk – aged – KP-RIK | 2 | 1506 |
| 12wk – aged - KPL | 1 | 644 |
| 12wk – aged – KPL/RIK | 1 | 192 |
| **Lung tumors at 17-18 weeks PTI** |  |  |
| **17-18wk - young** | **8** | **3000** |
| 17-18wk – young - KP | 8 | 3000 |
| **17-18wk - aged** | **4** | **2120** |
| 17-18wk – aged - KP | 4 | 2120 |
| **Lung tumors at 19-20 weeks PTI** |  |  |
| **19-20wk - young** | **6** | **2850** |
| 19-20wk – young - KP | 4 | 2071 |
| 19-20wk – young - KPL | 2 | 779 |
| **19-20wk - aged** | **6** | **3000** |
| 19-20wk – aged - KP | 3 | 1515 |
| 19-20wk – aged - KPL | 2 | 965 |
| 19-20wk – aged - KPT | 1 | 520 |

Unsupervised clustering and UMAP embedding of cancer cells was performed as described above. Cluster identities were assigned using marker genes identified in a previous study^23^.

#### Analysis of enrichment of young or aged cells in *KP* LUAD tumors

To test whether young or aged cells were enriched in certain cancer cell populations, a meta-cell-based approach was employed. Meta-cells are neighborhoods of similar cells representing discrete cell states that are identified in high dimensional space and can subsequently be projected onto low-dimensional representations of the data. Meta-cells were identified separately for each tumor stage using the SEACells algorithm^72^ based on equal numbers of young and aged cells. To control for batch effects, *harmony*-corrected^73^ principal components were used as input for meta-cell generation. The ratio of young and aged cells in each meta-cell was determined and projected onto the UMAP.

#### Identification of age-related gene signatures

The differentially expressed gene analysis in normal AT2 and LUAD was performed using the following pipeline. The raw gene expression count was normalized using log2TPM and filtered with expression threshold >10% of cells. The differential gene expression analysis was performed with MAST^74^ (1.16.0) in a two-part generalized linear model. DEGs were identified in all tumor cells with batch, sex, cell type and cellular detection rate added as covariates. Subsequently, the aging-related DEGs in all five LUAD cell states (AT2-like, AT1-like, high-plasticity state, endoderm-like, ribosomal) that molecularly define LUAD progression were performed separately with batch, sex, and cellular detection rate included in the model as covariates. A similar DEG analysis was also carried out in the healthy AT2 cells.

The inherited aging signature across wild-type AT2 cells and the five LUAD states was defined based on the following criteria: (1) significance in wild-type AT2 cells with FDR < 0.1; (2) same gene expression change trend across all cell types; (3) top 25 upregulated and downregulated based on the sum of absolute DEG MAST regression coefficient across all cell types.

#### Gene set enrichment analysis

Age-related gene signatures (*p*-value < 0.05; fold change > 0.1; n(young) = 1193 genes; n(aged) = 739 genes) in all tumor cells were tested for gene set enrichment using the EnrichR and PreRank functionalities implemented in the *gseapy* package^75^. The following gene set repositories were used for enrichment analysis: MSigDB Hallmark (2020), KEGG mouse (2019), Reactome (2022), GO Biological Process (GOBP; 2021). In addition to unsupervised gene set enrichment analysis, iron metabolism related gene sets were *a priori* identified from the aforementioned repositories and tested on the aged signature.

#### Processing and analysis of bulk-cell RNA-sequencing data

FASTQ files of bulk-cell RNA-sequencing data were aligned to a custom GRCm38 / mm10 reference containing additional transgenes using STAR (version 2.7.5a)^76^. Read counts were generated using the Python package HTSeq-count^77^. The generated sample-gene count matrices were analyzed using a combination of published packages and custom scripts. The specific analytical workflows employed are summarized below.

#### Analysis of gene expression during *ex vivo* transformation

Bulk RNA-sequencing data from *ex vivo* transformation experiments of young and aged cells was compiled object and normalized using the size factor normalization function of *DESeq2*^78^. Highly variable features were identified using a variance stabilizing transformation. Diffusion maps were generated in *scanpy* using the top principal components from normalized, log2-transformed count data. The pseudotime position of each sample was then calculated in *scanpy* in diffusion space using untransformed AT2 cells as root. To identify genes differentially expressed between young and aged ex vivo transformed cells independent of differentiation stage, gene expression models controlling for either passage or pseudotime were fitted in *DESeq2*.

#### DNA methylation data processing

The Bismark pipeline^79^ was adopted to map DNA methylation sequencing reads and determine cytosine methylation states. Using Trim Galore (version 0.6.4)^80^, raw reads with low-quality (less than 20) and adapter sequences were removed. The trimmed sequence reads were C(G) to T(A) converted and mapped to similarly converted reference mouse genome (mm10) using default Bowtie 2^81^ settings implemented by Bismark. Duplicated reads were discarded. The remaining alignments were then used for cytosine methylation calling by Bismark methylation extractor. For AT2 cells, around 22.5M CpG sites were detected with an average coverage of 33x in the eight samples. For lung tumor cells, 23M CpG sites were recovered with an average coverage of 40x in nine samples.

#### Differential methylation analysis

Differentially methylated CpGs (DMCs) were identified using the DSS R package^82,83^ on the basis of dispersion shrinkage method followed by Wald statistical test for beta-binomial distributions. Any CpGs with FDR < 0.1 and methylation percentage difference greater than 5% were considered significant DMCs. Pairwise comparisons were conducted between the aged group and young group for normal AT2 cells and LUAD cells, respectively.

#### Correlation of DMCs with gene expression changes at regulatory regions

To connect DMCs with promoters and enhancers in AT2 cells, we mapped DMCs to modifications of histone (H)3K4me1, H3K27ac, and H3K4me3^84^. The correlation between differential methylation level at AT2 DMCs and differential gene expression change at AT2 DEG (FDR < 0.05) in each AT2 cell histone mark category was calculated using Pearson correlation test via the *cor.test* function in R.

#### Mean promoter methylation level at aging signatures in AT2 and LUAD

The mean promoter methylation level at aging signature genes were calculated by taking the mean of all measured CpG sites overlapping the promoter region. The promoter is defined as 1kb upstream and 200 bp downstream 200 to the transcriptional start site (TSS). Wilcoxon test was performed to test the statistical significance of the difference in R. The correlation between the promoter differential methylation in AT2 and LUAD were measured using Pearson correlation via the *cor.test* function in R.

#### Analysis of gene expression change following DNMT1 inhibition in AT2 and LUAD cells

The raw gene expression count data were filtered with filterByExpr() command and normalized with the TMM method within the edgeR pipeline (version 3.34.1)^85^ in R. Subsequently, the differentially expressed genes (DEGs) were identified with an overdispersed Poisson model adjusting for batch and sex using edgeR (version 3.34.1).

#### Module score method comparing mouse and human LUAD gene expression

To study the relevance of the aged and young *KP* mouse model signatures in patients, a correlative strategy leveraging the TCGA-LUAD cohort was used. Patients with at least one non-synonymous KRAS mutation and at least 50% tumor purity were selected (*n* = 120). First, significantly differentially expressed genes (DEGs) from malignant cells from the aged versus young mouse model tumors were selected. Next, human orthologs of these DEGs were obtained using the R package *biomaRt*^86^.

Using R, we transformed the FPKM data to TPM and performed quantile normalization. To focus on the effect of aging rather than microenvironment infiltration, we regressed out the tumor purity score for each sample by using linear regression to model each gene’s expression according to tumor purity and setting the residuals as the new data values. Next, we computed composite scores for the orthologs of the age-related mouse DEGs using the previously described module score method^87^. Two module scores were computed for each patient for the aged and young DEGs respectively. Next, we performed correlation analysis of each patient’s module scores with their “age at index” variable using Pearson correlation via the *cor.test* function in R. To subselect genes from the mouse models that were especially strongly correlated with age in patients, we further utilized a jackknifing approach, leaving one gene at a time out of the module and re-computing the module score’s correlation with age. If the module score’s correlation with age was strengthened when leaving out a gene, then that gene was discarded. Pearson correlation of the jackknife-filtered aged and young module scores was reported at the end.

In addition to correlation, a multivariable regression approach was utilized to study the independent association of the aforementioned module scores with patient age in TCGA. Using the R *lm* function, the module score was provided as the dependent variable and the following clinical and molecular variables were provided as predictors: age; comprehensive tumor purity estimate; pack-years smoked; mutation burden; AJCC T, N and M scores; tumor lung site; and sex. Regression diagnostics were performed including tests for heteroscedasticity and normality of residuals, revealing no notable deviations. This regression approach was performed for the module scores derived from all the mouse DEGs and the jackknifed genes. The aged vs. young mouse gene lists, jackknifing classifications, and multivariable regression results are available in **Extended Data Table 17**.

## Data availability statement

Mouse lung AT2 cell chromatin immunoprecipitation data was obtained from the Gene Expression Omnibus (GEO) under accession code GSE158205^41^. All relevant data generated in this study will be deposited into a public repository and accession codes will be provided. No restrictions on data availability apply.

## Code availability statement

All custom code and mathematical algorithms will be deposited into github.org prior to publication of this work.

---

**Extended Data Figure 1.**
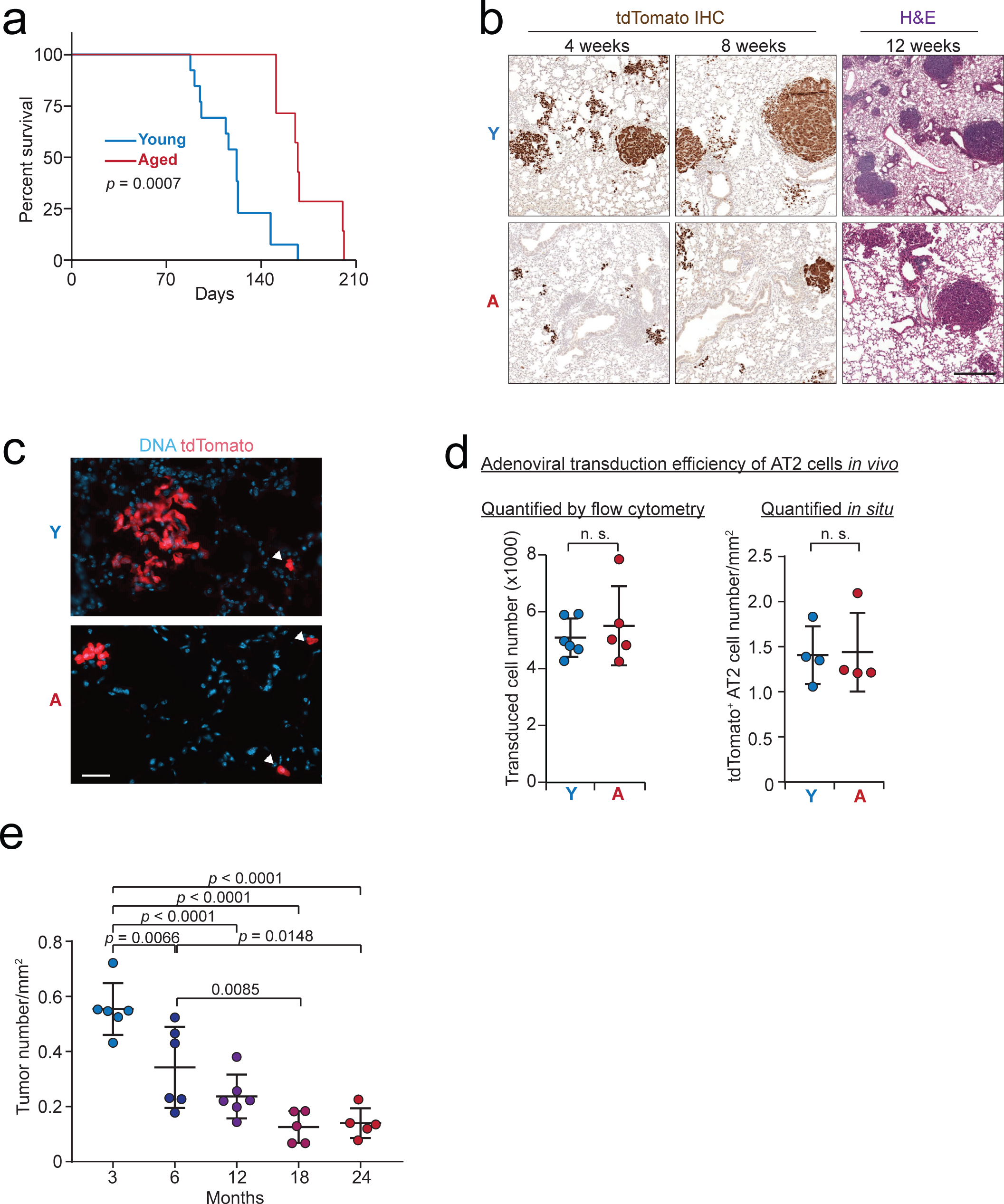
(**a**) Survival of aged vs. young *KP* LUAD mice post-tumor initiation (*n* = 13 young and 7 aged mice) (**Extended Data Table 1**). (**b**) Representative images of LUAD tumors in aged vs. young *KPT* (4 and 8 weeks) or *KP* (12 weeks) mice. At 4 and 8 weeks post-tumor initiation, LUAD cells were visualized by tdTomato immunohistochemistry. Scale bar: 200 µm. (**c**) Representative images of early LUAD lesions in aged vs. young *KPT* mice at two weeks post-tumor initiation. Transduced AT2 cells are visualized using a tdTomato fluorescence reporter allele. Arrowheads point to single-cell lesions (singletons). Scale bar: 50 µm. (**d**) Quantification of adeno-SPC-Cre transduction efficiency. Shown on the left is the total number of transduced AT2 cells, defined by the surface marker panel in **Extended Data Fig. 3b**, and quantified by flow cytometry. Successful transduction is defined by the switch from tdTomato to GFP fluorescence in aged and young *Rosa26^mTmG^* mice following intratracheal administration of adeno-SPC-Cre (*n* = 6 young and 5 aged mice). Shown on the right is the area density of tdTomato^+^ AT2 (SPC^+^) cells in tissue sections obtained from aged vs. young *Rosa26^LSL-tdTomato^* mice 3 days after intratracheal administration of adeno-SPC-Cre (*n* = 4 mice per group). (**e**) Quantification of tumor burden in 3, 6, 9, 12, 18, and 24 months-old *KP* mice at 12 weeks following tumor initiation with adeno-SPC-Cre (*n* = 5-6 mice per condition). Y: young, A: aged. *Log rank* test was used in (**a**). Student’s *t-*test was used in (**d**). One-way ANOVA was used in (**e**).

**Extended Data Figure 2.**
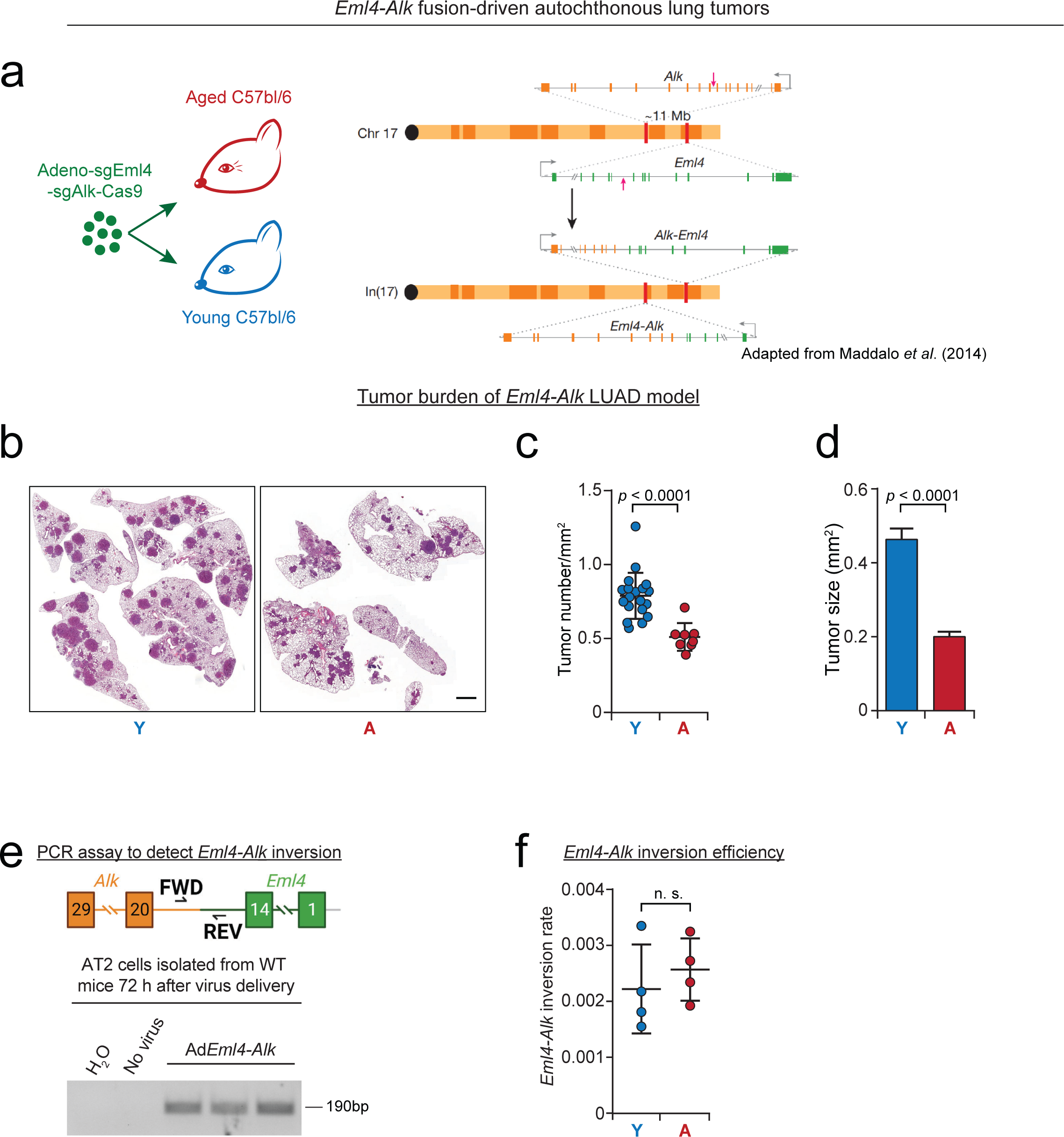
(**a**) Schematic summary of the *Eml4-Alk* LUAD model (image representing chromosomal rearrangement is reproduced from Maddalo et al.^27^). Lungs of aged (104-130 weeks old) and young (12-16 weeks old) wild-type C57BL/6 mice were intratracheally transduced with adeno-sgEml4-sgAlk-Cas9 virus, which induces an *Eml4-Alk* gene fusion via an intra-chromosomal inversion in chromosome 17. (**b-d**) Representative HE images (**b**) of pulmonary lobes showing the distribution of the tumors in the lungs, and the burden, quantified as tumor number per mm^2^ (**c**) (*n* = 19 young and 8 aged mice) and individual tumor size (mm^2^) (**d**) (*n* = 231 and 184 tumors from young and aged mice, respectively). Scale bar: 1 mm. (**e-f**) PCR assay used to detect *Eml4-Alk* inversion (**e**) and quantification of the inversion rate using the PCR assay in aged and young mice (**f**) (*n* = 4 mice per condition). Note that the PCR primers specifically amplify the fusion gene. Mean with SEM was shown in (**d**). Student’s *t* test was used in (**c-d**) and (**f**).

**Extended Data Figure 3.**
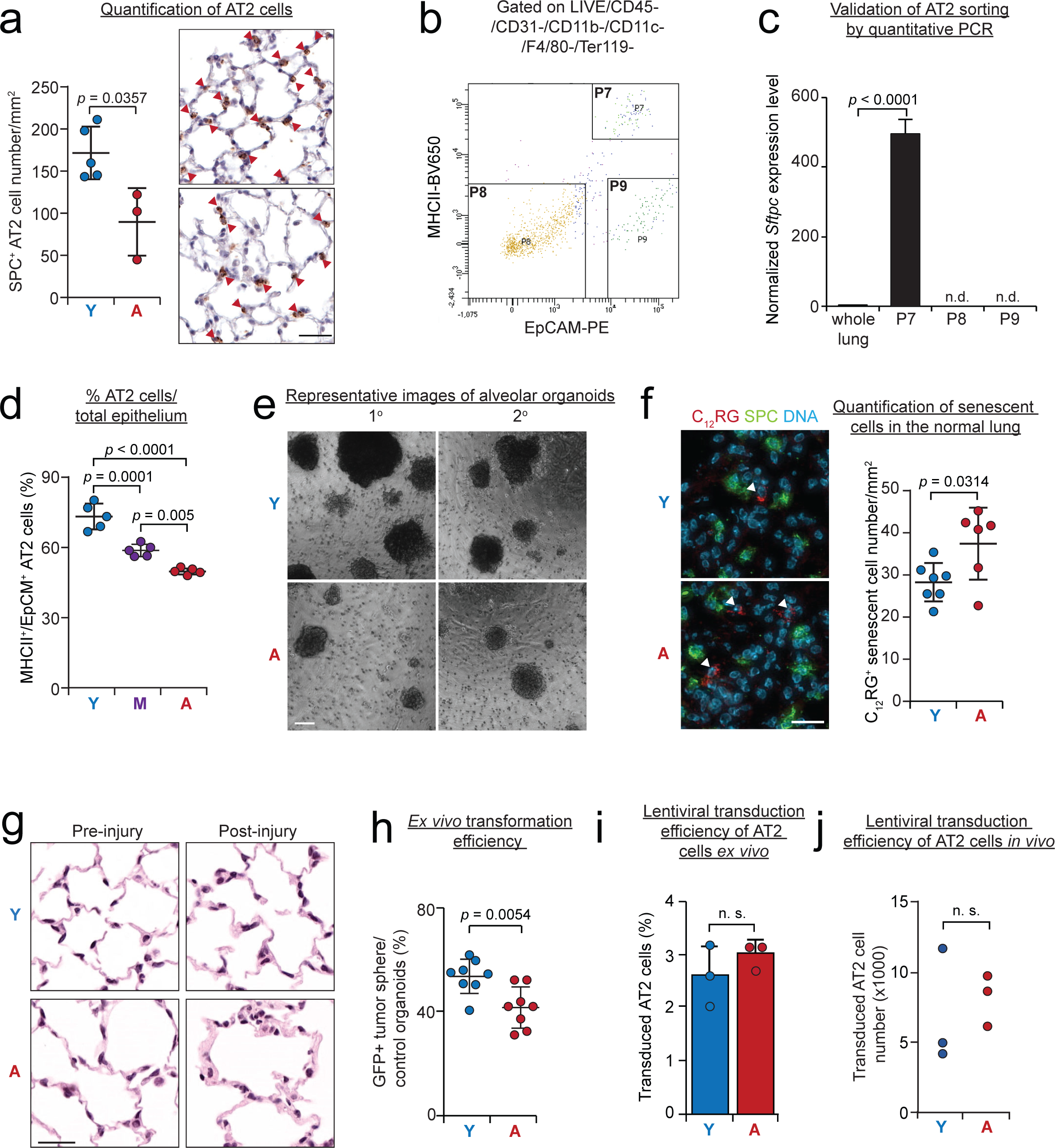
(**a**) Quantification of AT2 cells in aged and young mouse lungs. AT2 cells (red arrowheads) identified by surfactant protein-C (SPC) immunohistochemistry (*n* = 3 aged and 6 young mice). Scale bar; 50 µm. (**b**) FACS strategy for isolating mouse AT2 cells. AT2 cells are identified as DAPI^-^/Lineage (CD45, CD11b, CD11c, F4/80 and Ter119)^-^/EpCAM^+^/MHCII^+^ (P7). (**c**) Validation of sort purity by qPCR for the AT2 marker gene *Sftpc*. Note lack of detectable *Sftpc* expression in non-AT2 lineage-negative cells (P8 and P9). (**d**) Quantification of the percentage of AT2 cells in the total EpCAM^+^ epithelial cell pool in young (12-16 weeks old), middle-aged (52 weeks old), and aged (104-130 weeks old) wild-type C57BL/6 mice (*n* = 5 mice per condition). (**e**) Representative images of primary and secondary alveolar organoids. Scale bar: 100 µm. (**f**) Representative images (left) and quantification (right) of senescent cells identified by C_12_RG fluorescence-based detection of senescence-associated β-galactosidase activity. Note that C_12_RG positive senescent cells (arrowheads) do not express the AT2 cell marker SPC (*n* = 7 young and 6 aged mice). Scale bar: 50 µm. (**g**) Representative images of alveoli from young and aged animals pre- and post-hyperoxia injury (28 days). Scale bar: 50 µm. (**h**) Transformation efficiency of aged and young *KP-Cas9* AT2 cells calculated as the ratio of GFP^+^ transformed organoids/non-transformed alveolar organoids from the same animal (*n* = 8 biological replicates per condition). (**i**) Quantification of lentiviral transduction efficiency of aged and young AT2 cells in the *ex vivo* transformation assay. AT2 cells were isolated from both aged and young *KP* mice and transduced with lenti-PGK-GFP. Transduction efficiency was defined as the percentage of GFP^+^ AT2 cells over total number of AT2 cells (*n* = 3 biological replicates per condition). (**j**) Quantification of lentiviral transduction efficiency of aged and young AT2 cells *in vivo*. Aged and young *Rosa26^mTmG^* mice were transduced by Lenti-PGK-Cre and transduction efficiency was defined by the total number of GFP^+^ AT2 cells (GFP^+^/MHCII^+^/EpCAM^+^/lineage^-^/DAPI^-^), where Cre recombinase converts the tdTomato fluorescence to GFP fluorescence (*n* = 3 mice per condition). Y: young, A: aged. Mean with SD was shown in (**c**) and (**i**). *Mann-Whitney* test was used in (**a**), (**f**), (**i**) and (**j**). Student’s *t-*test was used in (**c**) and (**h**). One-way ANOVA was used in (**d**).

**Extended Data Figure 4.**
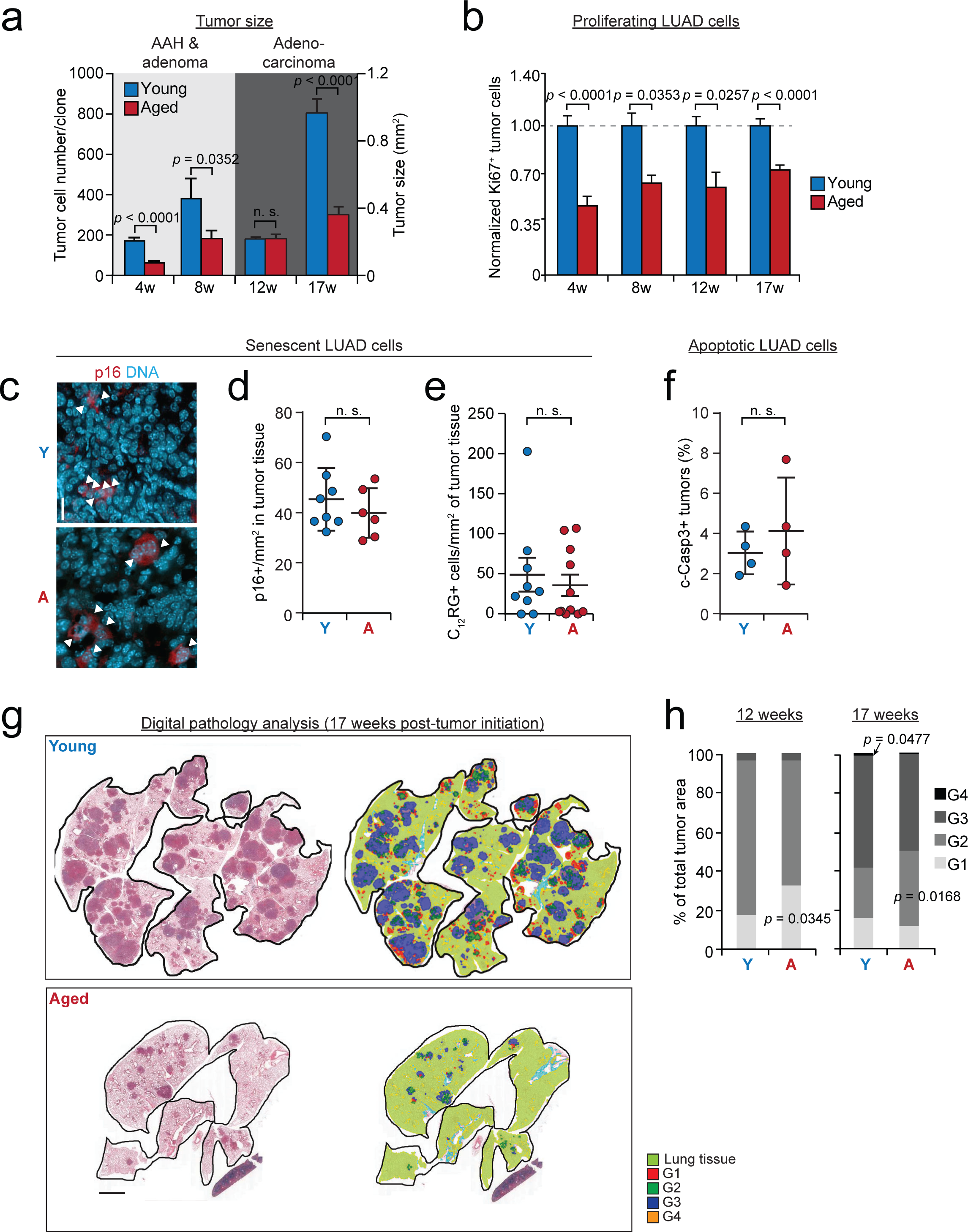
(**a**) Quantification of *KP* LUAD tumor size in aged vs. young mice at 4, 8, 12, and 17 weeks post-tumor initiation. Number of cancer cells per tumor nodule was quantified in *KPT* mice using tdTomato to visualize cancer cells at the 4- and 8-week time points; tumor size was quantified in *KP* mice from HE stained sections for at the 12- and 17-week time points. *N* = 109, 60, 371, and 358 tumors from young mice and *n* = 95, 66, 121, and 144 tumors from aged mice at 4, 8, 12, and 17 weeks. (**b**) Quantification of proliferating LUAD cells at different stages of tumor development. The proportion of Ki67 positive cells of total lung cancer cells (identified by endogenous reporter alleles or SPC immunofluorescence) was calculated and normalized to the average of young tumors at the corresponding time point. *N* = 55, 62, 127, and 306 tumors from young mice and *n* = 32, 62, 25, and 237 tumors from aged mice at 4, 8, 12, and 17 weeks. (**c-e**) Quantification of senescent cancer cells in young and aged *KP* LUAD tumors at 12 weeks post-tumor initiation. The senescent cells were identified by p16 (**c-d**) (*n* = 8 young and 6 aged mice) or C_12_RG staining (**e**) (*n* = 11 and 9 tumors from young and aged *KP* tumor-bearing mice, respectively). Representative images of p16 staining are shown in (**c**). Scale bar: 50 µm. (**f**) Quantification of cleaved caspase-3 (c-Casp3) positive LUAD tumors, defined as tumors with ≥ 1 c-Casp3^+^ cell (*n* = 4 mice per condition). (**g-h**) Histopathological grading of *KP* LUAD tumors in aged vs. young mice at 12 and 17 weeks post-tumor initiation. Representative images of the grading by an automated deep neural network (Aiforia Technologies) are shown (**g**) (*n* = 10 young and 5 aged mice for 12 week time point and *n* = 10 young and 10 aged mice for 17 week time point, respectively). The proportion of G1-G4 tumor area per total tumor area is shown in (**h**). Scale bar: 1 mm. Mean with SEM was shown in (**a-b**). Student’s *t* test was used in (**a-h**).

**Extended Data Figure 5.**
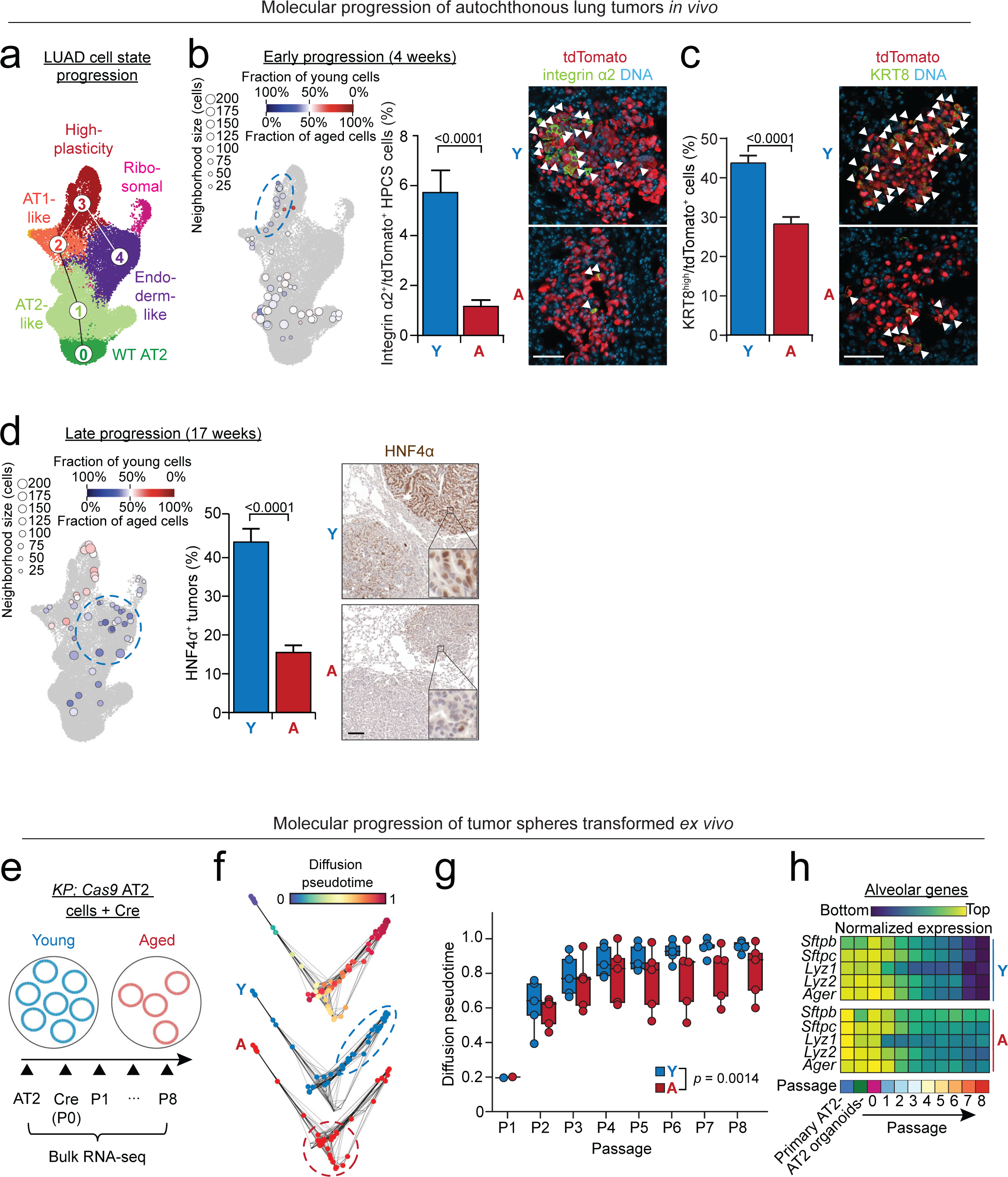
(**a**) Uniform manifold approximation projection (UMAP) embedding of LUAD single-cell transcriptomes isolated from young and aged *KP* tumors labeled based on previously defined transcriptionally distinct subsets^23^, at 4, 12, and 17 weeks post-tumor initiation. The numbers indicate order of cell state progression^23^. WT AT2: wild-type AT2 cells, the cell of origin (“0”). (**b**) MetaCell neighborhoods^72^, groups of similar cells representing discrete cell states, of 4-week-old *KP* LUAD tumors projected onto the UMAP introduced in (**a**). Circle size corresponds to the number of cells, while color signifies the fraction of young and aged cells forming a MetaCell neighborhood (blue: enriched in young; red: enriched in aged). Note enrichment of young cancer cells at the high-plasticity cell state (HPCS, dashed blue oval). Validation of higher proportion of young cancer cells in HPCS by immunofluorescence for the HPCS marker integrin α2 (green) at 4 weeks post-tumor initiation. tdTomato (red) marks cancer cells. *N* = 54 aged and 125 young tumors. Scale bar: 50 µm. Arrowheads indicate integrin α2 positive cells. (**c**) Quantification of KRT8-high tdTomato^+^ LUAD cells at 4 weeks post-tumor induction (*n* = 99 aged and 91 young tumors). Scale bar: 50 µm. Arrowheads point to KRT8-high cells. (**d**) MetaCell analysis of *KP* LUAD tumors at 17 weeks post-tumor initiation. Note enrichment of young cancer cells at the endoderm-like state (dashed blue circle), which is validated by immunohistochemical staining for the endoderm-like state marker HNF4α. Inset shows nuclear HNF4α immunolabeling in neoplastic cells (brown) in the tumors. The proportion of tumors containing >5% HNF4α+ cells per the total numbers of tumors is shown. *N* = 10 young and 11 aged tumor-bearing mice. Scale bar: 100 µm. (**e**) Schematic summary of experiment evaluating molecular progression of *KP* LUAD cells *in vitro*. Briefly, AT2 cells were isolated from aged vs. young *KP-Cas9* mice and transformed *ex vivo* by lentiviral Cre recombinase (P0, as shown in Fig. 1i). Bulk mRNA sequencing (RNA-seq) was performed over eight serial passages of tumor spheres (P1-P8). Non-*KP* alveolar organoids from both aged and young AT2 cells are also included. (**f**) Projection of bulk RNA-seq data from *ex vivo* transformed young and aged AT2 cells in diffusion pseudotime space. Upper panel: samples colored according to diffusion pseudotime using untransformed AT2 cells as the starting point (0). Lower panels: samples colored based on age (blue: young; aged: red). Note enrichment of young tumor spheres at later pseudotime points (blue dashed oval), whereas aged spheres are enriched at the midway point (dashed red circle). (**g**) Boxplots showing diffusion pseudotime distribution of young (blue) and aged (red) *ex vivo* transformed cells stratified according to passage (P). Individual samples are represented by data dots (*p* = 0.0014, *Wilcoxon* ranked sum test). *N* = 5 young and 5 aged biological *ex vivo* transformed tumor sphere cell lines. (**h**) Heatmap showing the mean expression of alveolar epithelial lineage markers [AT2 marker genes (S*ftpb, Sftpc, Lyz1* and *Lyz2*) and AT1 marker (*Ager*)]in young and aged *ex vivo* transformed cells stratified according to passage. The colormap shows the log_2_-fold change compared to sample mean for each gene harmonized for young and aged samples. Y: young, A: aged. Mean with SEM was shown in (**b-c**) and mean with SD was shown in (**d**). Student’s *t-*test was used in (**b-d**).

**Extended Data Figure 6.**
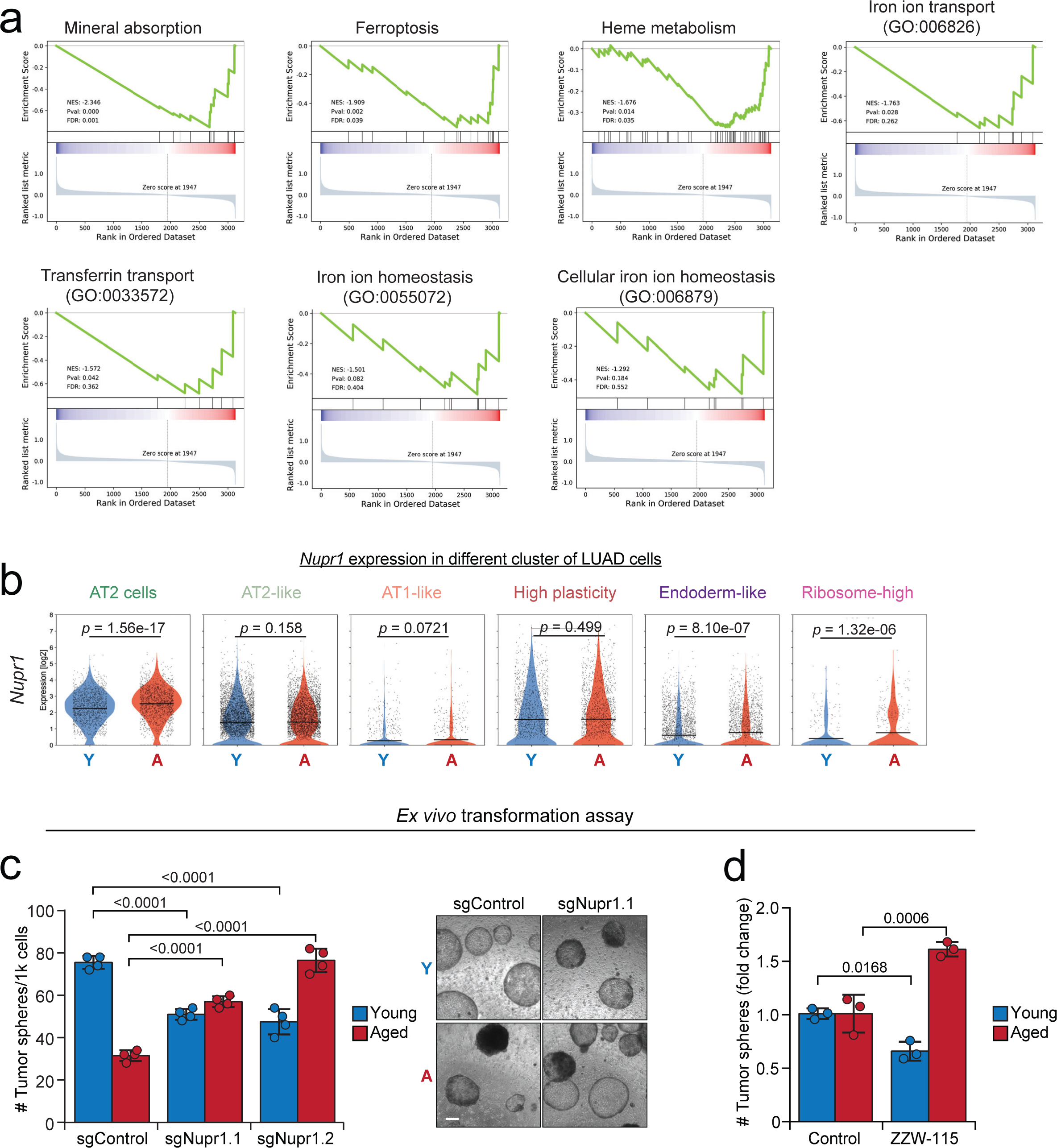
(**a**) Plots of gene set enrichment analysis (GSEA) showing the enrichment of gene sets linked to iron metabolism in age-related signatures introduced in Figure 2b and **Extended Data Table 3** ordered according to fold change. Red indicates enrichment in aged, blue enrichment in young. (**b**) Violin plots of showing the expression of *Nupr1* in young and aged AT2 cells and LUAD cell states. Rank-sum tests on single-cell gene expression were performed for statistical significance. (**c**) *Ex vivo* transformation of AT2 cells isolated from aged and young *KP-Cas9* mice with the indicated lentiviral vectors (Fig. 2f) delivering Cre + control sgRNA or two independent sgRNAs targeting *Nupr1* (*n* = 4 biological replicates per condition). Representative images of transformed tumor spheres are shown on the right. Scale bar: 100 µm. (**d**) *Ex vivo* transformation of AT2 cells isolated from aged and young *KP-RIK* mice, with or without treatment with the NUPR1 inhibitor ZZW-115 (2 µM) (*n* = 3 biological replicates per condition). Mean with SD was shown in (**c-d**). One-way ANOVA was used in (**c**-**d**).

**Extended Data Figure 7.**
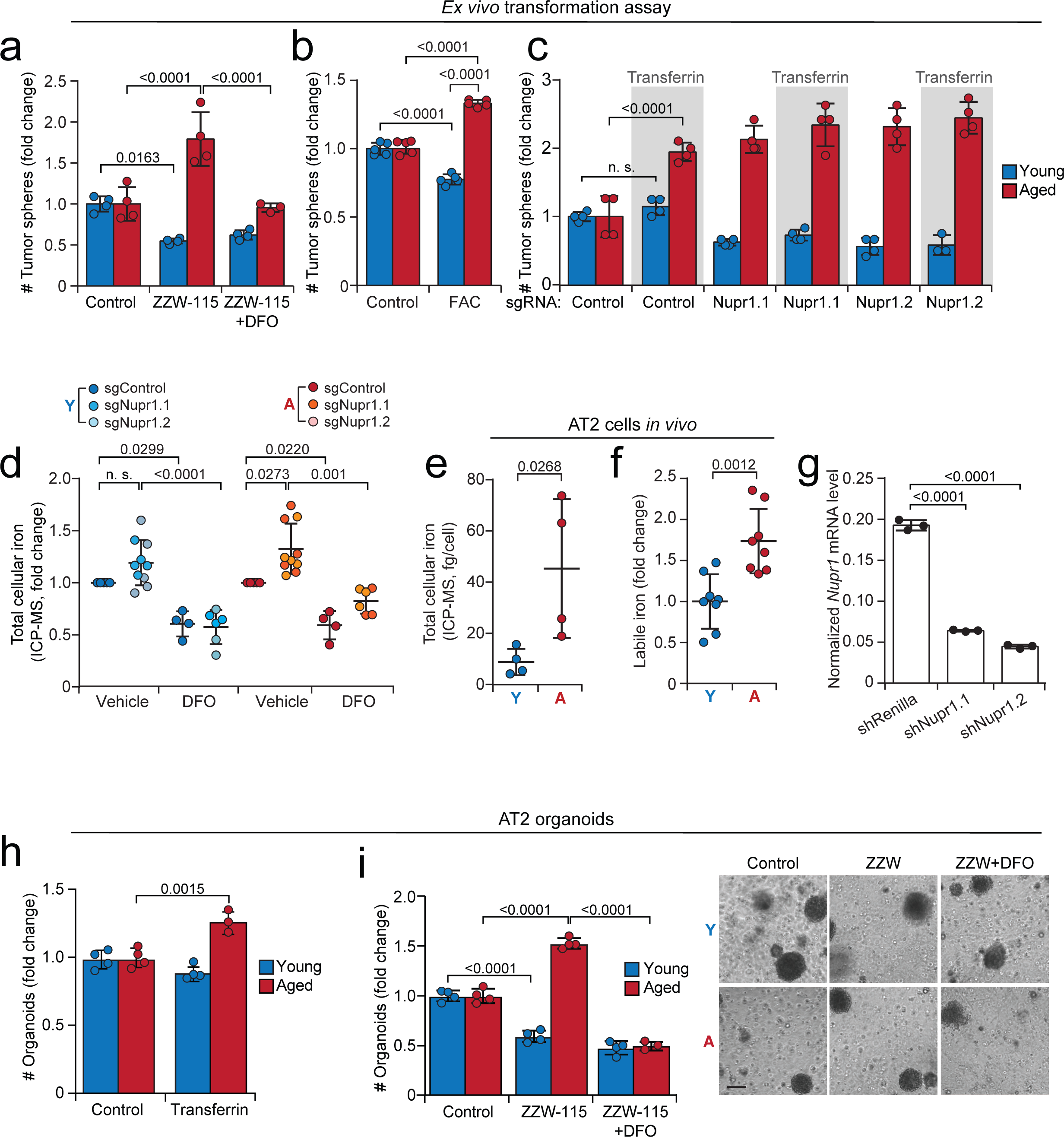
(**a**) *Ex vivo* transformation of AT2 cells isolated from aged and young *KP-RIK* mice by lentiviral PGK-Cre with or without treatment with ZZW-115 (2 µM) and deferoxamine (DFO, 2 µM) (*n* = 4 biological replicates per condition). (**b**) *Ex vivo* transformation of AT2 cells isolated from aged and young *KP-RIK* mice by lentiviral PGK-Cre with or without stimulation with 50 µM ferric ammonium citrate (FAC) (*n* = 5 biological replicates per condition). (**c**) *Ex vivo* transformation of AT2 cells isolated from aged and young *KP-Cas9* mice using lentiviral vectors delivering Cre recombinase and two sgRNAs targeting *Nupr1* or a control sgRNA with or without transferrin supplementation (50 µg/ml; *n* = 4 biological replicates per condition). (**d**) Normalized total cellular iron content measured by ICP-MS in LUAD cells derived from the *ex vivo* transformation experiment shown in Fig. 3c (*n* = 5, 10, 4, 6, 5, 10, 4, and 6 biological replicates per condition). (**e-f**) Measurement of total cellular iron in primary AT2 cells by ICP-MS (total iron, **c,** *n* = 4 biological replicates per condition) and staining of BioTracker Far-red dye (labile iron, **d,** *n* = 8 biological replicates per condition). (**g**) *Nupr1* mRNA in cultured *KP* LUAD cells transduced with lentiviral control shRNA (shRenilla) or shRNAs targeting *Nupr1* (*n* = 3 technical replicates per condition, a representative experiment that was repeated twice is shown). (**h**) Alveolar organoid formation by aged vs. young primary mouse AT2 cells stimulated with 50 µg/ml iron-loaded mouse transferrin or vehicle control (*n* = 3-4 biological replicates per condition). (**i**) Alveolar organoid formation by aged vs. young primary AT2 cells in the presence of vehicle control, ZZW-115 (2 µM), or ZZW-115 + 2 µM DFO (*n* = 3-4 biological replicates per condition). Representative images of alveolar organoids are shown on the right. Scale bar: 100 µm. Y: young, A: aged. Mean with SD was shown in (**a-c**) and (**g-i**). One-way ANOVA was used in (**a-d**), (**g-i**); *Mann-Whitney* test was used in (**e**) Student’s *t-*test was used in (**f**).

**Extended Data Figure 8.**
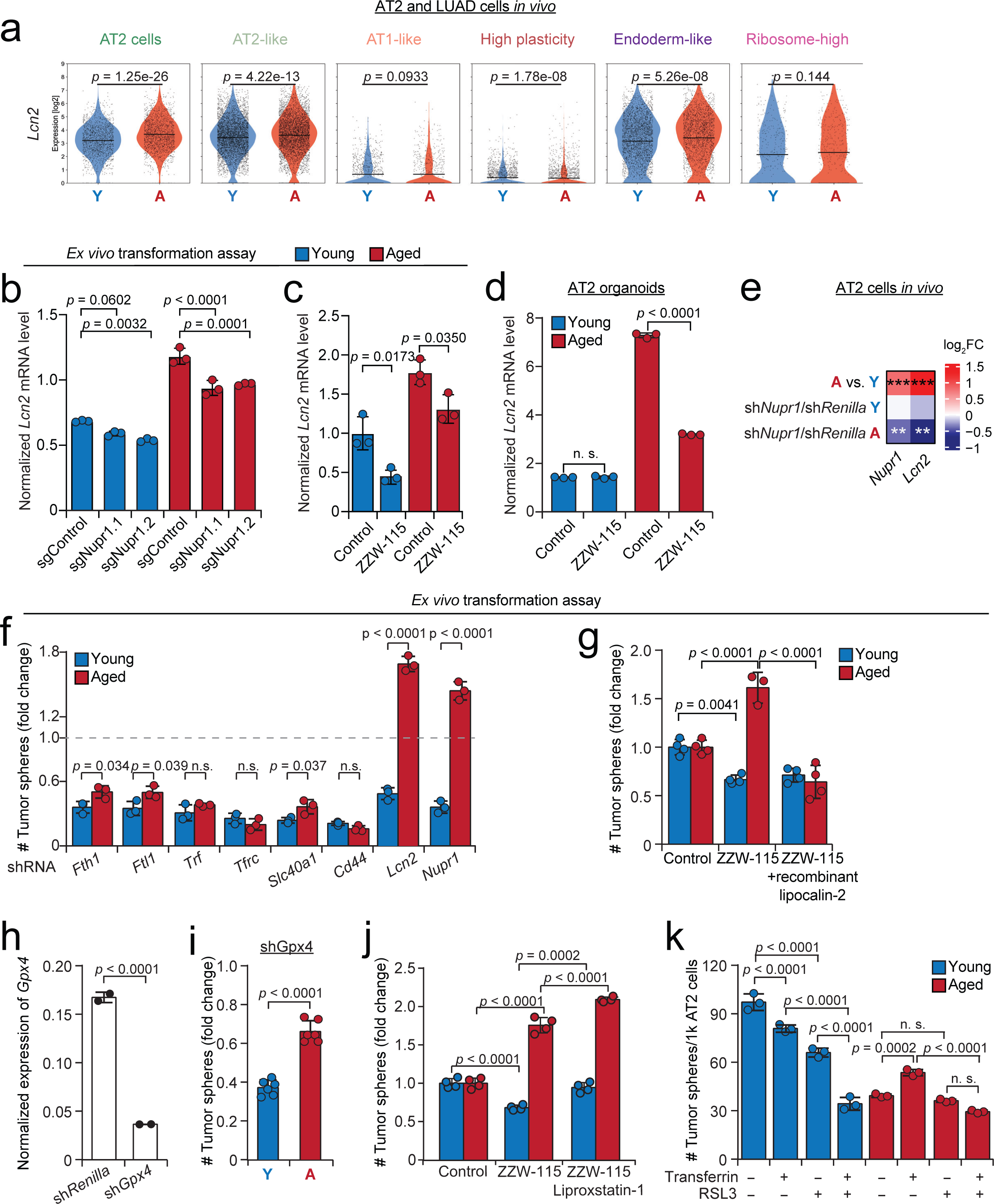
(**a**) Violin plots showing expression of *Lcn2* in young and aged AT2 cells and LUAD cell states. Rank-sum tests on single-cell gene expression vectors were performed for significance testing. (**b-d**) *Lcn2* mRNA level in transformed AT2 cells from the experiment in **Extended Data Fig. 6c** (**b**, *n* = 3 technical replicates), **Extended Data Fig. 6d** (**c**, *n* = 3 technical replicates), and normal alveolar organoids treated with or without NUPR1 inhibitor ZZW-115 (2 µM; **d**, *n* = 3 technical replicates). (**e**) Expression of *Nupr1* and *Lcn2* in AT2 cells expressing shRNAs targeting *Nupr1* or *Renilla* control *in vivo*. Gene expression was assessed by by single-cell mRNA sequencing of GFP^+^ AT2 cells isolated from the experiment in Fig. 3g. **, *p* < 0.01; ***, *p* < 0.001. Numerical *p* value was shown in **Extended Data Table 19**. (**f**) *Ex vivo* transformation of AT2 cells isolated from aged and young *KP* mice with lentiviral PGK-Cre + shRNAs targeting the indicated genes (*n* = 3 biological replicates per group). Data are shown as fold change compared to sh*Renilla* control. (**g**) *Ex vivo* transformation of AT2 cells isolated from aged vs. young *KP-RIK* mice using lentiviral PGK-Cre, with or without 100 ng/ml recombinant mouse lipocalin-2 and/or 2 µM ZZW-115 (*n* = 3-4 biological replicates per group). (**h**) Validation of shRNA targeting *Gpx4* by quantitative PCR (*n* = 2 technical replicates per condition). (**i**) *Ex vivo* transformation of aged and young AT2 cells of *KP* mice with shRNA targeting *Gpx4* (*n* = 6 biological replicates per condition). Data are shown as fold change compared to sh*Renilla* control. (**j**) *Ex vivo* transformation of AT2 cells isolated from aged vs. young *KP* mice with PGK-Cre, with or without NUPR1 inhibitor ZZW-115 (2 µM) and ferroptosis inhibitor liproxstatin-1 (5 µM). *N* = 4 biological replicates per condition. (**k**) *Ex vivo* transformation of AT2 cells isolated from aged vs. young *KP* mice using lentiviral PGK-Cre, with or without mouse transferrin or ferroptosis inducer RSL-3 (*n* = 3 biological replicates per condition). Mean with SD was shown in (**b-d**) and (**f-k**). One-way ANOVA was used in (**b-d**), (**g**), (**j**) and (**k**); Student’s *t* test was used in (**f**) and (**h-i**).

**Extended Data Figure 9.**
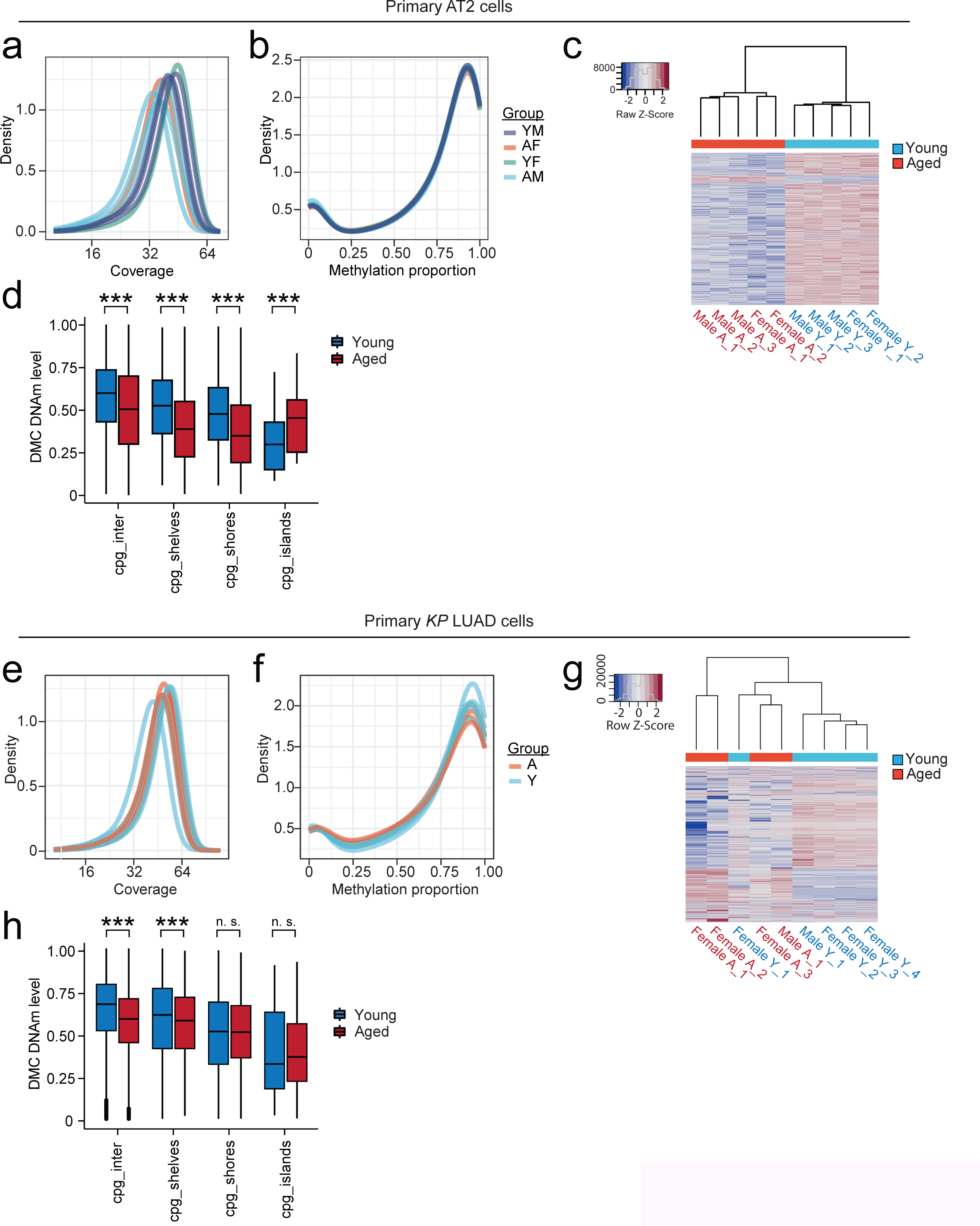
(**a**) Density plot showing Enzymatic Methyl-Sequencing (EM-Seq) coverage in primary AT2 cell samples. (**b**) Density plot of the methylation proportion in AT2 cell samples. (**c**) Heatmap showing unsupervised clustering of aged vs. young AT2 cells based on DNA methylation profiles. (**d**) Number of differentially methylated cytosines (DMCs) in aged vs. young AT2 cells based on location of CpG residues. (**e**) Density plot showing EM-Seq coverage in primary *KP* LUAD cell samples. (**f**) Density plot of the methylation proportion of *KP* LUAD samples. (**g**) Heatmap showing unsupervised clustering of aged vs. young *KP* LUAD cells based on DNA methylation profiles. (**h**) Number of differentially methylated cytosines (DMCs) in aged vs. young *KP* LUAD cells based on location of CpG residues.

**Extended Data Figure 10.**
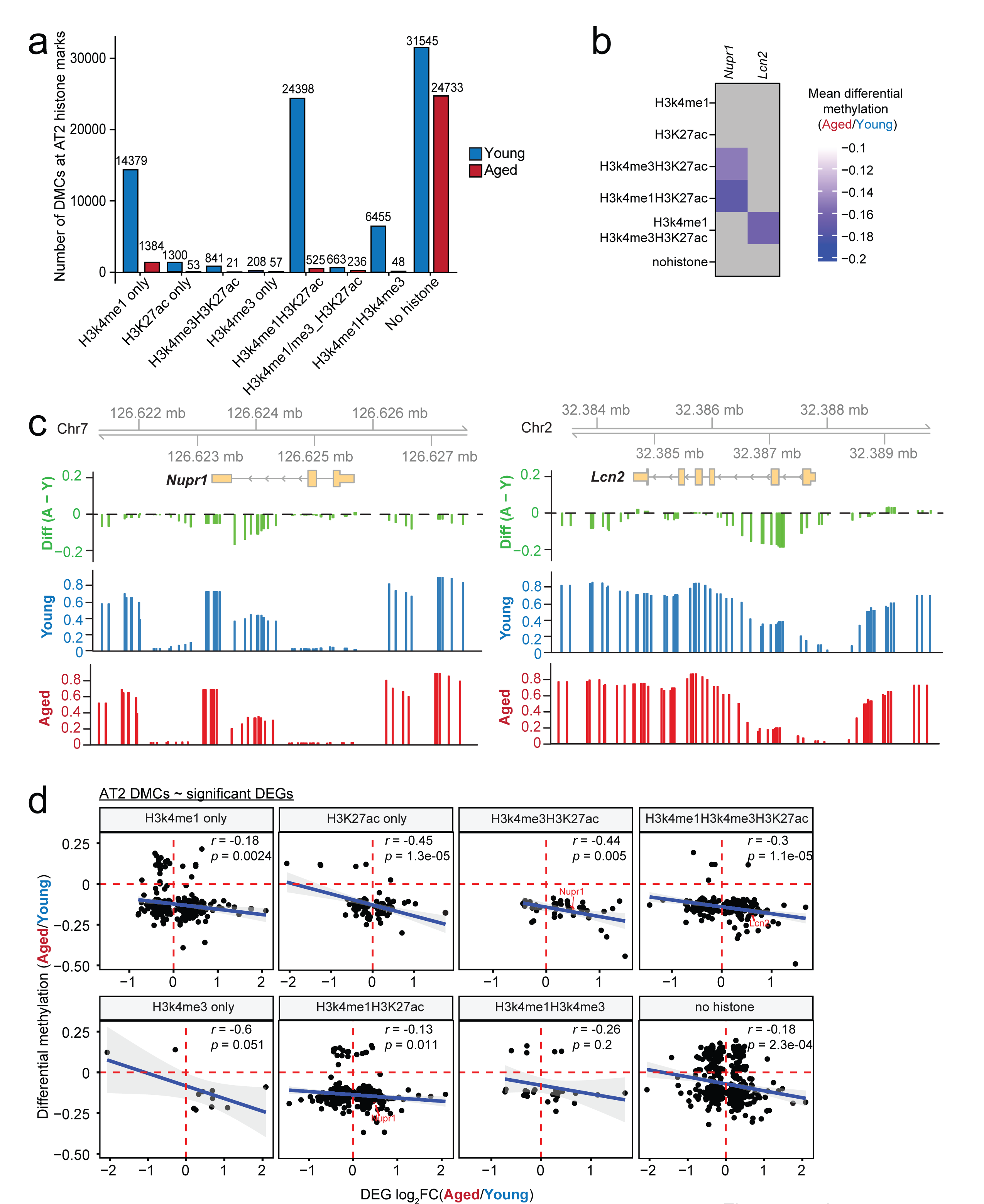
(**a**) The number of DMCs at sites marked by distinct histone modifications in AT2 cells. The height of the bars indicates higher methylation level in young (blue) or higher methylation in aged (red). The number of methylated CpGs detected is shown. (**b**) The mean differential methylation change of AT2 DMCs at histone marks at *Lcn2* and *Nupr1*. Grey means no DMC was detected. The color bar indicates the methylation change. Purple-blue color indicates demethylation in aged compared to young mice. Information of individual DMC was shown **Extended Data Table 9**. (**c**) Genome browser tracks of CpGs in the gene body of *Nupr1* and *Lcn2* in aged vs. young AT2 cells, measured by EM-Seq. Green indicates differential methylation between aged (red) and young (blue): green bars below zero indicate CpG hypomethylation in aged AT2 cells. (**d**) Pearson correlation between DMCs and DEGs in aged vs. young AT2 cells in each category of histone modification (FDR < 0.05). The regression line (blue) and its confidence interval (light grey) were plotted using a linear model.

**Extended Data Figure 11.**
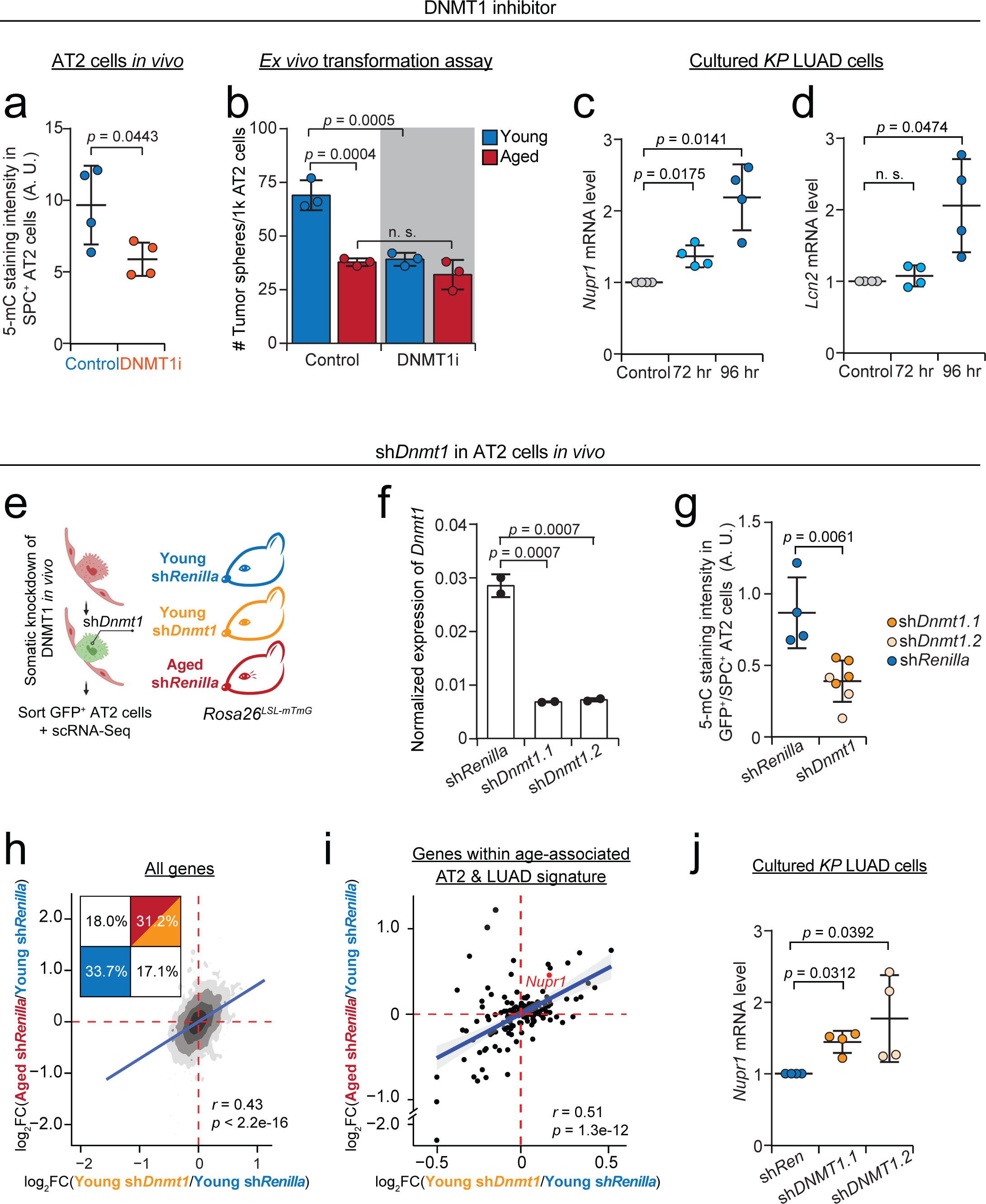
(**a**) Quantification of 5-methylcytosine (5-mC) immunofluorescence in lungs of young mice administered the DNMT1 inhibitor GSK3685032 for 8 days or vehicle controls. AT2 cells were identified as SPC^+^ cells. *N* = 4 mice per condition. (**b**) *Ex vivo* transformation of AT2 cells from aged and young *KP* mice with or without DNMT1 inhibition. *N* = 3 biological replicates. (**c-d**) Expression of *Nupr1* and *Lcn2* in *KP* LUAD cell lines with or without DNMT1 inhibition (*n* = 4 independent *KP* lines). The mRNA levels are normalized to the housekeeping gene *Gapdh* and further normalized to vehicle control. (**e**) Transcriptomic profiling of AT2 cells following shRNA-mediated knockdown of *Dnmt1* compared to *Renilla* control *in vivo*. (**f**) Validation of shRNA targeting *Dnmt1* by quantitative PCR (*n* = 2 technical replicates per condition). (**g**) Quantification of 5-mC immunofluorescence in AT2 cells of young *Rosa26^mTmG^*mice transduced with lentiviral PGK-Cre plus shRNAs targeting *Dnmt1* or *Renilla* control (*n* = 4 and 7 mice per condition). Given that successful delivery of the shRNAs is marked by switch of endogenous fluorescence from tdTomato to GFP, 5-mC in the GFP^+^ AT2 cells was quantified. (**h**) Contour plot showing correlation between the fold change (FC) of DEGs in AT2 cells isolated from young mice administered shDNMT1s or control shRNA (shRenilla) (*x*-axis) compared to aged vs. young AT2 cells expressing sh*Renilla* control (*y*-axis). Grayscale represents the probability distribution. Pearson correlation was calculated. The percentage of genes in each quadrant is significantly different from random (binomial test *p* < 2.2^-16^). The regression line (blue) was plotted using a linear model. (**i**) Scatter plot showing the correlation between the fold change of DEGs from the signature shared by AT2 cells and LUAD cells (Fig. 2b**; Extended Data Table 3**) in AT2 cells expressing sh*Dnmt1* or sh*Renilla* (*x*-axis) isolated from young mice vs. aged or young sh*Renilla*-treated mice (*y*-axis). Pearson correlation was calculated. The regression line (blue) and its confidence interval (light grey) were plotted using a linear model. (**j**) Expression of *Nupr1* in *KP* LUAD cell lines expressing shRNAs targeting *Dnmt1* or *Renilla* control (*n* = 4 independent *KP* lines). The mRNA level is normalized to the housekeeping gene *Gapdh* and further normalized to sh*Renilla*. Mean with SD was shown in (**b**) and (**f**). Student’s *t* test was used in (**a**), (**c**), (**d**), (**g**) and (**j**); One-way ANOVA was used in (**b-f**).

**Extended Data Figure 12.**
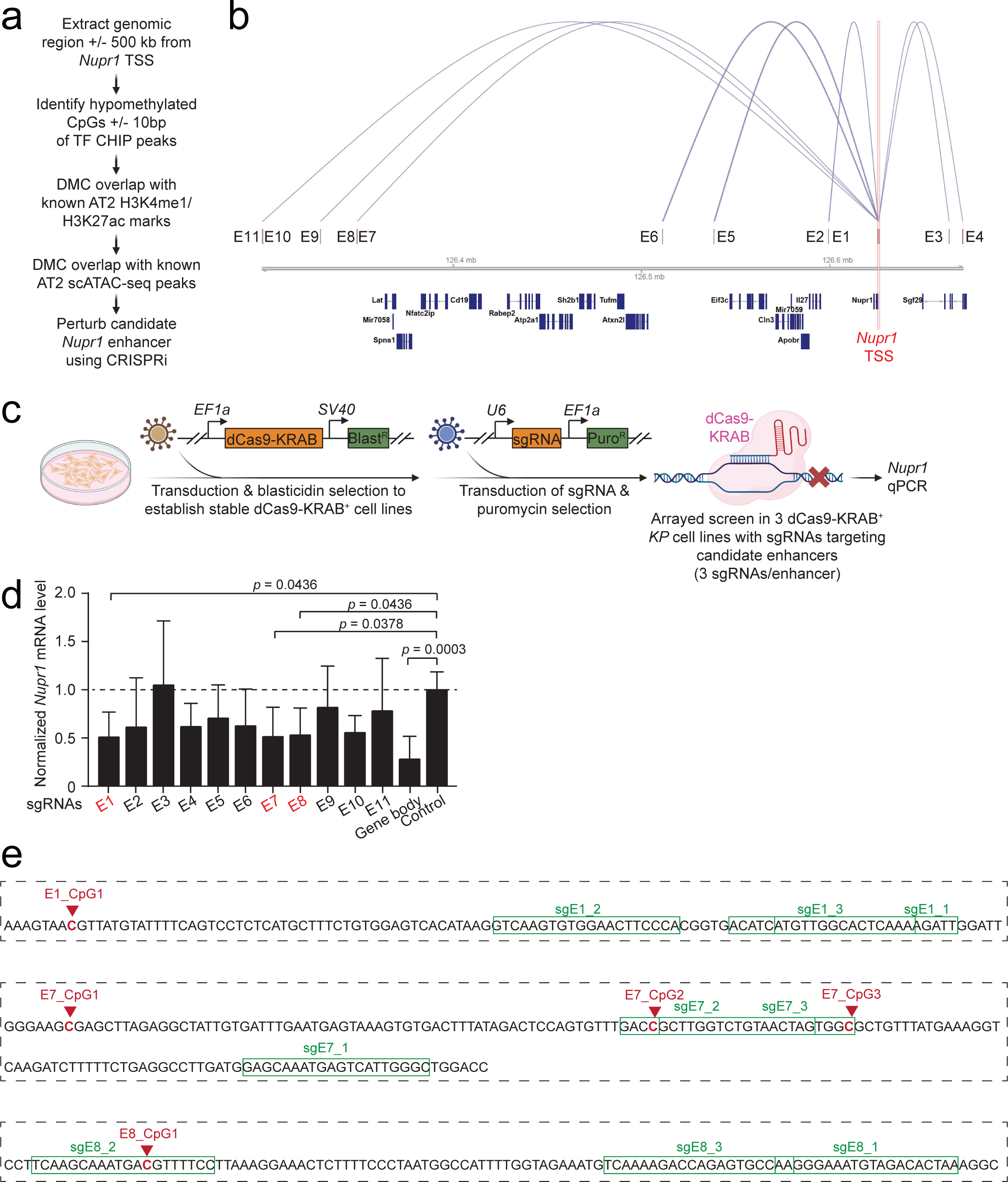
(**a**) Strategy of the selection of candidate distal *Nupr1*enhancers, including methylation status by EM-Seq, proximity to *Nupr1* transcription start site (TSS), overlap with transcription factor (TF) binding sites known to regulate *Nupr1*^38^, enhancer-associated histone markers^88^, and chromatin accessibility by single-cell ATAC-seq^41^. (**b**) Genomic distribution of the 11 candidate enhancers. (**c**) CRISPR interference (CRISPRi)-mediated functional testing of the candidate enhancers. (**d**) *Nupr1* quantitative PCR in *KP* LUAD cells expressing sgRNAs targeting the candidate enhancers (**Extended Data Table 10**, *n* = 9 data point from 3 biologically independent cell lines, each transduced with 3 different sgRNAs target the same enhancer). (**e**) Sequence context of differentially methylated CpGs in the high-confidence enhancers (E1, E7 and E8) and sgRNAs targeting these enhancers in CRISPRi assay. Mean with SD is shown in (**d**). *Kruskal-Wallis* test was used in (**d**).

**Extended Data Figure 13.**
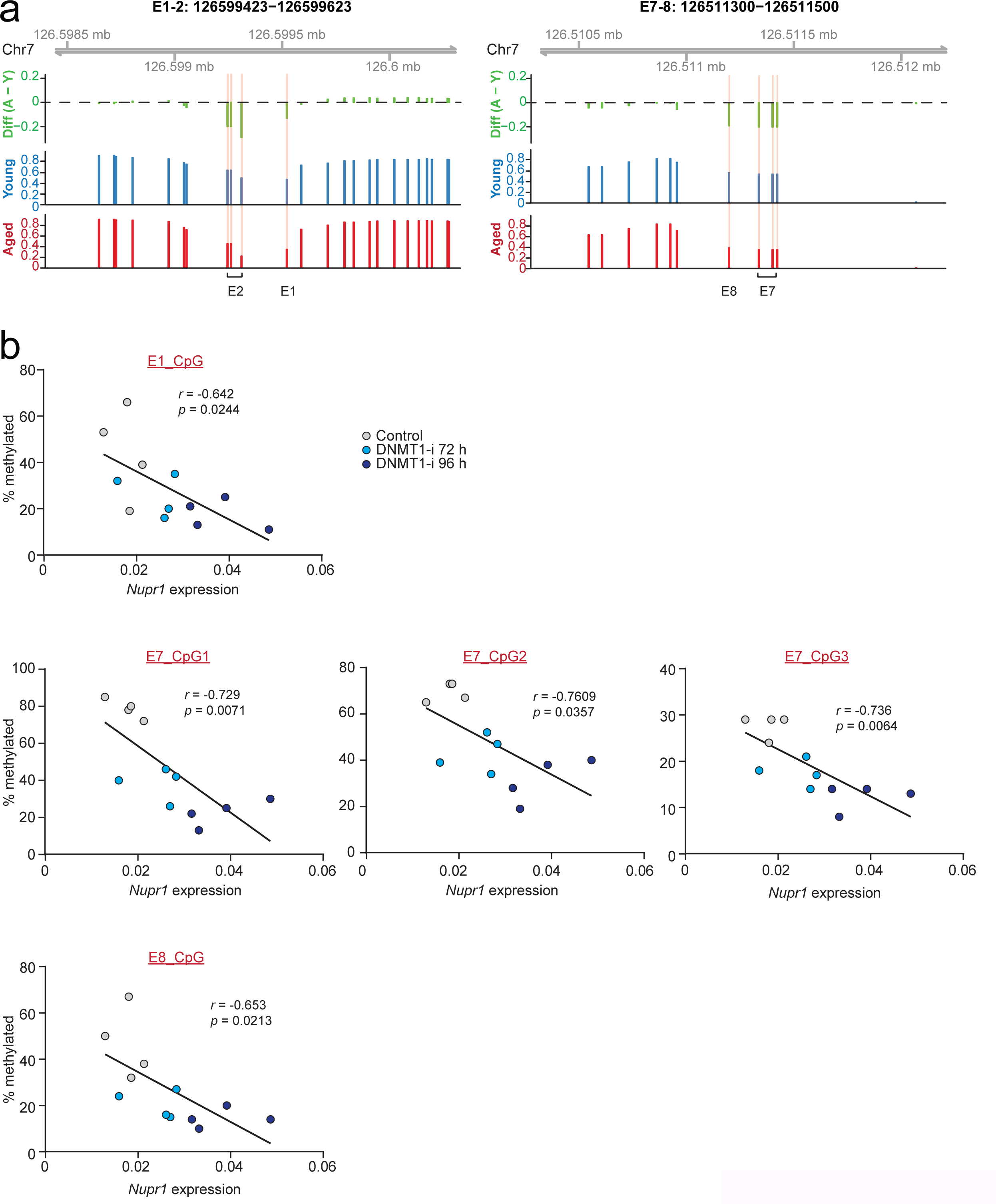
(**a**) Genome browser tracks showing differentially methylated CpGs at the high-confidence *Nupr1* enhancers (E1, E7 and E8) in aged vs. young AT2 cells, measured by EM-Seq. Green indicates differential methylation between aged (red) and young (blue): green bars below zero indicate CpG hypomethylation in aged AT2 cells. (**b**) Correlation between hypomethylation at the five CpGs at *Nupr1* enhancers hypomethylated with aging and *Nupr1* gene expression with or without DNMT1 inhibition. Pearson correlation coefficient (*r*) was calculated. The regression lines were calculated using a linear model.

**Extended Data Figure 14.**
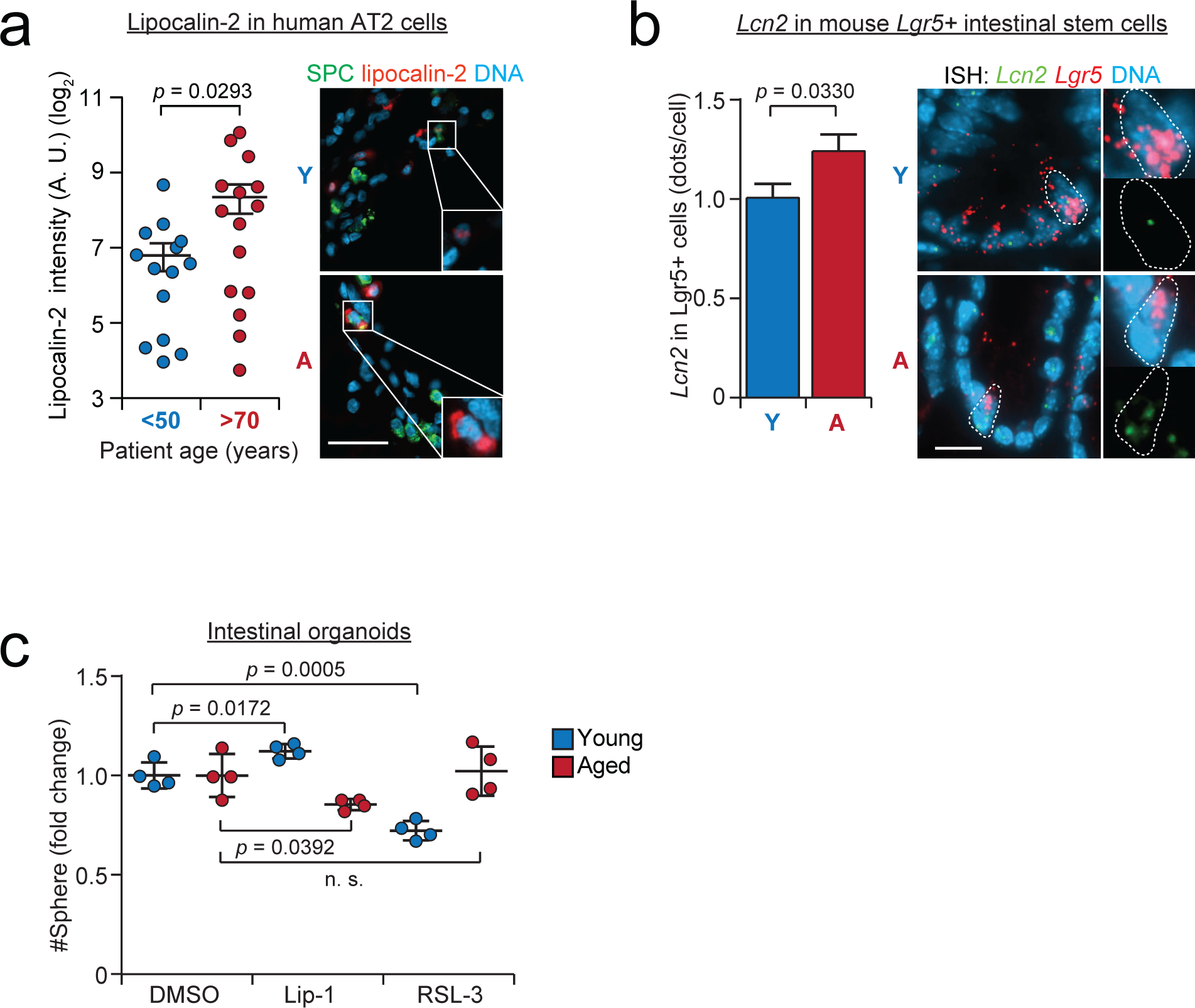
(**a**) Quantification of lipocalin-2 (*n* = 14 young/middle-aged cases vs. 15 aged cases) protein level in normal lung tissues of aged and young or middle-aged patients. The intensity of lipocalin-2 (red) immunofluorescence was quantified only in cells expressing the AT2 cell marker SPC (green). A. U.: arbitrary units. (**b**) Quantification of *Lcn2* mRNA (number of green dots) in aged vs. young mouse intestinal stem cells marked by the expression of *Lgr5* (*n* = 291 *Lgr5^+^* cells in the aged and *n* = 289 *Lgr5^+^* cells in the young). (**c**) Quantification of organoid formation by intestinal crypts isolated from aged vs. young mice, with or without liproxstatin-1 and RSL-3 (*n* = 4 technical repeats per condition, a representative experiment repeated 3 times is shown). Scale bar: 20 µm in (**a-b**). Y: young; A: aged. Mean with SEM was shown in (**b**). Student’s *t* test was used in (**a-c**).

**Extended Data Table 1:** Raw data from all laboratory experiments.

**Extended Data Table 2:** Differential gene expression analysis of aged vs. young AT2 cells transformed *ex vivo* controlled for passage or diffusion pseudotime (dpt).

**Extended Data Table 3:** Aging-associated differential gene expression shared between AT2 cells and LUAD (FDR < 0.05).

**Extended Data Table 4:** Gene set enrichment analysis using genes in the aged vs. young differential gene expression signatures shared between AT2 cells and LUAD.

**Extended Data Table 5:** Differential gene expression analysis of aged and young autochthonous *KP-Cas9* lung tumors in the context of *Nupr1* knockout or control sgRNA.

**Extended Data Table 6:** Differentially methylated cytosines in aged vs. young AT2 cells.

**Extended Data Table 7:** Differentially methylated cytosines in aged vs. young *KP* LUAD cells.

**Extended Data Table 8:** Aging-associated differentially methylated cytosines in AT2 cells at aging-associated differentially expressed genes annotated based on histone marks in AT2 cells.

**Extended Data Table 9**: Aging-associated differentially methylated cytosines in the proximity of transcription starting sites (TSS) of *Nupr1* and *Lcn2*, annotated based on histone marks in AT2 cells.

**Extended Data Table 10:** Differential gene expression following DNMT1 inhibition in young AT2 cells.

**Extended Data Table 11:** Differential gene expression results following sh*Dnmt1*expression in young AT2 cells.

**Extended Data Table 12:** Information of selected *Nupr1* enhancers.

**Extended Data Table 13:** Sequences of short hairpin RNAs (including knockdown efficiency), single guide RNAs, and PCR primers used in the study.

**Extended Data Table 14:** Details of pyrosequencing assay.

**Extended Data Table 15:** List of archived normal human lung resections.

**Extended Data Table 16:** List of archived human lung cancer sections and freshly resected normal human lung tissues.

**Extended Data Table 17:** Correlation of aged vs. young *KP* LUAD gene expression with human patients with KRAS mutant LUAD (TCGA data).

**Extended Data Table 18:** List of reagents.

**Extended Data Table 19:** Numerical *p* values that are not shown in the figures.

**Extended Data Table 20:** Conversion rate of methylated cytosines in the EM-Seq assay.

**Extended Data Table 21:** Veterinary pathology review of aged and young lung tissues 28 days after hyperoxia-induced lung injury.

